# Distinct mechanisms of CNV formation at the human 15q13.3 locus

**DOI:** 10.64898/2026.03.03.709017

**Authors:** Wolfram Höps, David Porubsky, DongAhn Yoo, Michelle de Groot, Amber den Ouden, Ronny Derks, Kendra Hoekzema, Kornelia Neveling, Alexander Hoischen, Helger G. Yntema, Human Pangenome Reference Consortium (HPRC), Pilar Caro, Alessandro De Falco, Bregje van Bon, Nicola Brunetti-Pierri, Christian P. Schaaf, Evan E. Eichler, Christian Gilissen

## Abstract

Human chromosome 15q13.3 is a hotspot for recurrent pathogenic copy number variants (CNVs) associated with schizophrenia, developmental delay and epilepsy. We generated long-read genome assemblies for 10 patient-parent trios and compared them to 581 population control haplotypes from diverse ancestry where this complex region is fully resolved. We find that both the long (“BP4-BP5”) and short (“*CHRNA7*”) forms of 15q13.3 CNVs arise predominantly by non-allelic homologous recombination (NAHR) whose susceptibility has been promoted by large inversion polymorphisms. While three BP4-BP5 breakpoints cluster in a 2 kbp PRDM9-enriched recombination hotspot, most CNVs are structurally distinct and affect the disease-relevant genes *ARHGAP11B* and *CHRFAM7A* differentially. Population analyses reveal at least 18 distinct structural haplotypes in 15q13.3 and show that the risk for BP4-BP5 CNVs is stratified more than 10-fold depending on ancestry, with 68.4% of Europeans but only 5.1% of East Asians predisposed, establishing inversion polymorphisms as major determinants of population-specific recurrent CNV susceptibility. Comparison to six ape species indicates that the duplication architecture promoting instability expanded recently and is largely human-specific.

## Introduction

The 15q11-q14 region of the human genome is highly variable and prone to copy number changes, such as deletions and duplications. These copy number variants (CNVs) occur recurrently in unrelated individuals between long (e.g., >10 kbp) and highly similar (>95%) repeats known as segmental duplications (SDs). This suggests meiotic non-allelic homologous recombination (NAHR) as the predominant mechanism of CNV formation ^1,2^. The 15q11-q14 region harbors five SD-rich breakpoint regions (BP1 to BP5), which predispose to various recurrent CNVs that give rise to different genomic disorders, such as Angelman and Prader-Willi syndromes ^3^ (Figure 1A). In the 15q13.3 region, CNVs of ∼1.5-2 Mbp occur between BP4 and BP5 (referred to as BP4-BP5 CNVs), as well as smaller deletions/duplications (∼350-680 kbp) between a repeat known as distal CHRNA7-LCR and BP5 (referred to as *CHRNA7* CNVs). These deletions and duplications in the 15q13.3 region give rise to variable neurodevelopmental phenotypes, often including intellectual disability (ID), autism, epilepsy, and schizophrenia ^1,4–8^, and other phenotypes such as hypotonia. The region flanked by BP4 and BP5 includes several protein-coding genes, notably *CHRFAM7A* (a *CHRNA7* and *FAM7A*/*ULK4* fusion), *ARHGAP11B* (Rho GTPase activating protein 11B), *FAN1* (Fanconi-associated nuclease 1), *KLF13* (Kruppel-like factor 13), *OTUD7A* (OTU deubiquitinase 7A), and *CHRNA7* (cholinergic receptor nicotinic alpha 7 subunit). The smallest critical region encompasses only *CHRNA7* ^9^. Whereas the *CHRNA7* deletions may occur as an inherited or a de novo event, all reported *CHRNA7* duplication events are inherited; usually from unaffected or mildly affected parents ^9^.

**Figure 1.**
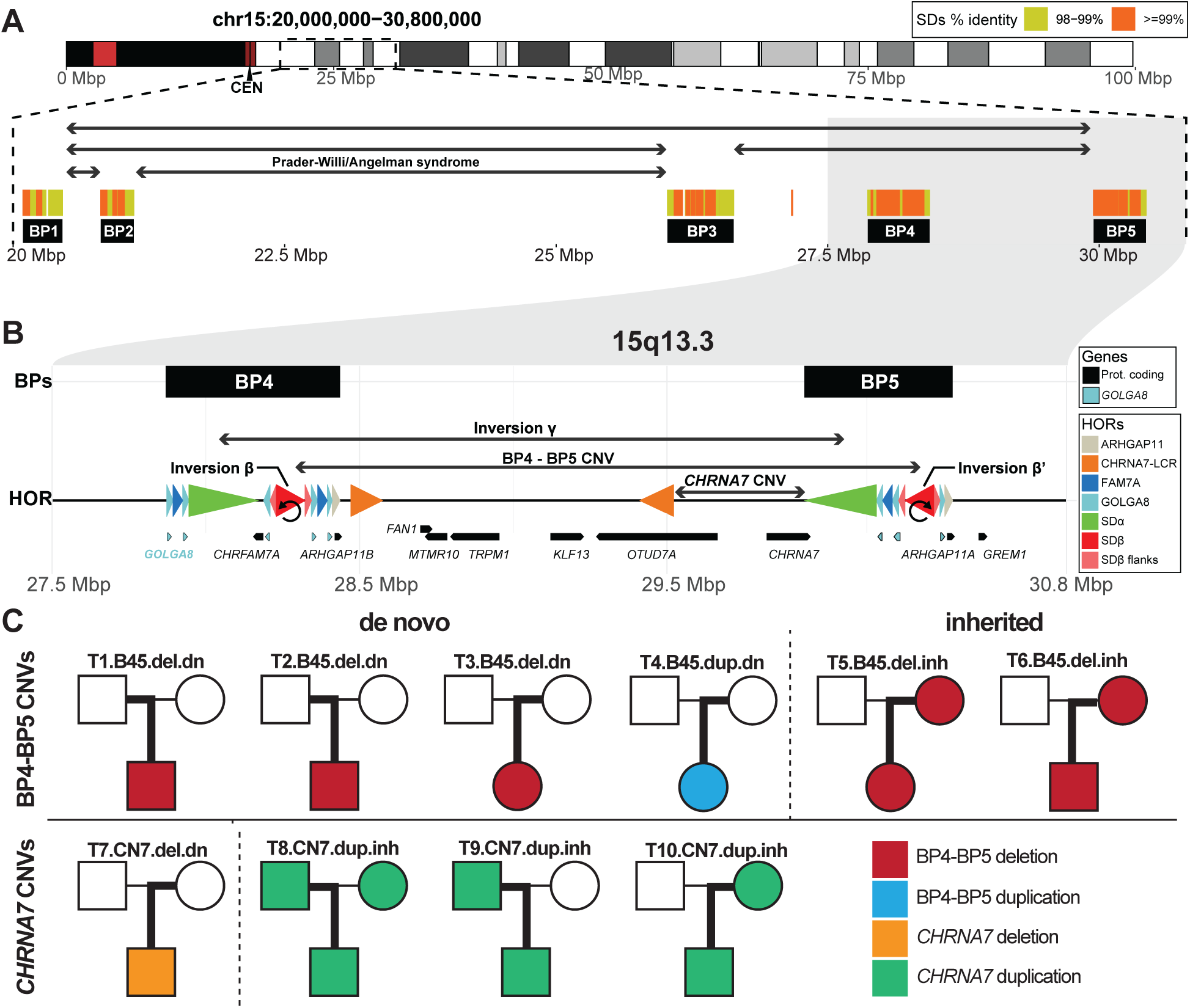
Overview of the chromosome 15q11.2-13.3 region and patient families. A) Overview of genomic hotspots at chromosome 15q associated with recurrent CNVs and inversions at the location of five breakpoints (BP1-5) with respect to segmental duplications and deletion/duplication syndromes including Prader-Willi/Angelman B) Zoomed-in view of the region of interest (15q13.3 microdeletion/microduplication syndrome) flanked by BP4-BP5. The higher order repeat (HOR) structure of this region in T2T-CHM13 is shown by direction pointing arrowheads colored by the HOR unit (see legend). We highlight inversion polymorphisms of SDβ and SDβ′ with circular arrows. We also mark approximate positions of known inversions and CNVs in the region. Below there is an annotation of protein-coding genes (black) and multiple copies of *GOLGA8* genes (cyan). C) Overview of studied patient-parent trios. Each trio carries an ID indicating trio number, CNV region, deletion/duplication, and de novo/inherited. The pedigrees are colored by carriership status of the different types of 15q13.3 CNVs (color legend: bottom right). Thick lines indicate the parent of origin of each offspring’s 15q13.3 CNV.

NAHR can occur between paralogous sequences on the same chromatid, sister chromatids, or on the homologous chromosome. In the latter two, deletions and duplications are reciprocal products ^10^. The mutational outcomes of NAHR depend on the relative orientation of the flanking SDs. If in opposite orientation with respect to each other, they predispose to inversions (in<u>tra</u>chromosomal NAHR), isodicentric chromosomes, and acentric palindromic fragments (in<u>ter</u>chromosomal NAHR). In contrast, directly oriented repeats mediate deletions, duplications (intra or interchromosomal), and acentric fragments (in<u>tra</u>chromosomal) ^11^. Previous studies have drawn two levels of connections between recurrent inversions and disease loci susceptible to CNVs, such as the 7q11.23 (Williams-Beuren syndrome), 22q11.2, 3q29, and 15q13.3 loci ^12,13^. First, recurrent inversions tend to affect the same regions as recurrent CNVs, as both are products of NAHR and are thus mediated by the same flanking SDs. Second, we and others have hypothesized that structural variations, particularly inversions, may predispose to meiotic CNV formation through NAHR by reorienting flanking SD sequences from opposite to direct orientations ^1,12^. Indeed, a survey of 28 diploid genome assemblies revealed 37 loci harboring complex SVs, which could be understood as the result of multiple, subsequent NAHR events, where one SV predisposes to the next ^14^.

Five different polymorphic inversions in the 15q13.3 region were previously described in the general population (inversions β, β′, γ, γ-del and δ in Figure 1B; Inv-β, Inv-γ, etc), and these, together with other CNVs of the breakpoint regions, are known to give rise to at least five structural haplotypes ^15^. However, direct evidence of an inversion-mediated CNV formation mechanism among patients carrying a 15q13.3 CNV has so far been lacking, primarily due to the inability of short-read sequencing technologies to fully assess the structure of flanking SDs and their relative orientation.

In this study we investigated the structure of chromosome 15q13.3 based on 581 fully assembled human haplotypes from diverse human ancestries and long-read sequencing on 10 patient-parent trios carrying CNVs in the 15q13.3 disease region, including five CNVs that arose de novo. Our goal was to test whether, as hypothesized, large-scale 15q13.3 CNVs in patients occur preferentially on parental alleles that carry a likely CNV-predisposing structural haplotype, to identify the underlying mutational mechanisms and investigate whether differences between individual CNVs may help explain phenotypic variability.

## Results

### Architecture of the 15q13.3 region

The chromosome 15q11.2-13.3 region (T2T-CHM13; chr15:20,000,000-30,800,000) harbors five blocks of SDs marking common inversions and CNV breakpoints (BP1-5) ^15^. Focusing on the 15q13.3 region (T2T-CHM13; chr15:27,500,000-30,800,000), we first defined a higher-order repeat (HOR) structure based on the T2T-CHM13 reference sequence (Methods, Supplementary Figure 1). This approach re-identifies the four largest SD pairs termed SDα, SDβ, CHRNA7-LCR, and *GOLGA8* elements as described previously ^15^. However, our finer HORs also reveal a more nuanced structure with opposite oriented flanking repeats of SDβ marked as ‘SDβ flanks’ along with a ‘FAM7A’ repeat, and a 26 kbp repeat encompassing *ARHGAP11A/B* genes, ‘ARHGAP11’ (Figure 1B).

### Assessment of ten chromosome 15q13.3 CNV patient trios Patients, sequencing and genome assembly

To study the formation of both groups of recurrent 15q13.3 CNVs (BP4-BP5 and *CHRNA7* CNVs), we selected 10 parent-child trios of index patients who had previously been diagnosed with 15q13.3 syndrome of varying severity and genetic testing had indicated a large-scale CNV in the 15q13.3 region. Among the ten trios, six (T1-T6) carry BP4-BP5 CNVs and four (T7-T10) carry *CHRNA7* CNVs, and both groups present a mix of deletions and duplications and de novo and inherited events (Figure 1C, Supplementary Data 1 and 2). All trios are of European ancestry (Supplementary Figure 2). We generated ∼30X coverage of long-read high-fidelity (HiFi) PacBio sequencing data for all patients and their parents, and ∼15X coverage ONT-UL for three selected individuals, and derived local de novo assemblies of the 15q13.3 region, which we validated extensively (Methods), including via orthogonal Optical Genome Mapping for 27% (8/30) of samples (Supplementary Figure 3, Supplementary Note 1). This approach resulted in end-to-end assemblies of the region in 20/20 haplotypes of the index patients and 34/40 haplotypes of the parents (Supplementary Data 3). Patient haplotypes were fully phased via pedigree-based phasing and parent haplotypes via custom parent-child duo phasing in the case of de novo events (Methods). Previously determined HOR sequences were mapped onto assemblies to guide haplotype analysis (Methods). Because of the different structure of BP4-BP5 and *CHRNA7* CNVs, we initially analysed the two groups separately.

### Inversions enable NAHR as the predominant mechanism for BP4-BP5 CNVs

We first investigated the 6/10 families carrying a BP4-BP5 CNV (Figure 1C): three de novo deletions (T1-T3), one de novo duplication (T4), and two inherited deletions (T5-T6). A summary of identified CNV structures and mechanisms is indicated in Figure 2A. Among four de novo CNVs, three (T2-T4) display an analogous sequence organization both in the CNV-carrying alleles and the parental allele on which they arose (Figure 2B). In all three trios, the offspring’s CNV flanks map to a parental allele carrying Inv-β (Figures 2A, 2B, and Supplementary Figure 4), which brings SDβ and SDβ′ into direct orientation. This architecture enables deletion or duplication of the intermediary BP4-BP5 sequence via NAHR (Supplementary Figure 5). Both flanks of the deletion in T2 map to the same paternal haplotype, indicating an intrachromosomal or interchromatidal event, and the duplication in T4 arose interchromatidally, as the duplicated segment carries no third haplotype (Supplementary Figure 6). In all three cases, the CNVs delete or duplicate the genes *ARHGAP11B*, *FAN1*, *MTMR10*, *TRPM1*, *KLF13*, *OTUD7A*, and *CHRNA7*, concordant with the majority of reported BP4-BP5 CNVs ^9^ (Figure 1B). The offspring’s de novo breakpoints all map to their respective parental SDβ/SDβ′ units and exhibit stretches of uninterrupted sequence homology between the donating and receiving ends (640, 82 and 153 bp for T2, T3, T4, respectively; Supplementary Figure 7, Supplementary Data 4), consistent with NAHR as the underlying mechanism.

**Figure 2.**
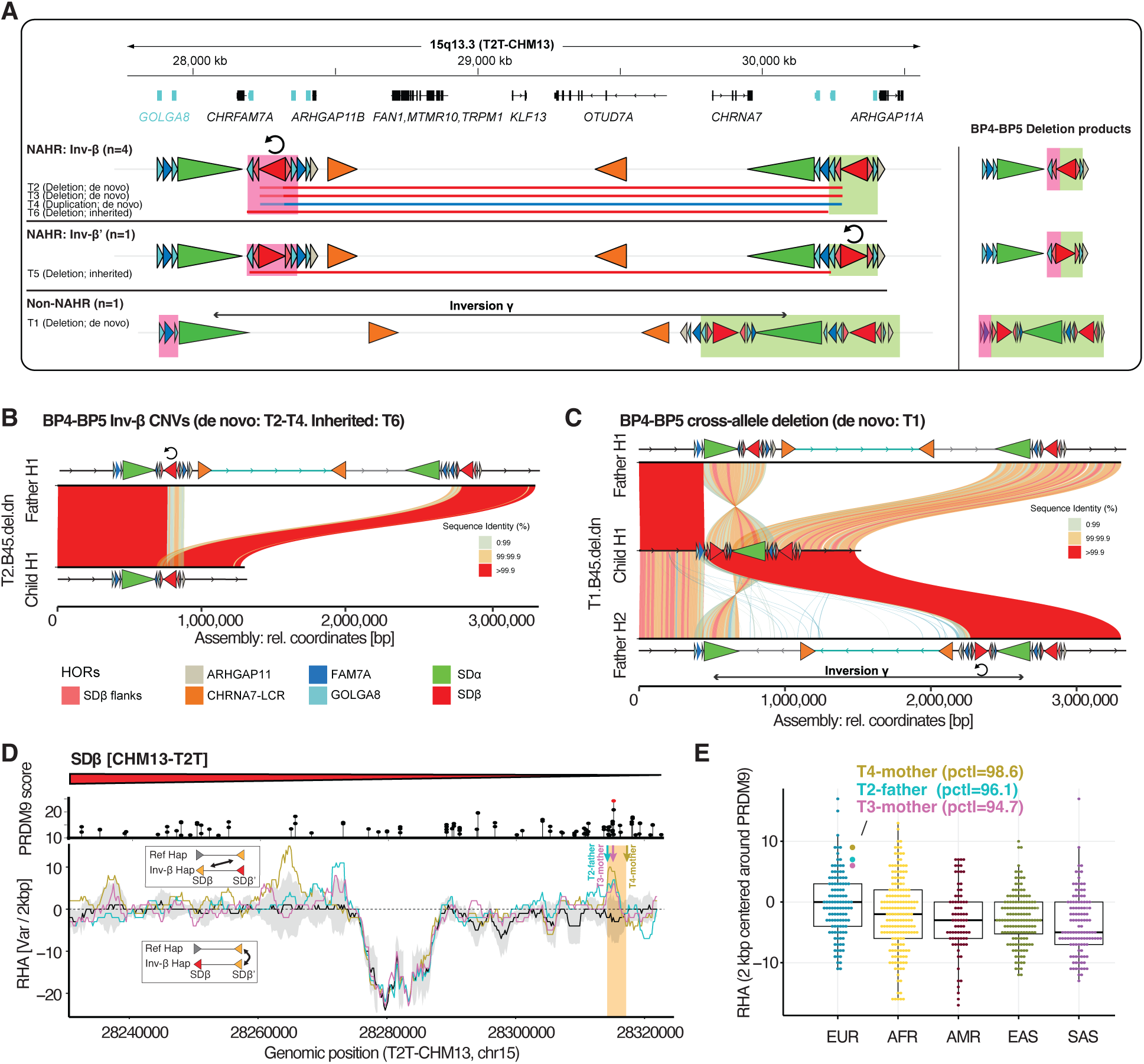
Structure and formation mechanisms of BP4-BP5 CNVs. A) Overview of the parental configurations, breakpoints, and products of observed BP4-BP5 CNVs. Coordinates and gene models are indicated on top, with all *GOLGA8* copies colored light blue. The three different formation mechanisms are indicated by HOR schematics indicating the key parental sequence configurations preceding CNV formation. Schematics of the derived CNV products per group are indicated on the right. The identity of pre- and post-breakpoint homologous regions are indicated by red and green boxes, respectively, and the same colors are indicated on the CNV alleles. B) SVbyEye plot showing the structure of the de novo BP4-BP5 CNV in T2 in relation to the parental haplotype on which it arose. HORs (colored arrowheads) are overlaid over all assemblies. Trios T2-T4 and T6 display analogous CNV structures (Supplementary Figure 4). C) SVbyEye plot showing the de novo CNV observed in T1, with again the index allele aligned to the parental alleles. The CNV is a rearrangement involving both parental homologous chromosomes. D) Comparison of sequence similarity between allelic and non-allelic copies of SDβ/SDβ′, in controls and in the parents of de novo CNVs (T2–T4). For each 2 kbp sliding window, we counted the SNVs separating the allelic repeat pair and the SNVs separating the non-allelic pair; the relative homology advantage (RHA) is the former minus the latter (Methods). Because fewer SNVs means greater similarity, a positive RHA indicates that the non-allelic copies are more similar to each other than the allelic copies are, possibly favoring NAHR (see insets). Colored lines show the three parental pairings (T2–T4); the black line and grey band show the median and 2.5th–97.5th percentile range across 612 RHA scores from unrelated samples (Methods). PRDM9 binding-site scores are plotted along the top (Methods). Colored arrows mark the three de novo breakpoints, and the yellow band marks their shared breakpoint region. E) Distribution of RHA scores in the 2 kbp window at the PRDM9 motif (highlighted in D), for control samples meeting the inclusion criteria (Methods) and for the three parental pairings from which de novo CNVs on SDβ/SDβ′ arose (T2-T4). Each point is one RHA score. Box plots show the median (center line), 25th-75th percentiles (bounds) and whiskers at 1.5× the interquartile range. Background: 612 RHA scores from 153 independent control samples (Methods; 108 European). Parental scores (T2–T4) are labelled with their percentile rank in the all-ancestry background.

The inherited BP4-BP5 deletion in T6 displays a CNV structure identical to that observed in T2 and T3. Comparison with 581 assembled population haplotypes ^16–18^ indicates that the CNV-carrying allele most closely matches a haplotype harboring an Inv-β configuration and suggests a NAHR-driven breakpoint (440 bp homology) located within the FAM7A HOR that forms part of the extended direct-facing duplication block generated by Inv-β (Figures 2A, Supplementary Figures 4, 8 and 9). The breakpoint region partly overlaps the first exon of *CHRFAM7A,* whose upstream sequence context gets exchanged as a result, though with untested effects on gene function (Supplementary Figure 9). The inherited BP4-BP5 deletion in T5 differs in the orientation of the remaining SDβ (Figure 2A) and best matches a population haplotype carrying an Inv-β′ configuration (Supplementary Figure 8). The breakpoints localize to the actively transcribed region of *GOLGA8* repeats resulting in a *GOLGA8* fusion (Supplementary Figures 8 and 9), again with breakpoint homology (1083 bp) suggesting NAHR as the underlying mechanism. T5 demonstrates that the Inv-β′ configuration, analogous to Inv-β, generates a recombination substrate for NAHR at BP4-BP5.

For T1, the index has undergone a mechanistically different CNV, which affects the same genes, yet yields a different SD structure. The transmitting parent carries one haplotype with an Inv-β and one haplotype with both an Inv-β and Inv-γ. One of the offspring’s CNV breakpoints maps to a *GOLGA8* element located on a different paternal haplotype (Figure 2C). The breakpoints display only 3 bp of microhomology suggesting a mechanism distinct from NAHR.

### Inversion β - mediated BP4-BP5 breakpoints cluster in a recombination hotspot

We further investigated the three de novo BP4-BP5 CNVs that occurred on the Inv-β allele in T2-T4. On SDβ, the three breakpoints cluster within just 2.5 kbp of each other (T2T-CHM13: chr15:28,314,880-28,317,299) and within 0, 325, and 2,078 bp distance to the single highest-scoring PRDM9 motif on SDβ (Figure 2D and Supplementary Figure 10, Methods), a degree of clustering that is unlikely to occur by chance (p = 0.0026, one-sided permutation test). These observations indicate that this region is likely a hotspot for recombination.

We considered that particularly high sequence similarity in T2-T4 between SDβ and SDβ′ could have promoted recurrent NAHR but found no evidence for this (Supplementary Note 2). We then hypothesized that recombination may involve competition between the correct allelic pairing on the homologous chromosome (SDβ′-SDβ′) and the CNV-inducing, non-allelic alternatives (SDβ-SDβ′) with the same or the homologous chromosome. NAHR might be especially likely if any non-allelic pairing offers a better local match than the allelic one (Supplementary Figure 11). To quantify this effect, we defined a relative homology advantage (RHA) score that measures relative sequence similarity between the allelic and non-allelic pairings based on variant counts (Methods). Positive RHA values indicate that a non-allelic pairing shows greater local homology than its allelic competitor. To assess whether the CNV-producing pairings in T2-T4 showed a higher RHA than expected by chance, we compared them against a background of 612 RHA scores from 153 control samples (HPRC and HGSVC; Methods). The three parental scores were significantly elevated in the 2 kbp region around the high-scoring PRDM9 motif (p = 5.2 × 10⁻⁵, two-sided Fisher’s exact test; ranked in the top 3.9%, 5.3% and 1.4% of all 612 background values). This enrichment remained significant against ancestry-matched European controls (n = 108 RHA values; p = 7.97 × 10⁻³, two-sided Fisher’s exact test; ranks 6.7%, 9.2% 2.5%) (Figure 2D, E, Supplementary Figures 12, 13).

In total, five of six BP4-BP5 CNVs in our dataset (83%, T2-T6) are best explained by NAHR on an Inv-β or Inv-β′ background, with three events facilitated by breakpoint clustering within a PRDM9-associated recombination hotspot and a single-nucleotide polymorphism (SNP) configuration that favors non-allelic over allelic recombination. A single CNV arose through replication-based mechanisms or alternative recombination pathways, suggesting that inversion-facilitated NAHR is the dominant, but not exclusive, mechanism for BP4-BP5 CNV formation.

### CHRNA7 CNVs mediated by inverted *CHRNA7*-LCR repeats

We next analyzed the 4/10 trios with the smaller recurrent *CHRNA7* CNV (Figure 1C). Our dataset entails one trio with a de novo *CHRNA7* deletion (T7) and three with duplications passed from parent to offspring (T8-T10). In T8, both parents carry a *CHRNA7* duplication, with the index inheriting only the paternal copy. The de novo *CHRNA7* deletion allele (T7) lacks ∼751 kbp of unique sequence between *OTUD7A* (exons 1-3) and *CHRNA7* and a copy of SDα but also generates an inverted duplication of a 149 kbp segment containing *ARHGAP11B* (Figure 3A). Parental assemblies and breakpoint mapping reveal that the event arose by crossover between CHRNA7-LCR and CHRNA7-LCR′ segments located on reference and Inv-δ configurations, respectively, creating an imbalanced fusion between the two alleles (Figure 3A). The breakpoint region harbors 43 bp of perfect homology (Supplementary Figure 7), or 557 bp with only one single-nucleotide variant (SNV), again supporting NAHR as the likely mechanism. The four resolved *CHRNA7* duplications (T8 (two distinct *CHRNA7* Dup alleles), T9, T10) display a similar but reciprocal outcome (Figure 3B and Supplementary Figure 14), with deleted and duplicated regions switched, and this time affecting the copy number of SDβ, rather than SDα (duplication: ∼426 kbp; deletion: ∼301 kbp). T8-maternal, T9, and T10 additionally carry an inversion of the intermediary sequence between CHRNA7-LCR and CHRNA7-LCR′ (Supplementary Figure 14). Although not observed de novo, these structures are consistent with a similar NAHR process as in T7 but mediated by the Inv-γ haplotype rather than the Inv-δ (Figure 3C).

**Figure 3.**
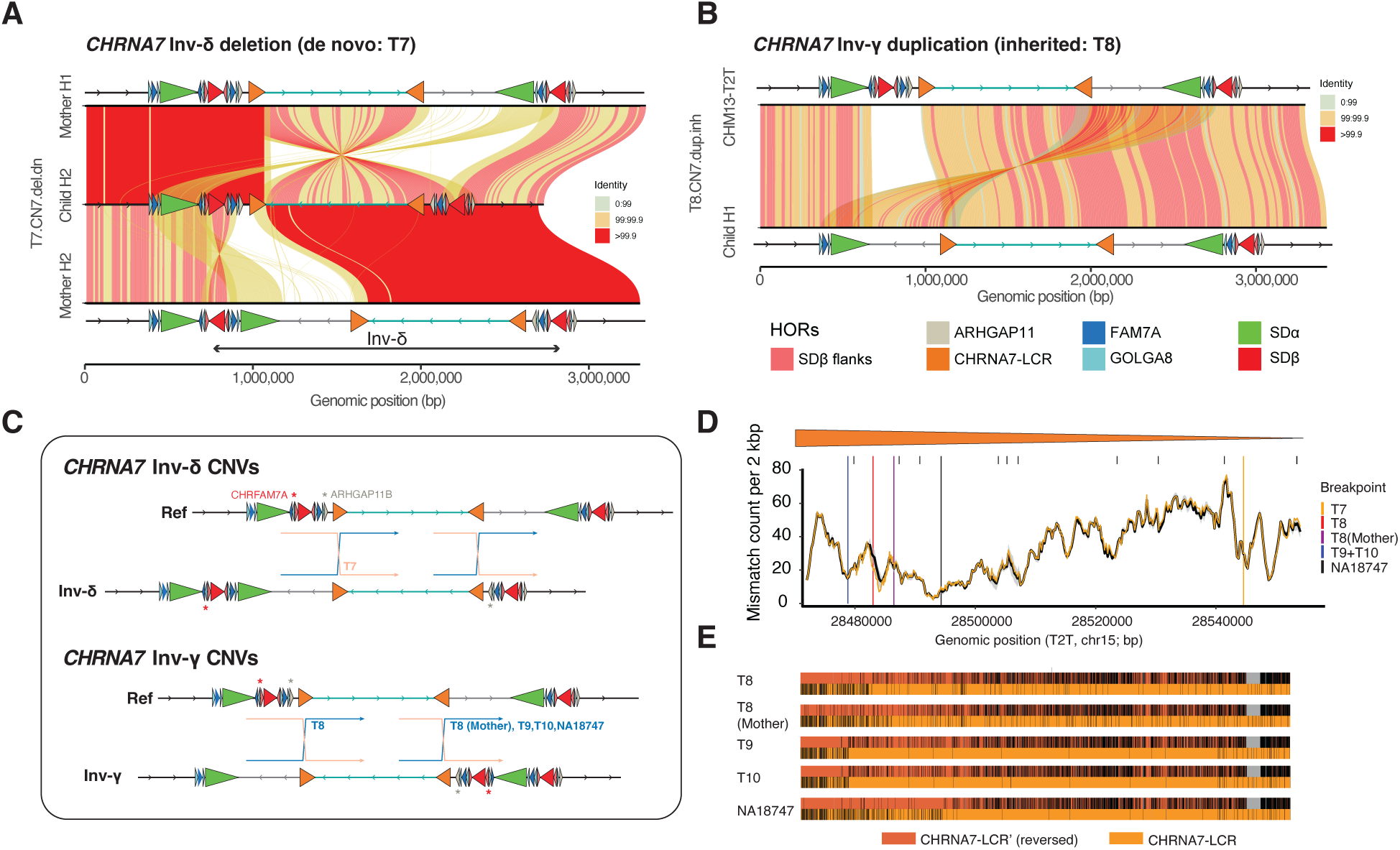
Structure and formation mechanisms of *CHRNA7* CNVs. A) SVbyEye plot showing the de novo *CHRNA7* deletion in T7, with the index allele aligned to the parental alleles. The CNV appears as a rearrangement along CHRNA7-LCR copies (orange triangles) located on a direct and an inverted parental allele. B) SVbyEye plot showing the paternally inherited *CHRNA7* duplication in T8, with the index allele aligned to the T2T-CHM13 reference. C) Derived model for the occurrence of *CHRNA7* CNVs, illustrated via two exemplary genotypes carrying either a heterozygous Inv-δ (top) or Inv-γ (bottom). Red asterisk = *CHRFAM7A*, grey asterisk = *ARHGAP11B* (shown by the colored labels at top of the Inv-δ Ref). Expected crossover breakpoint positions are indicated by blue (*CHRNA7* duplication producing) and orange (*CHRNA7* deletion producing) arrows. We predict NAHR between a reference and an allele carrying either Inv-γ or Inv-δ to give rise to eight subclasses of *CHRNA7* CNVs, three of which were observed in our data. All observed *CHRNA7* duplications are depicted in Supplementary Figure 14. D) Gliding window of the similarity between CHRNA7-LCR and CHRNA7-LCR′ in all control samples with heterozygous Inv-γ or Inv-δ (n=46; 96 CHRNA7-LCR/LCR’ pairs tested), with median and standard deviation indicated as a ribbon. The score of the mother of T7 is overlaid in yellow. The exact breakpoint in T7, and the estimated ancestral breakpoints in the other samples, are indicated by horizontal lines. E) Pairwise alignments between the fused CHRNA7-LCR/CHRNA7-LCR′ segments in CNV alleles, and reference sequences (T2T-CHM13) of CHRNA7-LCR (orange) and CHRNA7-LCR′ (red) per *CHRNA7* CNV allele. Mismatches are indicated in black. The approximate ancestral breakpoint positions emerge as a ‘switch’ from preferred match between the reference alleles. All duplications show signs of ancestral recombination as predicted by our model. T9 and T10 likely originate from the same event, while CNVs in T8 (child, father), T8 (mother), and NA18747 emerged as separate events (Supplementary Figure 15).

### Fusion signatures indicate independent recurrent *CHRNA7* duplication events

As a byproduct of the crossover, we expect *CHRNA7* CNV alleles to carry a CHRNA7-LCR-CHRNA7-LCR′ fusion. To test whether this is the case, we employed paired alignments between the CNV allele’s CHRNA7-LCR and the CHRNA7-LCR and -LCR′ sequences from the T2T-CHM13 reference and counted mismatches between the test sequence and the two reference SDs per bin ^19^. We identified one additional *CHRNA7* duplication allele in the 581-haplotype population cohort ^16^ (sample NA18747) and included it in this analysis. We indeed observed fusion signatures in all 6/6 tested *CHRNA7* CNVs (T7, T8, T8-mother, T9, T10, NA18747), with all inferred breakpoints falling into regions of low sequence divergence (Figure 3D,E, p = 0.0342, one-sided permutation test for four regions within CHRNA7-LCR), and consistent with breakpoint positions observed in ^20^ (Supplementary Figure 15). Only two of the five duplication breakpoints are shared (T9, T10), implying that the *CHRNA7* duplication has arisen at least four times independently. This is in accordance with the notion that *CHRNA7* deletions and duplications are reciprocal events that should arise at the same rates. The concept of allelic versus non-allelic competition does not apply to *CHRNA7* CNVs, as each CHRNA7-LCR has only one homologous ‘candidate’ partner with the same directionality at any time.

In summary, the structural features and fusion patterns point to a shared interchromosomal NAHR mechanism involving heterozygous inversions γ/δ that produces a predictable spectrum of *CHRNA7* CNVs.

### Chromosome 15q13.3 population diversity and inversion polymorphism

To characterize common structural haplotypes at 15q13.3, we analyzed 581 fully assembled haplotypes of healthy individuals from Human Pangenome Reference Consortium (HPRC)^16^, the Human Genome Structural Variation Consortium (HGSVC)^17^ and T2T-CHM13. The dataset represents individuals of diverse ancestries, with a skew towards Africans (AFR: 33.5% of all haplotypes) (Supplementary Figure 16). Among a subset of 551 haplotypes, we distinguish 18 different HOR structures that have been observed at least twice, thus being supported by more than one independent assembly (Figure 4A), while the remaining 30 haplotypes were observed only once (Supplementary Figure 17). We have further validated correct assembly of polymorphic inversion at this locus using Strand-seq and confirm all assessed haplotypes (n=118) being correctly assembled (Supplementary Figure 18). Among 18 structures (referred to as haplotype groups), we observe numerous haplotype configurations, including previously described inversions of the whole region between BP4-BP5 (Inv-γ, Inv-γ-del; haplotype groups 3, 7, respectively) as well as smaller inversions of SDβ and SDβ′ repeats, namely Inv-β (haplotype groups 2, 8, 13, 14, 17) and Inv-β′ (haplotype groups 9 and 17), respectively. The haplotype diversity is also reflected in gene copy numbers, as 10.2% of haplotypes lack *CHRFAM7A*, 0.4% carry two copies of *CHRFAM7A,* and 0.4% lack *ARHGAP11B* (Supplementary Figure 19, Supplementary Data 5). We also tested for associations between these haplotype groups and gene expression within the 15q13.3 region using long-read RNA-seq data available from HPRC for 206 individuals (Methods, Supplementary Figures 20 and 21). Only three weak associations were detected (Hap7-ENSG00000290723, Hap2-HERC2P10, and Hap2-ENSG00000261628). The limited number and modest strength of these associations suggest extensive haplotype recurrence and homogenization across structural classes, consistent with structural variants arising at a rate that exceeds gene-level evolutionary divergence, as expected for recurrent inversion hotspots ^12^.

**Figure 4.**
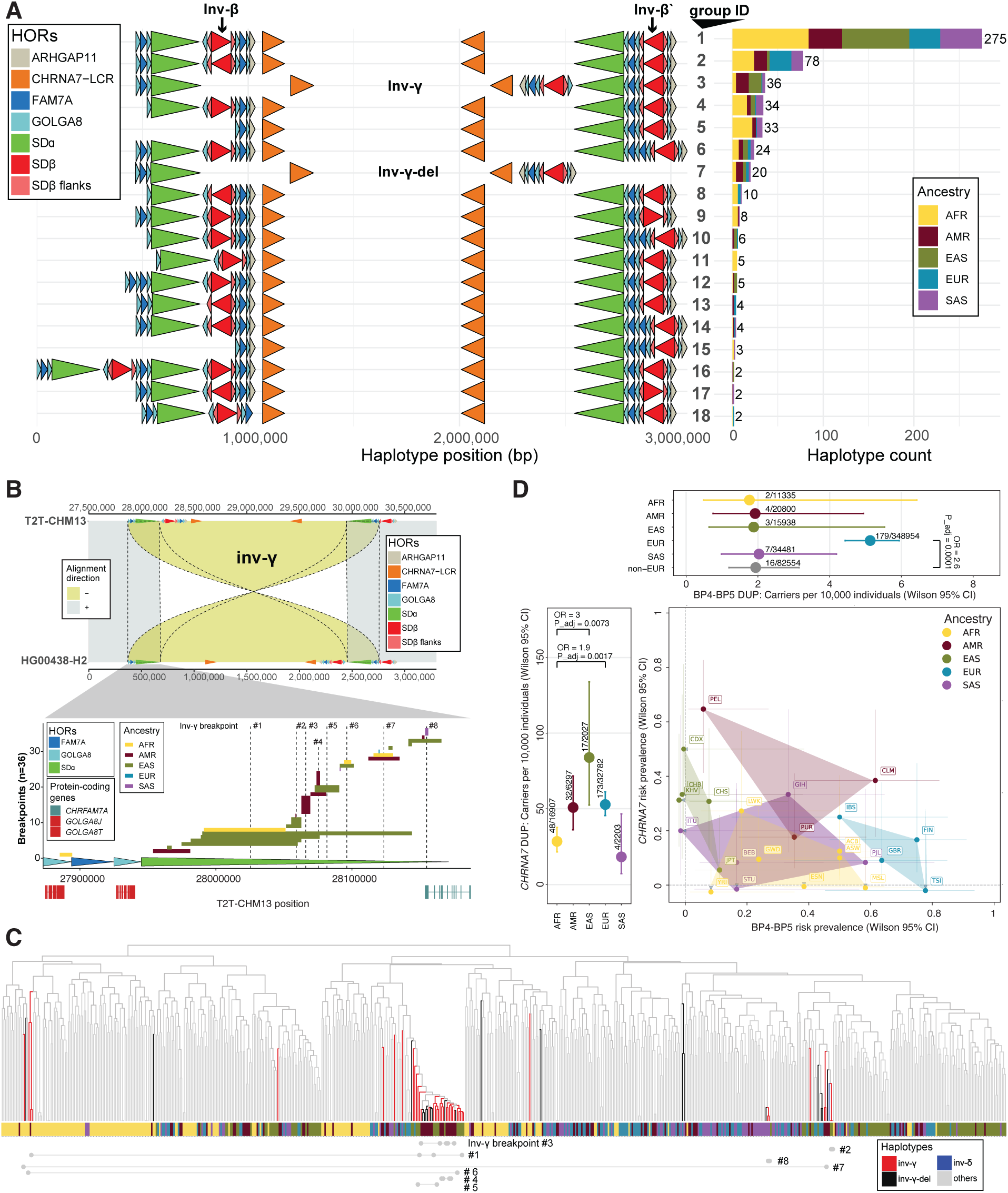
Human population structural diversity of the chromosome 15q13.3 region. A) HOR annotation of the 18 haplotype structures observed at least twice among 581 population haplotypes. Each haplotype structure is defined by forward or backward-pointing arrowheads colored by the HOR unit (see legend). Right side: a stacked barplot showing the frequency of each HOR structure among continental population groups (African (AFR), Admixed American (AMR), East Asian (EAS), European (EUR), South Asian (SAS)). B) SVbyEye plot of a representative Inv-γ, with the direct (reference) haplotype on top and the inverted haplotype below; each carries its HOR annotation above. Dashed outlines highlight redundant alignments at the inverted repeats implicated in inversion formation. Beneath the plot, from top to bottom, are the mapped Inv-γ breakpoint from other samples, the inverted HORs implicated in inversion formation, and the protein-coding genes (*GOLGA8* in red, *CHRFAM7A* in petrol). Inv-γ breakpoints (#1 to #8) are shown relative to a proximal repeat copy, using single reference coordinates from T2T-CHM13. C) Top: Phylogenetic tree based on SNVs in the unique region between BP4 and CHRNA7-LCR of all n=581 population haplotypes (Methods). Branches are colored by the presence of Inv-γ, -γ-del or -δ. Sample ancestry is indicated below each branch. Bottom: Dumbbell plot indicating samples which share a breakpoint region (identified in Panel B). D) *Center:* Scatterplot indicating the percentage of 1kGP samples carrying at least one Inv-β (predisposing to a recurrent BP4-BP5 CNV, x-axis) or a heterozygous Inv-γ, -γ-del or -δ (predisposing to a recurrent *CHRNA7* CNV, y-axis). The 288 samples are grouped by self-reported ancestry obtained from the 1000 Genomes Project panel. Point estimates are shown with 95% Wilson score confidence intervals on both axes (Methods). Shaded hulls enclose the populations of each ancestry (illustrative only). *Top and left panels:* Number of carriers per 10,000 individuals derived from the gnomAD-SV and gnomAD-CNV resources for the *CHRNA7* duplication (left) and BP4-BP5 duplication (top) stratified by ancestry (Methods); points are estimates and whiskers are 95% Wilson score confidence intervals. The number of carriers and samples are indicated next to each dot. Significant two-sided Fisher’s exact tests are indicated by brackets (odds ratio and BH-adjusted p-values).

### Inversion breakpoint mapping

In contrast to previous studies, the current dataset provides fully assembled haplotypes allowing us to map inversion breakpoints at high resolution. This is exemplified by a gapless assembly of a large 5.26 Mbp inversion between BP2-BP3 ^12^, including two novel inversions between BP1-BP2 and BP3-BP4 (Supplementary Figures 22 and 23). The most frequent, the BP4-BP5 inversion, represented by Inv-γ and Inv-γ-del, was observed 66 times among 581 haplotypes (population frequency 11.36%). This inversion is population stratified, which is reflected in a significant depletion of this inversion among samples of African ancestry (p = 2.3 x 10^-4^, Fisher’s exact test, two-sided, Bonferroni corrected) while admixed Americans are significantly enriched (p = 7.3 x 10^-6^, Fisher’s exact test, two-sided, Bonferroni corrected) (Supplementary Figure 24).

We refined the inversion breakpoint of Inv-γ (haplotype group 3) to an average range of 19,861 bp (median: 6,488 bp). All but one breakpoint mapped within the SDα inverted repeat and we observe clustering of inversion breakpoints in eight distinct regions (Figure 4B). The recurrent origin of this inversion is further supported by distinct positioning of inversion breakpoints (Inv-γ breakpoints 2-5 and 8) on a phylogenetic tree constructed from common SNVs (MAF ≥5%) found inside this inversion (Methods), which also suggests inversion breakpoint reuse in different individuals (Inv-γ breakpoints 1, 6, 7) ^12,21^ (Figure 4C). In the case of Inv-γ-del, we narrowed down the inversion breakpoint to an average range of 2,040 bp (median: 529 bp) with the majority of breakpoints (n=14/24) mapping within the FAM7A HOR (Supplementary Figure 25). The remaining inversions (n=10) between BP4-BP5 were observed only in a single haplotype with the majority being mediated by SDα repeats (n=6), while others are more complex (Supplementary Figure 26). Among these, we find a unique instance of Inv-δ in sample HG01192 with inversion breakpoints located within a ∼157 kbp region overlapping with SDβ repeats (Supplementary Figure 27). Using a haplotype concordance test against the 1000 Genomes Project panel, we found this inversion being inherited from the mother (HG01191) along with an additional family duo that carries the inversion (HG01056 - child and HG01054 - father) (Supplementary Figure 28).

#### Population-specific distribution of CNV-predisposing alleles

The CNV formation mechanisms identified in our patient samples allow us to re-interpret population haplotypes with respect to their risk to generate CNVs in their offspring (Figure 4D). Among 581 resolved haplotypes, there are 288 samples for which both maternal and paternal haplotypes were fully resolved (and four with only one haplotype resolved and T2T-CHM13). Of these, 31.9% (92 out of 288) of individuals carried at least one allele predisposing for a BP4-BP5 CNV (haplotype groups 2, 8, 9, 13, 14 and five singletons, Supplementary Figure 17, Supplementary Data 6). The proportion of BP4-BP5 CNV-predisposed individuals was significantly enriched in Europeans (26/38 samples, 68.4%, p = 1.2×10⁻⁵, two-sided binomial test, FDR-adjusted) and depleted in East Asians (3/59, 5.1%, p = 4.16×10⁻^6^, two-sided binomial test, FDR-adjusted), corresponding to a >10-fold population-based risk stratification. *CHRNA7* CNV-predisposed heterozygous Inv-γ and -δ carriers are less frequent (53/288, 18.4%, haplotype groups 3, 7 and ten singletons), with an enrichment in admixed American individuals (20/50, 40%, p = 0.002, two-sided binomial test, FDR-adjusted) primarily driven by a single human population (PEL; Peruvians in Lima; 11/17, 64.7%, p = 0.001, two-sided binomial test, FDR-adjusted). The predicted predispositions for both groups of CNVs are in line with population-stratified orientation of SDβ that is inverted in a significant portion of Europeans (p = 1.66 x 10^-6^, Fisher’s exact test, two-sided, Bonferroni corrected), while East Asians carry mostly the direct orientation (p = 1.66 x 10^-7^, Fisher’s exact test, two-sided, Bonferroni corrected). This puts SDβ and SDβ′ either in direct (in Europeans) or opposite (East Asians) orientation, predisposing to either pathogenic CNVs or recurrent inversions (Supplementary Figure 29). We next compared these predictions to duplication carrier frequencies across ancestry groups in gnomAD v4.1 (Figure 4D, Methods). Indeed, BP4-BP5 duplication carriers were 2.6-fold more frequent in Europeans than non-Europeans (5.13 vs 1.94 per 10,000; p = 3.5 × 10⁻⁵, Fisher’s exact test, two-sided), consistent with the enrichment of predisposing haplotypes. Similarly in line with our predictions, *CHRNA7* duplication frequencies were highest in EAS and AMR (83.9 and 50.8 per 10,000 individuals, respectively) populations whereas they were lowest in AFR and SAS (28.4 and 18.2 per 10,000 individuals, respectively). *CHRNA7* duplications in EUR were more frequent than predicted by haplotype risk alone (52.8 per 10,000), which may reflect recruitment biases, reduced selection against the mildly pathogenic *CHRNA7* duplication or additional mutational mechanisms not observed in our study.

#### Evolutionary history of the chromosome 15q13.3 region across primate lineages

We proceeded to distinguish the ancestral and derived changes in the BP4-BP5 region of chromosome 15q13.3 by performing comparative and phylogenetic analyses of the region among six ape species ^22^ and one Old World monkey (macaque) ^23^. We separately considered the seven distinct duplicons identified in the region (ARHGAP11, CHRNA7-LCR, FAM7A, GOLGA8, SDα, SDβ and SDβ flanks) and constructed phylogenetic trees for each duplicon independently (Methods; Supplementary Figures 30-36). Our analysis suggests that the complexity of duplication structure gradually increases as we approach the human lineage, and this is achieved through the stepwise accumulation of different SDs (Figure 5A). Consistent with the core duplicon hypothesis ^24^, an ancestral locus can be identified for all duplicated segments corresponding to a single-copy ortholog in nonhuman primate genomes, with the exception of FAM7A, which independently expanded in orangutans. We estimate that the flanking SDs (GOLGA8) expanded 9.6 [9.1-10.2] million years ago (MYA) (Supplementary Figure 36) prior to the radiation of the African great apes. Within the complete macaque genome, we find that several of the duplicons are, in fact, highly fragmented, suggesting that they themselves may have been composed of smaller nonsyntenic segments or have subsequently fragmented in the macaque lineage. The juxtaposition of SDα to SDβ flanks and FAM7A to form a larger higher-order unit (329 kbp) is predicted to have occurred >15.2 MYA, and, thus before orangutan divergence. Moving to the African great ape genomes, the HOR 15q13.3 SD grows to 435 kbp and consists of an SD structure of SDα- GOLGA8-SDβ-ARHGAP11. The configuration of SDβ and the two oppositely complemented flanks appears in the common ancestor of human and chimpanzee, a configuration predisposing to Inv-β. The full HOR duplication emerges only in the human lineage, leading to the generation of Inv-δ and Inv-γ.

**Figure 5.**
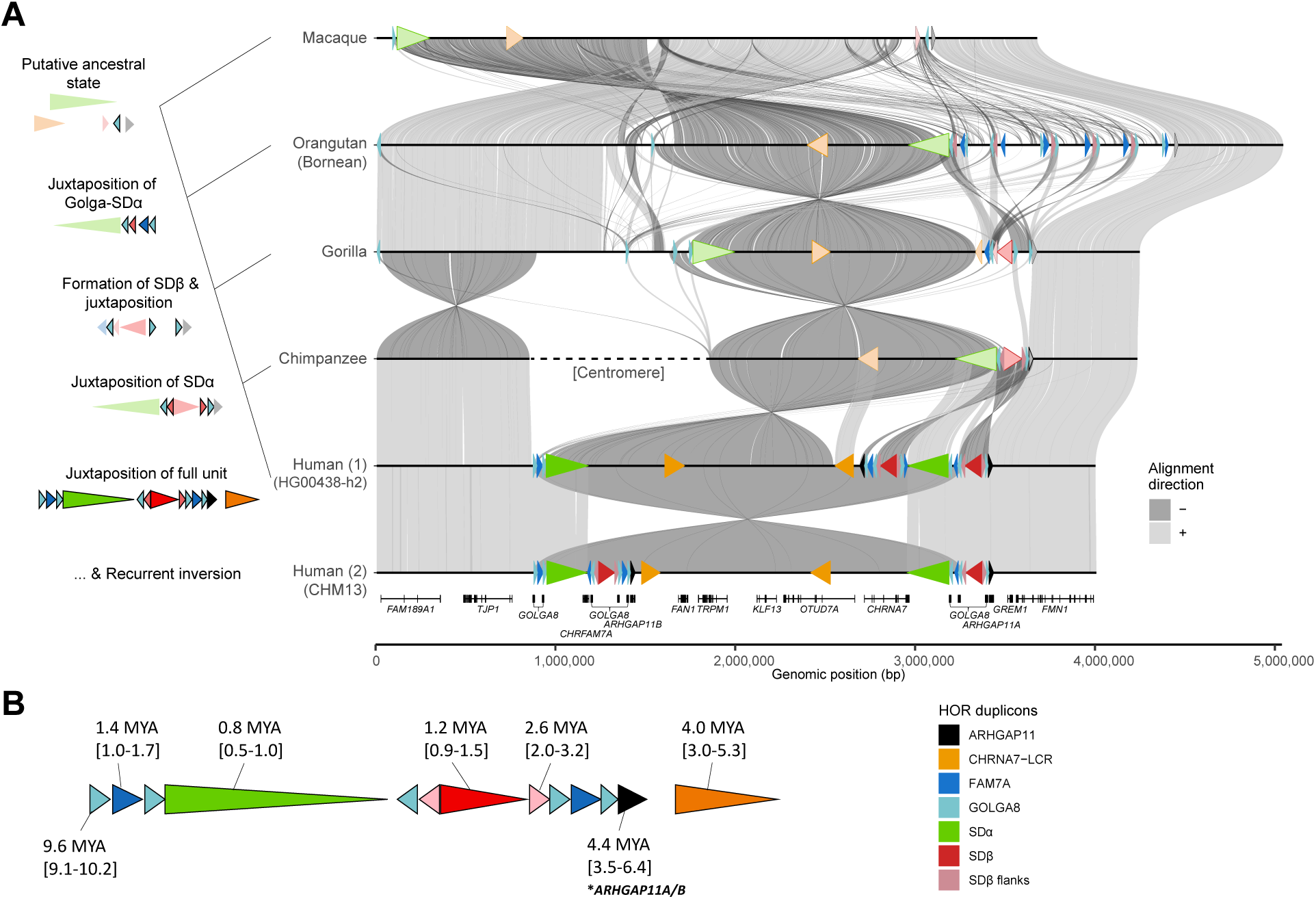
Evolution of the complex duplication structure associated with chromosome 15q13. A) Comparative analysis and model for evolution of the human 15q13.3 locus. The model (left) highlights a series of SD juxtapositions leading to the formation of human-specific duplication structures. The interpretation is based on phylogenetic analyses and comparative analysis (right). Syntenic sequences among complete primate genomes compared to human chromosome 15, 27-31 Mbp (T2T-CHM13 reference coordinates). Species included are macaque, Bornean orangutan, gorilla, chimpanzee, human (1) HG00438.h2, containing Inv-γ, and human (2) T2T-CHM13. HORs are indicated by different colors. Human duplicons that are single copy (unique) in their respective species are indicated by shaded color. Inverted alignments are shown by darker gray. Syntenic genes corresponding to T2T-CHM13 coordinates are shown at the bottom. B) Schematic summarizing the estimated timing of duplication for different duplicons (color) that define the duplication architecture of chromosome 15q13.3 BP4 and BP5. Times and confidence intervals are estimated based on the average coalescence from maximum likelihood trees constructed for each duplicon (Supplementary Figures 30-36), using macaque, Sumatran orangutan, or gorilla as an outgroup.

Based on our phylogenetic analysis, we distinguish two critical time points for human-specific restructuring. They first occurred 4.0 - 4.4 MYA (Figure 5B), where duplication of CHRNA7-LCR and the ARHGAP11 unit containing *ARHGAP11B* formed; a gene that has been implicated in the expansion of the human frontal cortex ^25–27^. The second time point we estimate is much more recent (0.8-1.4 MYA) where we find evidence for the formation of the largest duplications (702 and 482 kbp in size corresponding to BP4 and BP5 duplications, respectively). These structures predispose to NAHR-mediated inversions (*β*, γ, and δ), as well as microdeletions and duplications associated with disease. Leveraging the biallelic SNPs from the syntenic alignment of the fully assembled loci (HPRC2 and HGSVC3; Methods), we screened for Tajima’s D, nucleotide diversity, and haplotype homozygosity tests. Notably, we observe the human-specific duplication architecture that emerged most recently at BP4 maps to the top 1% of extended haplotype homozygosity blocks within chromosome 15 (Supplementary Figure 37) - a potential signal of a recent selective sweep in the human lineage. These findings suggest that the configurations predisposing to NAHR associated with disease may also confer a potential selective advantage in the human population.

## Discussion

It is well established that SDs mediate recurrent inversion and CNV formation through NAHR ^2,12,28^. However, SDs at the most complex loci associated with genomic disorders, such as 15q13.3, are simultaneously hypervariable ^29^ and hard to sequence using short-read techniques ^17^. This makes it challenging to study the variability and pathogenic CNV formation at many hotspots of genomic disease. Through a combination of long-read sequencing and de novo assembly methods, we now observe a level of inter-individual variability of the SD landscape that far exceeds previous estimates ^15^. Making use of a resource of 581 fully resolved haplotypes, we show that there are at least 18 structurally distinct architectures (or 48 including singletons). The high proportion of singleton haplotypes suggests that additional rare haplotypes likely remain to be discovered in larger cohorts and different populations. The structural haplotypes differ markedly in CNVs that they can promote via NAHR as a result of their different SD configurations. In light of this complexity, it is unsurprising that no definite breakpoints for BP4-BP5 or *CHRNA7* CNVs could previously be identified. In our study, identification of fully resolved population, parental, and patient haplotypes served as an essential scaffold for understanding the exact mechanisms underlying CNV formation in 15q13.3 patients.

### BP4-BP5 CNVs

The common inversions β and β′, emerged as the most important modulators of recurrent BP4-BP5 CNV risk. Among six BP4-BP5 CNVs in our patient group, five likely arose by NAHR involving the SDβ repeat affected by Inv-β or Inv-β′, which bring >100 kbp of >98% identical sequence into direct orientation, enabling deletion or duplication of ∼1,948 kbp of intermediary sequence. This mode of BP4-BP5 CNV formation has been suggested before ^15^ but could only now be definitively observed in patient data, due to the difficulty to ascertain Inv-β/β′ reliably without long-read assemblies. Within SDβ, three breakpoints cluster in just 2 kbp around a recombination hotspot region, suggesting that this region may be a driver of BP4-BP5 formation. For two CNVs (T1, T5), at least one breakpoint overlapped with a *GOLGA8* element, consistent with their known role as core duplicons ^24^ and promoting disease-associated genomic instability at 15q13.3 and 15q24 ^15^. Only one out of six BP4-BP5 CNVs arose via a nonhomology driven mechanism, with the breakpoint instead displaying 3 bp of microhomology, pointing to a replication-based mechanism such as microhomology-mediated break-induced replication (MMBIR) or fork stalling and template switching (FoSTeS) ^28^. The low allele frequency of the driving haplotype (Inv-β+Inv-γ) in controls (1 of 581; 0.17%) suggests that this configuration may be recurrently unstable and thus under purifying selection.

### *CHRNA7* CNVs

The second group of 15q13.3 rearrangements, *CHRNA7* CNVs, also arise predominantly by NAHR, however via a structurally different process. Homologous recombination is, by default, suppressed in a heterozygous inverted state ^30^. We found that the repeat pair CHRNA7-LCR and CHRNA7-LCR′ inside the inversion can undermine this suppression and allow for interchromosomal recombination inside the locus-spanning heterozygous Inv-γ or Inv-δ. The result of this fusion process is a reciprocal pair of a *CHRNA7* duplication and deletion. Despite their name, both are in fact mixed events in which ∼426 kbp, including *CHRNA7,* are duplicated (or, double-inherited) and ∼301 kbp, including *ARHGAP11B* are deleted (*CHRNA7* duplication), and *vice versa* for the *CHRNA7* deletion. *CHRFAM7A* is affected in CNVs arising from Inv-γ, but not in those from Inv-δ. This mechanism agrees with a previous study in which 50/55 of *CHRNA7* CNVs were proposed to arise this way ^8,20^. The joint generation of *CHRNA7* deletions and duplications suggests that the de novo rate of both should be similar, which is further supported by our breakpoint analysis that showed 4/5 *CHRNA7* duplication alleles arose independently. Given that *CHRNA7* duplications are relatively frequent in the general population and typically show low penetrance with minimal phenotypic effects, the lack of reported de novo cases ^9^ is likely attributable to ascertainment bias. Finally, the finite number of possible NAHR recombinations allows us to predict eight distinct *CHRNA7* CNVs, three of which we indeed observed in our data. We further expect to see similar deletion/duplication pairs in other recurrent CNV loci whenever a repeat pair is embedded in a large inversion region.

### The individual risk for 15q13.3 CNVs is unequally distributed

The risk for 15q13.3 CNVs is unevenly distributed across populations. European haplotypes are enriched for configurations predisposing to BP4-BP5 CNVs (68.4% of individuals carry at least one Inv-β/β’), making them more frequently predisposed to BP4-BP5 CNVs, whereas these alleles are largely depleted in East Asians (5.1% of individuals predisposed by an Inv-β/β′). These findings mirror other genomic disorders such as Koolen-de Vries syndrome ^31^ and the 22q11.2 microdeletion ^32^, whose recurrent CNV rates are likewise gated by population-stratified structural haplotypes. Given that all ten trios are of European ancestry, our analysis is thus likely biased towards Inv-β/β′ mediated CNVs, while in other populations other mechanisms may be predominant, or selective forces may also shape the allele distribution, as suggested by the notably extended haplotype homozygosity and low Tajima’s D signals observed among East Asians (Supplementary Figure 37).

We also tested whether (1) elevated genome-wide recombination rates or (2) rare PRDM9 alleles may have accompanied the de novo CNVs but found no evidence for either (Supplementary Data 7, Supplementary Figures 38 and 39). We lastly propose a model of ‘competition’ between allelic and non-allelic homologous recombination, in which the latter is made more likely if paralogs provide a better local sequence match than syntenic homologous chromosomes during meiosis. Competing repeats engaging in NAHR in trios T2-T4 had particularly high sequence similarity near their breakpoints, which may have made their pairing more likely. Although we see this as an interesting model that could be relevant for other recurrent CNVs, this observation requires independent validation in larger cohorts.

### Genes, phenotypes and disease severity

15q13.3 CNVs show marked but poorly understood phenotypic variability ^33–35^. First, additional variants outside the locus (“second-site hits”) likely contribute to 15q13.3-associated phenotypes ^33^: Additional variants are indeed present in three trios (T3, T5, T8, Supplementary Data 8), and three patients have one sibling with ASD (T3, T7) or receiving special education (T5). For two of these siblings, CNV status was confirmed absent (Supplementary Figure 40), showcasing the presence of additional factors in these families. Our data, however, also highlight structural heterogeneity between 15q13.3 CNVs at the sequence level that affect different genes. For example, the BP4-BP5 deletion of T6 affects *CHRFAM7A*, and Inv-γ mediated *CHRNA7* duplications are accompanied by deletions of *CHRFAM7A* and *ARHGAP11B* (Supplementary Figure 41). Further complicating matters, 0.4% of population haplotypes lack *ARHGAP11B* and 10.2% lack *CHRFAM7A*. This raises the possibility of phenotypes in 15q13.3 patients arising through a complex interplay of affected genes on both alleles. Given our relatively small and heterogeneous cohort, these observations remain preliminary and require validation in larger cohorts before implications for genotype-phenotype relationships can be drawn. If supported by future studies, these findings may ultimately improve genetic counseling by providing an assessment of recurrent CNV risk and predicted disease severity.

### Human 15q13.3 evolved recently in the great ape lineage

It seems counter-intuitive at first that the extensive segmental duplication in 15q13.3 rose to high frequency in humans despite their role in generating genomic instability associated with disease. A cross-species comparison of the region in humans and six great ape species reveals that this level of instability at 15q13.3 is specific to the human form of the locus, which is the product of a series of substantial rearrangements occurring recently in the hominid line. The key duplication events took place primarily at two timepoints, including 4.0-4.4 MYA corresponding to expansion time of duplicons (CHRNA7-LCR and ARHGAP11) overlapping with brain-critical genes, *ARHGAP11B* and *OTUD7A.* The second relatively younger timepoint of expansion (0.8-1.4 MYA), corresponds to coalescent times of FAM7A, SDβ and SDα duplicons, harboring inversion breakpoints (Figure 1B), which could again affect genes implicated in the evolution of the human brain, e.g. *CHRFAM7A, ARHGAP11B, OTUD7A* and *CHRNA7.* The human locus may pose the best readily available compromise between retaining these critical duplications, while minimizing the risk for recurrent copy number variation - the latter achieved by maintaining the major repeat pairs in opposite orientation in most haplotype configurations.

Our study demonstrates that investigating genomic regions previously inaccessible to standard sequencing approaches can provide important new insights into structural variation and the mechanisms underlying CNV formation. Such analysis also provides greater precision with respect to the genes deleted in patients especially those near or within the segmental duplications, e.g. *CHRFAM7A*, *ARHGAP11B* and *OTUD7A* where exact breakpoints were difficult or impossible to define based solely on microarray and short read-sequencing. Beyond 15q13.3, several additional genomic loci in which SD-mediated CNVs cause disease to remain insufficiently characterized. Generating complete haplotype-resolved assemblies and achieving base-pair resolution of CNVs in these complex regions will be essential for improving genotype-phenotype correlations and for advancing our understanding of mutation mechanisms and locus evolution.

## Methods

Our research complies with all relevant ethical regulations. Ethical study approval was granted under references S-212/2023 (Ethikkommission Medizinische Fakultät Heidelberg, University Hospital Heidelberg), dossier nr. 2023-16530 (Commissie Mensgebonden Onderzoek Radboudumc, Radboudumc, Nijmegen) and 100/2025 Render (PNRR-MR1-2023-12377314) (Comitato Etico Campania 3, AOU Federico II, Napoli, Italy).

### Patient recruitment and phenotypic data collection

We recruited 10 individuals with large CNVs at 15q13.3 from 10 unrelated families. All patients had been identified through standard-of-care genetic diagnostics and were referred to clinical genetics departments at Radboudumc (n=6, T1,T2,T3,T4,T5,T7), TIGEM; University of Naples (n=1, T8), or University Hospital Heidelberg (n=3, T6,T9,T10) for evaluation of neurodevelopmental or neurological symptoms. Patients were included in the study based on available sample collections in each of the three centers. Parents of all affected individuals were included regardless of CNV status. Phenotypes were recorded using Human Phenotype Ontology (HPO) terms and are summarized in Supplementary Data 1. Note that index T8 is part of a monozygotic twin pair where both twins carry the identical 15q13.3 CNV, and of which only one was included in the study. Upon receiving comprehensive information regarding the study’s objectives, all participants or their legal guardians provided written consent according to institutional ethical guidelines.

### HiFi sequencing

We generated HiFi long-read genomes on a *PacBio Revio* platform for all 10 families, totalling 30 individuals. We initially used one SMRT Cell per sample, but re-ran samples that yielded <20-fold coverage on the first run, eventually yielding a median depth of 27.5X (30/30 samples: >20-fold coverage, 25/30 samples: >25-fold coverage) and median read length of 16.4 kbp (Supplementary Data 2). All samples were processed in the same fashion, according to the manufacturer’s instructions (PacBio, Menlo Park, CA, USA). In brief, 7 µg gDNA was sheared on Megaruptor 3 (Diagenode, Liège, Belgium) to a target size of 15-18 kbp, libraries were prepared with SMRTbell prep kit 3.0 (PacBio, Menlo Park, CA, USA), size-selected >10 kbp on the BluePippin (Sage Science, Beverly, MA, USA), and sequenced for 24 h on the Revio system (ICS 12.0.4).

### HiFi data processing, alignments, initial variant calling

Alignment (pbmm2 v1.10.0) of HiFi reads was performed against the GRCh38 reference genome, which is gapless in the region and structurally identical to the T2T-CHM13 reference (Supplementary Figure 42). Depth-based CNV calling was performed using HiFiCNV v0.1.6.

### ONT sequencing

We generated ONT sequencing data for T2-Father, T7-Index and T7-Mother. High molecular weight (HMW) gDNA was extracted from frozen blood samples using the NEB Monarch HMW DNA extraction kit for Cells & Blood (#T3050L) following the manufacturer’s protocol with the following exceptions: 500 uL blood was used for the starting input with a shaking speed of 600 rpm during the lysis step. DNA was precipitated with 300 uL EEB (ONT) and solubilized at 4°C for two days. Libraries were constructed using the Ultra-Long DNA Sequencing Kit V14 (SQK-ULK114) following manufacturer’s protocol and were loaded onto a FLO-PRO114M R10.4.1 flow cell for sequencing with two nuclease washes and reloads after 24 and 48 hours of sequencing.

### Read-based confirmation of CNV carriership

We used the BedTools (v2.31.1) ^36^ “intersect” function to test for at least 25% reciprocal overlap between CNVs expected based on standard-of-care and depth-based CNV calls generated with HiFiCNV, confirming all expected 15q13.3 CNVs. To exclude the possibility of false or missing calls, we also created views of aligned reads, depth profiles (derived from HiFiCNV) and b-allele frequencies (derived from DeepVariant version 1.8.0 ^37^) in the Integrative Genomics Viewer (IGV, version 2.19.1) ^38^ for each sample to visually confirm the diagnoses (Supplementary Figures 43-52). With one exception, all CNV breakpoints were located inside BP4, BP5, or CHRNA7-LCR′ duplication blocks where they could not be identified using read mapping.

### Genome assembly generation with hifiasm

We generated de novo assemblies using *hifiasm* (v0.24.0-r702) ^39^ with the offspring samples run in trio mode with parental reads to guide phasing. To reduce computational time and allow for flexible parameter testing, we restricted assembly to chr15:25-50 Mbp by aligning all HiFi reads to GRCh38 using minimap2 (multi-mapping of reads enabled) and extracting the relevant subset of reads with SAMtools, before converting them back to fastq with the *samtools fastq* command. We tested different values of the hifiasm “-a” parameter, which controls the amount of assembly graph cleaning, and used “-a 1” for the index patient of T2 and the father of T8, so we could prevent assembly breaks appearing in default settings. In the case of the father of T2 and the mother and index of T7, assembly was performed using HiFi and ONT data jointly, by utilising the “-ont” option in hifiasm. Assembly statistics are summarized in Supplementary Data 9.

### Duo-phasing for parental assemblies of de novo CNVs

Duo-phasing of parental assemblies was performed for three families with de novo CNVs without ONT data (T1, T3, T4), with an approach inspired by the graphasing ^40^ assembly phasing workflow: Trio-phased child assemblies were first generated and validated. For transmitting parents (T1-father, T3-mother, T4-mother), unphased unitigs were produced with hifiasm ^39^ and assigned to “inherited” or “not inherited” categories using yak triophase, based on transmission to the child. Guided by assembly graph visualization in Bandage ^41^, parental unitigs were ordered to match child inheritance, and a custom script was used to merge unitigs into full contigs. Transmitting parents of T2 and T7 were already fully phased due to ONT sequencing.

### Assembly validation

To account for the possibility of assembly errors, all fully and partially resolved assemblies were carefully evaluated by the following procedures.

<u>(1) Coverage and b-allele frequency profiles.</u> We first validated the structural content of each assembly against read-based coverage and b-allele frequency (BAF) profiles. CNV segments from the assemblies were compared to those inferred from depth and allele-balance metrics to confirm consistency with expected large-scale duplications and deletions in BP4-BP5 and *CHRNA7* (Supplementary Figures 43-52).
<u>(2) Gene copy numbers.</u> We inferred the copy numbers of 15q13.3 genes (*CHRFAM7A, ARHGAP11B, MTMR10, TRPM1, KLF13, FAN1, OTUD7A, CHRNA7, ARHGAP11A*) both based on aligned reads and based on our assemblies. To overcome the difficulty of copy number estimation in repetitive regions, which affects most of these genes, we devised a dual method for read-based calling, where we called the copy number based on normalized depth of reads mapping to the gene, and by using devider ^42^ to estimate the number of differing haplotypes (methods - devider). Assembly-based calling was done by counting the number of matches any gene sequence, extracted from T2T-CHM13, has with the assembled haplotypes (counting assemblies that cover at least 90% of the gene sequence). Out of 270 sample-gene pairings (30 samples * 9 genes), 262 (97%) copy number estimates differed by less than one between the methods. Manual investigation of the remaining eight assignments indicated that the depth-based method had been overestimating copy numbers due to local fluctuations in read depth, eventually confirming all gene copy numbers (Supplementary Data 10).
<u>(3) Reads-to-assembly mapping.</u> Reads from each sample were mapped simultaneously against the two assembled haplotypes of that sample using minimap2 v2.30.0 ^43^, allowing reads to preferentially align to the best-supported sequence. Validation was performed in two ways. First, coverage profiles were inspected and showed regular coverage across all assemblies, with no internal regions lacking read support (Supplementary Figures 53-62). Second, structural variants were called using Sniffles v2.7.2 ^44^, as misassemblies are expected to appear as disagreements between reads and assembly sequence. Across all 60 haplotypes, 17 structural variants (52-654 bp) were detected (Supplementary Data 11). Sixteen were heterozygous and, upon manual inspection, were explained by reads mapping to the incorrect haplotype due to known gaps in the alternative haplotype. One homozygous event, a 90 bp insertion within a GA-rich repeat, was manually confirmed and is most consistent with mosaicism rather than an assembly error.
<u>(4) Breakpoint validation in long reads.</u> For all de novo breakpoints (T1-4,T7), we confirmed the presence of the breakpoint sequence in raw reads in the index patient, and absence thereof in the respective parents (Supplementary Data 12).
<u>(5) Population-based validation of large inversions.</u> To independently assess inversion calls, assembled haplotypes were compared to structurally resolved haplotypes from the HPRC dataset. A phylogenetic tree was constructed using common SNVs (MAF ≥5%) within the unique region between BP4 and CHRNA7-LCR′ (Methods). Patient and parent haplotypes inferred from de novo assemblies to carry Inv-γ or Inv-δ were evaluated for clustering with inversion-carrying haplotypes in this framework. All predicted inversion carriers grouped within inversion-associated clusters, whereas no other patient/parent haplotypes did so, supporting the inversion assignments (Supplementary Figure 63).
<u>(6) Correspondence to population haplotype groups.</u> We compared our de novo assemblies to the 48 haplotype groups identified in the independent population dataset. All resolved patient/parent haplotypes without BP4-BP5 and *CHRNA7* CNVs matched structurally with one of the 48 population haplotype groups (Supplementary Data 3).
<u>(7) Orthogonal validation with optical genome mapping.</u> Optical Genome Mapping (OGM) data was generated for 8 individuals from 4 families to orthogonally validate LRS-based assemblies (Supplementary Data 13). Ultra-high molecular weight DNA was isolated from EDTA blood using the Ionic Purification System and beta testing protocol and consumables for this application (Bionano Genomics, Inc). Labeled DNA was prepared using the Direct Label and Stain Labeling Kit (Bionano Prep DLS-G2, Bionano Genomics), loaded onto Stratys Core Chips, and imaged on a Stratys Instrument. Genome maps were assembled using the Guided Assembly pipeline in Bionano Access 1.8.3 and aligned to GRCh38. Non-reference junctions in OGM assemblies were compared to those in the LRS-based assemblies aligned to the same reference. To address the case where no consensus OGM assembly could be reached (T7-mother), we additionally developed an assembly-free validation approach: CNV-specific sequences of 300 kbp spanning structurally diagnostic junctions were extracted from the LRS assemblies and converted to predicted OGM label patterns. Raw molecule files (.bnx) were then queried for molecules matching these expected patterns using a custom script (see Code Availability).

Estimation of distinct alleles. Estimation of distinct alleles was performed using devider ^42^. Unlike standard phasing algorithms, which assume a fixed number of haplotypes (e.g., WhatsHap ^45^), devider dynamically determines the number of distinct haplotypes, enabling quantification of haplotype or gene counts at any position (Supplementary Data 10; Supplementary Figures 6 and 64). Analyses were performed iteratively in 10 kbp windows: reads overlapping a given position were phased with devider, and the number of haplotypes recorded, providing estimates every 10 kbp. For gene copy number estimation (assembly validation), all reads overlapping the gene midpoint were used for haplotype estimation.

### Definition of higher order structure (HOR) at 15q13.3

We first aligned extracted FASTA sequence from T2T-CHM13 reference at chr15:27500000-30800000 to itself using minimap2 (v2.24) with the following parameters: -x asm20 -c --eqx -DP -m200. Next, we took alignments that are >=20 kbp in size and at least 1 kbp apart. We first processed alignments with a single extra copy within the T2T-CHM13 sequence. For each such alignment we extracted the sequence from both locations and constructed multiple sequence alignment (MSA) using the R package DECIPHER (v2.28.0) ^46,47^. Then we reported a consensus sequence for each MSA alignment. Based on visual inspection of each MSA we allowed gaps of <=5 kbp to allow for presence of occasional structural variants between paralogous copies of HOR units. We followed by processing alignments that are shared among multiple locations with the T2T-CHM13 sequence. We stratified these by size and, again for each size group of self-aligning regions, we constructed an MSA and reported a single consensus sequence as described above. In total, we report seven consensus sequences that define the HOR structure of the region. We define the position of these HOR units in each haplotype by aligning reported consensus sequences to each haplotype using minimap2 (v2.24). We filter out partial alignments smaller than 1 kbp. We note that our HOR structures do not capture structural variation within paralogous copies of each HOR unit but rather smooth over them. In order to obtain such fine-scale differences we advise to use the DupMasker tool or sequence alignments between paralogous copies.

### Definition and assignment of 48 haplotype groups

As described in the section “Definition of higher order structure (HOR) at 15q13.3” we have defined the HOR structure of each selected haplotype (n=581). Then in every haplotype we assign each HOR unit a unique numeric identifier. This identifier is either a positive or a negative number for direct and reverse oriented HOR units, respectively. Next, we calculate the distance between each possible pairs of haplotypes as described before ^32^. All pairwise distances are organized into the distance matrix, which is then used to construct the UPGMA tree using the function ‘upgma’ from the R package phangorn (v2.11.1) ^48^. By visual inspection of the clustered haplotypes we define 48 nonredundant haplotype groups that are separated by the ‘cutree’ function (set parameter: k=48) from the base R package.

### Relative Homology Advantage (RHA) quantification

We defined a “relative homology advantage” (RHA) score to quantify whether local sequence homology favours non-allelic over allelic pairing at SDβ. We extracted the region spanning the high-scoring PRDM9 motif on SDβ (±10 kbp) from the T2T-CHM13 reference and mapped it to each haplotype assembly to recover all copies of the region, expected to be one SDβ and one SDβ′ per haplotype. Each recovered copy was mapped back to the reference and single-nucleotide variants were called jointly across all copies with bcftools mpileup. RHA is defined over an ordered triplet: a non-allelic (NAHR) partner, the true allelic partner, and a deciding partner against which both are compared. RHA is defined as RHA = d(SDβ′₁, SDβ′₂) − d(SDβ, SDβ′₂), where d counts disagreeing SNPs over the central 2 kbp window (the ±10 kbp region served only for extraction). Positive values indicate that the non-allelic SDβ-SDβ′ pairing matches the deciding sequence more closely than the true allelic pairing does, a configuration expected to favour NAHR. Each sample with one SDβ and one SDβ′ per haplotype yields four such triplets, and hence four RHA scores.

We computed RHA scores for the three SDβ-SDβ′ pairings on which the de novo CNVs in families T2–T4 arose. As a background, we drew on the 288 individuals whose HPRC and HGSVC assemblies resolve 15q13.3 gaplessly on both alleles. Because RHA is not defined for rearranged or duplicated haplotypes, we retained the 153 samples in which both haplotypes carried exactly one SDβ and one SDβ′ copy with no locus-spanning inversion, yielding 612 background RHA scores (108 for EUR samples) that were compared against the three parental scores.

### Expression analysis

Long-read transcript expression data were obtained from the Human Pangenome Reference Consortium (HPRC) via the Human Pangenome Data Explorer for 206 lymphoblastoid cell line (LCL) samples generated using PacBio SMRT Iso-Seq/Kinnex full-length non-chimeric (FLNC) reads. All samples overlapped with the haplotype-resolved assemblies used in this study. FLNC reads were aligned to GRCh38 using pbmm2 v1.13.1 with the --preset ISOSEQ option and default parameters. Isoform processing and quantification were performed using IsoQuant v0.17.0 (profile, merge, and call steps) with reference annotation from GENCODE v49 ^49^. Gene-level counts were obtained by summing transcript counts per gene. Expression values were normalized to TPM using the total number of FLNC reads per sample. Analyses were restricted to genes with transcription start sites within chr15:29,000,000-33,500,000 on hg38. For each retained gene and each of the 18 haplotype groups analysed, the association between expression and haplotype dosage (0/1/2) was tested with a two-sided Spearman rank correlation. p-values were adjusted for multiple testing using the Benjamini-Hochberg correction.

### De novo breakpoint identification and validation

Briefly, we first annotated SDs, genes, and CNVs on all assemblies, and generated visualizations to assess structural context and identify approximate breakpoint locations by visualizing alignments between child and parental assemblies using NAHRwhals ^14^ and SVbyEye ^50^. Breakpoints were orthogonally confirmed using a newly devised tool, *crosshair*, which utilizes the assembly unitigs to identify recombination sites as transitions from perfect to SNP-rich alignment (Supplementary Figure 65), followed by a local three-way MSA between the affected allele and both parental haplotypes (Supplementary Figures 6B and 66). As a final layer of validation, we used k-mer analysis to show that each breakpoint-specific sequence was present in the child but absent in both parents (Supplementary Data 12).

### Phylogenetic tree construction of population haplotypes

To generate phylogenetic trees regarding carriership of inversions γ, γ-del, and δ (Figure 4C), all assemblies were first aligned to T2T-CHM13 using minimap2 v.2.30.0 ^43^ with preset -*x asm5*. Variants were then called from these alignments using the BCFtools v.1.23 ^51^ functions *mpileup* and *call*. For each sample, we noted the presence of inversion γ, γ-del, and δ using the haplotype group visualizations as a basis. The multi-sample VCF and inversion labels were then used as input for a custom R script (R version 4.4.1), in which phased SNP genotypes were converted into haplotype matrices, filtered to retain only biallelic SNPs with minor allele frequency >5%, and pairwise genetic distances were calculated using Euclidean distance across SNPs. Hierarchical clustering using the *hclust* function in R was performed on the resulting distance matrix, and the dendrogram was converted to a phylogenetic tree using the ape package for visualization.

### Population BP4-BP5 and *CHRNA7* CNV risk estimation

Assignment of risk groups for BP4-BP5 and *CHRNA7* CNVs per individual from the population cohort was based on the presence of inversions on both haplotypes: samples were considered predisposed for BP4-BP5 CNVs if they carried an Inv-β, Inv-β′ or inversion of SDα ("Inv-α") on at least one haplotype (group.IDs: 2,8,9,13,14,20,31,36,40,46). *CHRNA7* CNV predisposition was assumed if one haplotype carried Inv-γ, Inv-γ-del, Inv- δ or a similar rearrangement (group.IDs: 3,7,19,26,28,29,30,31,32,33,35,37) on one haplotype, and a direct-facing on the second haplotype (group IDs: 1,2,3-6,8-18,20-25,27,34,36,38-48). The proportion of predisposed individuals per population was estimated with a 95% Wilson score confidence interval ^52^ to account for finite per-population sample sizes.

### Ancestry-stratified CNV carrier frequencies

Carrier frequencies of BP4-BP5- and *CHRNA7* duplications across genetic ancestries were obtained from the publicly available gnomadSV and gnomadCNV databases ^53^. For the BP4-BP5 interval, carrier counts were obtained from the gnomAD v4.1 exome CNV callset (464,297 individuals). The *CHRNA7* duplication interval does not fully encapsulate coding genes and is therefore absent from the exome-based gnomAD CNV callset; carrier counts were instead obtained from the gnomAD v4.1 structural variant callset (63,046 individuals, GATK-SV). "Non-Finnish European" and "Finnish" (NFE and FIN) groups were merged into a common EUR stratum; Ashkenazi Jewish, Amish, Middle Eastern, and "Remaining" groups were excluded. Ancestry-stratified carrier frequencies were expressed per 10,000 individuals with 95% Wilson score confidence intervals. Differences in carrier frequency between ancestry groups were assessed using two-sided pairwise Fisher’s exact tests and p-values were Bonferroni corrected. The pre-specified EUR vs non-EUR contrasts were tested separately by two-sided Fisher’s exact test and corrected across the three pre-specified hypotheses (Benjamini–Hochberg). Effect sizes are reported as odds ratios with 95% confidence intervals. Fisher’s exact test yields no test statistic or degrees of freedom. Deletions were not present in sufficient numbers in these databases for meaningful ancestry-stratified analysis (7 and 1 non-European alleles for BP4-BP5 and *CHRNA7* deletions, respectively).

### Genome-wide crossover analysis

We developed a method to detect meiotic crossovers from trio-based long-read sequencing data. As a first step, we filtered out high-quality input variants to improve phasing and reduce false positives as follows. We excluded variants overlapping known problematic regions, including centromeres, reference gaps, SDs, and regions flagged by the Genome in a Bottle (GIAB) Consortium ^54^. Additional filters removed variants with QUAL < 30, genotype quality (GQ) < 20, read depth (DP) < 10, or allele frequency (AF) < 0.2. Crossover detection was performed with WhatsHap ^45^ (v2.3) for each autosome using the --recombination-list flag. We phased each child in the trio using corresponding parental BAM files and retained only simple crossovers: those with no other recombination detected within ±100 kbp. This threshold follows the criteria used by Palsson et al. ^55^ where switches within <100 kbp were considered possible gene conversions or complex recombination clusters. We used this filtered set of crossovers to analyse crossover rates in the transmitting parents of de novo CNVs and compare these to nontransmitting parents. To validate our pipeline, we benchmarked against the G3 recombination dataset published by ^56^, which combines inheritance vectors, Strand-seq, and phased assemblies into a high-confidence map (Supplementary Figure 38). Compared to ^56^, we observe a median of ∼70% precision and ∼50% recall of crossover events. While this performance hints at remaining problems with crossover calling, we reason that this approach displays sufficient precision to flag outliers in genome-wide recombination rates.

### Evolutionary history of common repeat segments and Inv-γ

The evolutionary analysis focused on seven duplicon units (ARHGAP11, CHRNA7-LCR, FAM7A, GOLGA8, SDα, SDβ and SDβ flanks; Supplementary Data 14) across a broader panel of primates, including six great ape genomes ^22^ and a macaque genome ^23^. To identify orthologous copies, sequences were aligned to both macaque and nonhuman ape genomes with minimap2 v2.28 ^43^ using the parameters ‘-cx asm20 --secondary=yes -N 1000 -eqx’. Orthologous repeat segments were subsequently aligned with MAFFT v7.525 ^57^ with ‘--auto’ option. Conserved regions corresponding to each duplicon unit were obtained with Gblocks v0.91b ^58^ for phylogenetic reconstruction. Maximum-likelihood trees were generated in IQ-TREE v2.3.6 ^59^ with 1000 bootstrap replicates, using macaque orthologs as the outgroup. Divergence times were inferred by recalibrating against the estimated macaque-human divergence of 28.8 MYA ^60^. Confidence intervals were computed by replicating 100 trees with the IQ-TREE parameter ‘--date-ci 100’.

More specific phylogenetic dating of inversion γ was estimated by identifying the breakpoints of Inv-γ. Putative breakpoints were identified by aligning the haplotypes with and without the inversion (HG00438.h2 and T2T-CHM13v2.0, respectively), specifically examining the duplicon cluster and its 100 kbp flanking sequence where the Inv-γ breakpoints map. Two 50 kbp windows located upstream and downstream of the breakpoint were assessed for the phylogenetic tree reconstruction. The putative age of Inv-γs was predicted by computing the coalescent time among the haplotypes containing the inversion, recalibrating the time estimated by chimpanzee-human divergence of 6.4 MYA ^60^.

### Selection signature

The selection signature across 15q13.3 (chr15:27-31 Mbp) was assessed based on the syntenic alignment of assemblies of human samples, including genome assemblies generated by Human Pangenome Reference Consortium release 2 (HPRC2; n=233) and Human Genome Structural Variation Consortium release 3 (HGSVC3; n=59) ^18^, with no samples being excluded from these datasets. The signature of selection was examined using polarized and unpolarized iHS tests implemented in REHH R package ^61^, Tajima’s D of VCF-kit v. 0.2.9 ^62^, and nucleotide diversity of VCFtools v.0.1.17 ^63^. For the polarized iHS test, the SNVs were polarized using chimpanzee genome ^22^ and filtered for biallelic SNVs. For identifying the candidate region, the proportion of SNVs with |iHS| > 2 were calculated across a 20 kbp window genome-wide. The top 5% windows with highest proportion of |iHS| > 2 SNVs were considered as a signature of selection. Tajima’s D and nucleotide diversity were calculated across the 20 kbp window with a sliding step of 10 kbp.

### Inversion breakpoint mapping

To narrowly map the inversion breakpoints within flanking inverted repeats, we used two different methods depending on whichever worked best for different inversions (Inv-γ or Inv-γ-del). The first method uses breakpoint mapping using an MSA of inverted repeats from inverted and direct reference haplotype ^32^. The other method relies on mapping 201-mers from the direct reference haplotype on top of the inverted haplotype both in direct and reverse orientation. For both inversions we started by selecting the best direct reference haplotype based on the fraction of shared 31-mers between inverted and direct haplotype. The pool of possible direct haplotypes for Inv-γ-del was selected from structural haplotype group 5 (n=33, Figure 4A). In the case of Inv-γ, the pool of possible direct haplotypes was based on structural haplotype group 1 (n=275, Figure 4A). To map the inversion breakpoints of Inv-γ-del (structural haplotype group 7; n=20), we chose the MSA method. In this method we start by extracting a DNA sequence corresponding to the FAM7A and GOLGA8 HOR from both inverted and direct reference haplotype (the best matching one as described above). We reverse complement the distal copies of inverted repeats in both inverted and direct haplotype. Then, we construct the MSA using the R package DECIPHER. Next, we extracted paralog-specific variants (PSVs) and defined the inversion breakpoint as a region where inverted haplotypes partly match proximal and distal PSVs of the direct haplotype; for more details, see ^12^. To map the inversion breakpoints of Inv-γ (structural haplotype group 3; n=36), we chose the k-mer method because it was more robust to extended regions of homozygosity within the SDα repeats. In this method we split the selected direct reference haplotype (the best matching one as described above) into overlapping 201-mers and removed any repetitive k-mers. We then look for exact matches of reference k-mers within an inverted haplotype both in direct and reverse orientation. We kept only those k-mers that had a single exact match either in direct or reverse orientation. Then, we look for a changepoint where the majority of direct matching k-mers switch to the majority of reverse matching k-mers as putative inversion breakpoints. To find this position we walk through these k-mers in a sliding window defined as an upstream and downstream window encompassing 2% of the total number of matched k-mers. For each subsequent pair of upstream and downstream windows, we calculate the absolute difference between counts of direct and reverse mapping k-mers. We define the putative inversion breakpoint as a position where the difference between upstream and downstream window is the largest in both proximal and distal inverted repeat. We observed that inversion breakpoint is very sensitive to a chosen reference direct haplotype. Because of this we refined the chosen reference haplotype for 5 out of 36 inverted haplotypes (Supplementary Data 15).

### *PRDM9* groups

To determine *PRDM9* haplotypes, we first extracted five sequencing reads spanning the zinc-finger (ZF) array region from both haplotypes. Full sequences of the *PRDM9* alleles described by Alleva et al. ^64^ were aligned to these reads using minimap2 (version 2.30-r1287) ^43^. For each read, the best-matching allele was defined as the *PRDM9* allele with (i) an alignment length within ±10 bp of the allele length and (ii) the lowest number of mismatches (NM) to the sequencing read. Finally, we reported the *PRDM9* allele matched by most out of the five sequencing reads to alleviate sequencing errors that may affect calls made by individual reads.

### *PRDM9* motif identification

We identified *PRDM9* motif occurrences using FIMO v5.5.8 ^65^ with the JASPAR ^66^ position weight matrices (PWMs) MA1723.1 and MA1723.2. Scans were run on the T2T-CHM13 chromosome 15 sequence using default background parameters. FIMO reports a PWM log-odds match score, which we used to quantify motif strength within the analyzed interval.

## Data Availability

Genome assemblies generated by the HGSVC and HPRC used in this study are available under NCBI BioProject IDs PRJEB83624 (HGSVC) and PRJNA730822 and PRJNA730823 (HPRC releases 1 and 2, respectively). Non-human primate genome assemblies used in this study are available at NCBI under accessions GCF_028858775.2, GCF_029281585.2, GCF_028885625.2, GCF_028878055.2, GCF_028885655.2, GCF_029289425.2 and GCF_037993035.1. The raw sequencing data for the ten clinical patient trios generated in this study have been deposited in the European Genome-phenome Archive (EGA) under accession code EGAD50000002913. These data are available under restricted access to protect the privacy of clinical study participants; access can be obtained by submitting a formal application via the EGA website, which is reviewed by the Radboudumc Data Access Committee (DAC) and, if approved, governed by a Data Access Agreement (DAA) defining the terms of use. Access is granted to researchers for non-commercial academic use. Requests and enquiries should be made through the EGA portal. Local assemblies and assembly visualisations restricted to the 15q13.3 region have been deposited under restricted access at Zenodo (10.5281/zenodo.19571509) and are released by the record owner under the same conditions. All remaining data supporting the findings are provided in the Supplementary Data and the Source Data files.

## Code Availability

Software generated for this study is available at GitHub under the following accessions Multiple sequence alignments for breakpoint inference:

https://github.com/WHops/breakpoint_msa

Crosshair for unitig-based breakpoint inference:

https://github.com/WHops/crosshair.git

Assembly contig merging for “duo-based” phasing:

https://github.com/WHops/contig-merger.git

RHA and allelic outcompetition:

https://github.com/WHops/SD_outcompetition

Assembly-based variant calling and phylogenetic clustering:

https://github.com/WHops/asm_fastas_to_snps

Custom mapping of OGM reads to a custom CNV-specific 300kbp sequence:

https://github.com/WHops/OGM-junction-tester

## Acknowledgements

We thank all patients and their families for participation in this study. We thank members of the HGSVC for providing access to whole-genome assemblies used in our study. We thank Tonia Brown for edits in the preparation of this manuscript. We thank the Radboud Technology Center Genomics and Ronald van Beek for help with sequencing and optical genome mapping.

## Funding Statement

This project has received funding from the European Union’s Horizon 2020 research and innovation programme under the Marie Skłodowska-Curie grant agreement no. 101150006 (to W.H.) and from the Dutch Organisation for knowledge and innovation in health, healthcare and well-being (ZonMw) and the European Joint Programme on Rare Diseases (EJP RD): dossier nr. 04630042210002. This project has further received funding from the Federal Ministry of Research, Technology and Space (BMFTR), partner of the EJP RD, funding reference no. 01GM2307 (to C.P.S.). The EJP RD initiative has received funding from the European Union’s Horizon 2020 research and innovation programme under grant agreement No. 825575. C.P.S. was supported by a Marsilius Fellowship of Heidelberg University. Research reported in this publication was supported, in part, by the National Human Genome Research Institute of the National Institutes of Health (NIH) under award numbers R01HG002385 and R01HG010169 (to E.E.E.). The content is solely the responsibility of the authors and does not necessarily represent the official views of the NIH. E.E.E. is an investigator of the Howard Hughes Medical Institute. Several of the authors of this publication are members of the European Reference Network on Rare Congenital Malformations and Rare Intellectual Disability ERN-ITHACA. This study was generated in part within the European Reference Network on Rare Congenital Malformations and Rare Intellectual Disability (ERN-ITHACA) (grant agreement N°101156387). This article is subject to HHMI’s Open Access to Publications policy. HHMI lab heads have previously granted a nonexclusive CC BY 4.0 license to the public and a sublicensable license to HHMI in their research articles. Pursuant to those licenses, the author-accepted manuscript of this article can be made freely available under a CC BY 4.0 license immediately upon publication.

### Human Pangenome Reference Consortium: Funding & Acknowledgements

We would like to acknowledge the National Human Genome Research Institute (NHGRI) for funding the following grants supporting the creation of the human pangenome reference: U41HG010972, U01HG010971, U01HG013760, U01HG013755, U01HG013748, U01HG013744, R01HG011274, and the Human Pangenome Reference Consortium (BioProject ID: PRJNA730823). This research was supported in part by the Intramural Research Program of the National Institutes of Health (NIH). The contributions of the NIH author(s) are considered Works of the United States Government. The findings and conclusions presented in this paper are those of the author(s) and do not necessarily reflect the views of the NIH or the U.S. Department of Health and Human Services. This work utilized the computational resources of the NIH HPC Biowulf cluster (https://hpc.nih.gov).

## Author Contributions

Conceptualization: W.H., C.G., D.P., D.Y., E.E.E., C.P.S.

Methodology & software: W.H., D.P., D.Y., M.d.G.

Sequencing & Patient Contact: A.d.O., R.D., K.N., A.H., H.G.Y., K.H., P.C., A.d.F., N.B.-P., B.v.B.

Data resource: HPRC.

Formal analysis: W.H., D.P., D.Y., M.d.G.

Validation: W.H., C.G., D.P., E.E.E.

Writing: W.H., C.G., D.P., D.Y., E.E.E., with input from all authors.

## Competing Interests

E.E.E. is a scientific advisory board (SAB) member of Variant Bio, Inc. All other authors declare no competing interests.

### Human Pangenome Reference Consortium Version 2 Authors

Derek Albracht^9^, Ivan A. Alexandrov^10^, Jamie Allen^11^, Alawi A. Alsheikh-Ali^12^, Nicolas Altemose^13^, Casey Andrews^14^, Dmitry Antipov^15^, Lucinda Antonacci-Fulton^9^, Mobin Asri^16^, Marcelo Ayllon^2^, Jennifer R. Balacco^17^, Floris P. Barthel^18^, Edward A. Belter Jr^9^, Halle D. Bender^16^, Andrew P. Blair^16^, Davide Bolognini^19^, Katherine E. Bonini^20^, Christina Boucher^21^, Guillaume Bourque^22,23,24^, Silvia Buonaiuto^25^, Shuo Cao^25^, Andrew Carroll^26^, Ann M. Mc Cartney^16^, Monika Cechova^16^, Mark J.P. Chaisson^27^, Pi-Chuan Chang^26^, Xian Chang^16^, Jitender Cheema^11^, Haoyu Cheng^28^, Claudio Ciofi^29^, Hiram Clawson^16^, Sarah Cody^9^, Vincenza Colonna^25^, Holland C. Conwell^30^, Robert Cook-Deegan^31^, Mark Diekhans^16^, Maria Angela Diroma^29^, Daniel Doerr^32,33,34^, Zheng Dong^14^, Danilo Dubocanin^13^, Richard Durbin^35,36^, Jana Ebler^32,37^, Evan E. Eichler^2,8^, Jordan M. Eizenga^16^, Parsa Eskandar^16^, Eddie Ferro^21^, Anna-Sophie Fiston-Lavier^39,40^, Sarah M. Ford^30^, Willard W. Ford^41^, Giulio Formenti^17^, Adam Frankish^11^, Mallory A. Freeberg^11^, Qichen Fu^14^, Stephanie M. Fullerton^42^, Robert S. Fulton^9^, Yan Gao^43^, Gage H. Garcia^2^, Obed A. Garcia^44^, Joshua M.V. Gardner^16^, Shilpa Garg^45^, Erik Garrison^25^, Nanibaa’ A. Garrison^46,47,48^, John E. Garza^9^, Margarita Geleta^49^, Mohammadmersad Ghorbani^50^, Tina A. Graves-Lindsay^9^, Richard E. Green^30^, Cristian Groza^51^, Bida Gu^27^, Andrea Guarracino^18,25^, Melissa Gymrek^52^, Maximilian Haeussler^16^, Leanne Haggerty^11^, Ira M. Hall^53,54^, Nancy F. Hansen^15^, Yue Hao^18^, Mohammad Amiruddin Hashmi^12^, David Haussler^16^, Prajna Hebbar^16^, Peter Heringer^32,33,34^, Glenn Hickey^16^, Todd L. Hillaker^16^, S. Nakib Hossain^11^, Neng Huang^43,55^, Sarah E. Hunt^11^, Toby Hunt^11^, Alexander G. Ioannidis^13,16^, Nafiseh Jafarzadeh^16^, Nivesh Jain^17^, Erich D. Jarvis^17,38^, Maryam Jehangir^18^, Juan Jiang^14^, Eimear E. Kenny^20^, Juhyun Kim^15^, Bonhwang Koo^17^, Sergey Koren^15^, Milinn Kremitzki^9,14^, Charles H. Langley^56^, Ben Langmead^57^, Heather A. Lawson^14^, Daofeng Li^14^, Heng Li^43,55^, Wen-Wei Liao^53,54^, Jiadong Lin^2^, Tianjie Liu^14^, Glennis A. Logsdon^58^, Ryan Lorig-Roach^16^, Jonathan LoTempio Jr^59^, Hailey Loucks^16^, Jane E. Loveland^11^, Jianguo Lu^60^, Shuangjia Lu^53,54^, Julian K. Lucas^16^, Walfred Ma^27^, Juan F. Macias-Velasco^9,14,61^, Kateryna D. Makova^62^, Maximillian G. Marin^43,55^, Christopher Markovic^9^, Tobias Marschall^32,37^, Franco L. Marsico^25^, Fergal J. Martin^11^, Mira Mastoras^16^, Capucine Mayoud^39^, Brandy McNulty^16^, Jack A. Medico^17^, Julian M. Menendez^16^, Karen H. Miga^16^, Anna Minkina^63^, Matthew W. Mitchell^64^, Saswat K. Mohanty^65^, Younes Mokrab^50,66,67^, Jean Monlong^68^, Shabir Moosa^50^, Avelina Moreno-Ochando^69,70^, Shinichi Morishita^71^, Jonathan M. Mudge^11^, Katherine M. Munson^2^, Njagi Mwaniki^72^, Nasna Nassir^12^, Chiara Natali^29^, Shloka Negi^16^, Lingbin Ni^2^, Adam M. Novak^16^, Pilar N. Ossorio^73^, Chie Owa^71^, Sadye Paez^17^, Benedict Paten^16^, Clelia Peano^19,74^, Adam M. Phillippy^15^, Brandon D. Pickett^15^, Laura Pignata^25^, Nadia Pisanti^72^, David Porubsky^2,3^, Pjotr Prins^25^, Anandi Radhakrishnan^16^, T. Rhyker Ranallo-Benavidez^18^, Brian J. Raney^16^, Mikko Rautiainen^75^, Alessandro Raveane^19^, Luyao Ren^2,38^, Arang Rhie^15^, Fedor Ryabov^76,77^, Samuel Sacco^30^, Farnaz Salehi^25^, Michael C. Schatz^57,78^, Laura B. Scheinfeldt^64^, Aarushi Sehgal^41^, William E. Seligmann^30^, Mahsa Shabani^79^, Kishwar Shafin^26^, Shadi Shahatit^39^, Ruhollah Shemirani^20^, Vikram S. Shivakumar^57^, Swati Sinha^11^, Jouni Sirén^16^, Linnéa Smeds^65^, Steven J. Solar^15^, Marco Sollitto^17,29^, Nicole Soranzo^19,35,80^, Andrew B. Stergachis^2,63^, Marie-Marthe Suner^11^, Yoshihiko Suzuki^71^, Arda Söylev^32,37^, Ahmad Abou Tayoun^81,82^, Jack A.S. Tierney^11^, Chad Tomlinson^9^, Francesca Floriana Tricomi^11^, Mohammed Uddin^12,83^, Matteo Tommaso Ungaro^30,84^, Rahul Varki^21^, Flavia Villani^25^, Ivo Violich^16^, Mitchell R. Vollger^63^, Brian P. Walenz^15^, Charles Wang^85^, Lisa E. Wang^20^, Ting Wang^9,14,61^, Aaron M. Wenger^86^, Conor V. Whelan^17^, Zilan Xin^14^, Zheng Xu^14^, Kai Ye^87^, DongAhn Yoo^2^, Wenjin Zhang^14^, Ying Zhou^43^, Xiaoyu Zhuo^14^, Giulia Zunino^19^

### Affiliations

1 Department of Human Genetics, Radboud University Medical Center, Nijmegen, The Netherlands

2 Department of Genome Sciences, University of Washington School of Medicine, Seattle, WA, USA

3 European Molecular Biology Laboratory (EMBL), Genome Biology Unit, Heidelberg, Germany

4 Institute of Human Genetics, Heidelberg University and Heidelberg University Hospital, Heidelberg, Germany

5 Telethon Institute of Genetics and Medicine (TIGEM), Pozzuoli, Italy

6 Department of Translational Medical Sciences, University of Naples Federico II, Naples, Italy

7 Genomics and Experimental Medicine Program, Scuola Superiore Meridionale (SSM, School of Advanced Studies), Naples, Italy.

8 Howard Hughes Medical Institute, University of Washington, Seattle, WA, USA

9 McDonnell Genome Institute, Washington University School of Medicine, St. Louis, MO 63108, USA

10 Department of Human Molecular Genetics and Biochemistry, Faculty of Medical and Health Sciences, Tel Aviv University, Tel Aviv 69978, Israel

11 European Molecular Biology Laboratory, European Bioinformatics Institute (EMBL-EBI), Wellcome Genome Campus, Hinxton, Cambridge CB10 1SD, UK

12 Center for Applied and Translational Genomics (CATG), Mohammed Bin Rashid University of Medicine and Health Sciences, Dubai Health, Dubai, UAE

13 Department of Genetics, Stanford University, Palo Alto, CA 94304 USA

14 Department of Genetics, Washington University School of Medicine, St. Louis, MO 63110, USA

15 Genome Informatics Section, Center for Genomics and Data Science Research, National Human Genome Research Institute, National Institutes of Health, Bethesda, MD 20892, USA

16 UC Santa Cruz Genomics Institute, University of California, Santa Cruz, CA 95060, USA

17 The Vertebrate Genome Laboratory, The Rockefeller University, New York, NY 10065, USA

18 Bioinnovation and Genome Sciences, The Translational Genomics Research Institute (TGen), Phoenix, AZ 85004, USA

19 Human Technopole, Milan, Italy

20 Institute for Genomic Health, Icahn School of Medicine at Mount Sinai, New York, NY 10029, USA

21 Department of Computer and Information Science and Engineering, University of Florida, Gainesville, FL 32611, USA

22 Canadian Center for Computational Genomics, McGill University, Montréal, QC H3A 0G1, Canada

23 Department of Human Genetics, McGill University, Montréal, QC H3A 0G1, Canada

24 Victor Phillip Dahdaleh Institute of Genomic Medicine, Montréal, QC H3A 0G1, Canada

25 Department of Genetics, Genomics and Informatics, University of Tennessee Health Science Center, Memphis, TN 38163, USA

26 Google LLC, Mountain View, CA 94043, USA

27 Quantitative and Computational Biology, University of Southern California, Los Angeles, CA 90089, USA

28 Department of Biomedical Informatics and Data Science, Yale School of Medicine, New Haven, CT 06510, USA

29 Department of Biology, University of Florence, Sesto Fiorentino, FI 50019, Italy

30 Department of Ecology and Evolutionary Biology, University of California, Santa Cruz, CA 95060, USA

31 Arizona State University, Consortium for Science, Policy & Outcomes, Washington, DC 20006, USA

32 Center for Digital Medicine, Heinrich Heine University Düsseldorf, Düsseldorf, NRW, DE

33 Department for Endocrinology and Diabetology at the Medical Faculty and University Hospital Düsseldorf, Heinrich Heine University Düsseldorf, Düsseldorf, NRW, DE

34 Paul-Langerhans-Group Computational Diabetology, German Diabetes Center (DDZ) and Leibniz Institute for Diabetes Research, Düsseldorf, NRW, DE

35 Wellcome Sanger Institute, Genome Campus, Hinxton, CB10 1RQ, UK

36 Department of Genetics, University of Cambridge, Cambridge, CB2 3EH, UK

37 Institute for Medical Biometry and Bioinformatics, Medical Faculty and University Hospital Düsseldorf, Heinrich Heine University, Düsseldorf, NRW, DE

38 Howard Hughes Medical Institute, Chevy Chase, MD 20815, USA

39 ISEM, Univ Montpellier, CNRS, IRD, Montpellier, FR

40 Institut Universitaire de France, Paris, FR

41 Department of Computer Science and Engineering, University of California San Diego, La Jolla, CA 92093, USA

42 Department of Bioethics & Humanities, University of Washington School of Medicine, Seattle, WA 98195, USA

43 Department of Data Science, Dana-Farber Cancer Institute, Boston, MA 02215, USA

44 Department of Anthropology, University of Kansas, Lawrence, KS 66045, USA

45 School of Health Sciences, University of Manchester, Manchester M13 9PL, UK

46 Traditional, ancestral and unceded territory of the Gabrielino/Tongva peoples, Institute for Society & Genetics, University of California, Los Angeles, Los Angeles, CA 90095, USA

47 Traditional, ancestral and unceded territory of the Gabrielino/Tongva peoples, Institute for Precision Health, David Geffen School of Medicine, University of California, Los Angeles, Los Angeles, CA 90095, USA

48 Traditional, ancestral and unceded territory of the Gabrielino/Tongva peoples, Division of General Internal Medicine & Health Services Research, David Geffen School of Medicine, University of California, Los Angeles, Los Angeles, CA 90095, USA

49 Department of Electrical Engineering and Computer Science, University of California, Berkeley, Berkeley, CA 94720, USA

50 Medical and Population Genomics Lab, Sidra Medicine, Doha, Qatar

51 Montreal Heart Institute, Montréal, QC, Canada

52 Department of Pediatrics, University of California San Diego, La Jolla, CA 92093, USA

53 Center for Genomic Health, Yale University School of Medicine, New Haven, CT 06510, USA

54 Department of Genetics, Yale University School of Medicine, New Haven, CT 06510, USA

55 Department of Biomedical Informatics, Harvard Medical School, Boston, MA 02115, USA

56 Department of Evolution and Ecology and the Center for Population Biology, University of California, One Shields, Davis, CA 95616, USA

57 Department of Computer Science, Johns Hopkins University, Baltimore, MD 21218, USA

58 Department of Genetics, Epigenetics Institute, Perelman School of Medicine, University of Pennsylvania, Philadelphia, PA 19104, USA

59 Department of Pediatrics, Division of Genetics, School of Medicine, University of California, Irvine, CA 92697, USA

60 Sun Yat-sen University, Guangzhou, China

61 Edison Family Center for Genome Sciences & Systems Biology, Washington University School of Medicine, St. Louis, MO 63110, USA

62 Department of Biology and Center for Medical Genomics, Penn State University, University Park, PA 16802, USA

63 Division of Medical Genetics, Department of Medicine, University of Washington School of Medicine, Seattle, WA 98195, USA

64 Coriell Institute for Medical Research, Camden, NJ 08103, USA

65 Department of Biology, Penn State University, University Park, PA 16802, USA

66 Department of Biomedical Science, College of Health Sciences, Qatar University, Doha, Qatar

67 Department of Genetic Medicine, Weill Cornell Medicine-Qatar, Doha, Qatar

68 IRSD - Digestive Health Research Institute, University of Toulouse, INSERM, INRAE, ENVT, UPS, Toulouse, FR

69 MATCH biosystems, S.L., Elche, Spain

70 Universidad Miguel Hernández de Elche, Elche, Spain

71 Department of Computational Biology and Medical Sciences, The University of Tokyo, Kashiwa, Chiba 277-8561, Japan

72 Department of Computer Science, University of Pisa, Pisa, Italy

73 Law School, University of Wisconsin-Madison, Madison, WI 53706, USA

74 Institute of Genetics and Biomedical Research, UoS of Milan, National Research Council, Milan, Italy

75 Institute for Molecular Medicine Finland, Helsinki Institute of Life Science, University of Helsinki, Helsinki, Finland

76 The Center for Bio- and Medical Technologies, Moscow, RUS

77 Centre for Biomedical Research and Technology, HSE University, Moscow, RUS

78 Department of Biology, Johns Hopkins University, Baltimore, MD 21218, USA

79 University of Amsterdam, Amsterdam, Netherlands

80 School of Clinical Medicine, University of Cambridge, Cambridge, CB2 0SP, UK

81 Center for Genomic Discovery, Mohammed Bin Rashid University, Dubai Health, UAE

82 Dubai Health Genomic Medicine Center, Dubai Health, UAE

83 GenomeArc Inc, Mississauga, ON, Canada

84 Department of Biology and Biotechnologies “Charles Darwin”, University of Rome “La Sapienza”, Rome 00185, IT

85 Center for Genomics, Loma Linda University School of Medicine, Loma Linda, CA 92350, USA

86 PacBio, Menlo Park, CA 94025, USA

87 The first affiliated hospital of Xi’an Jiaotong University, Xi’an Jiaotong University, Xi’an, Shaanxi, 710049, China

