## Supplementary Materials for "Distinct mechanisms of CNV formation at the human 15q13.3 locus"

### Supplementary Materials for:
### Distinct mechanisms of CNV formation at the human 15q13.3 locus

#### Supplementary Notes

**Supplementary Note 1: Orthogonal validation of 8/30 samples with Optical Genome Mapping.** OGM genome maps were generated for 8 individuals from 4 families (Methods) covering the following CNV structures:

- 2x BP4-BP5 deletion (T6-index, T6-mother)
- 3x *CHRNA7* duplications (T9-index, T9-father, T10-mother)
- 1x *CHRNA7* deletion (T7-index)
- 1x Inv-δ (T7-mother)
- 1x Inv-β’ (h1) and deletion inside BP4 (h2); (T6-father)

For 7 of 8 individuals, OGM assemblies confirmed all non-reference junctions present in the LRS-based assemblies (Supplementary Data 13, Supplementary Figure 3). For T7-mother (carrying Inv-δ), no consensus OGM assembly could be generated, reflecting the fact that this sample uniquely carries an SDα-β-α configuration spanning ~790 kbp with no unique sequence between the two SDα copies. We therefore developed an OGM-assembly-free approach matching predicted OGM label patterns derived from the LRS assemblies against raw OGM molecules (Methods). This initially failed, prompting manual re-inspection of the LRS read data, which revealed a likely ~70 kbp Inv-β misassembly. This region lies approximately 464 kbp proximal to the child's CNV breakpoint and is thus not directly involved in CNV formation. The misassembly was located on the non-inherited portion of the maternal haplotype, meaning our standard parent-child cross-validation could not have detected it. After correction, the assembly-free OGM validation confirmed the Inv-δ configuration. The corrected assembly is reflected in all figures and analyses. We note that a residual possibility of local misassembly exists in non-inherited portions of unusually long segmental duplications in parents, but such regions are distal to CNV breakpoints and do not affect the structural or mechanistic conclusions of this study.

**Supplementary Note 2: Sequence similarity at the recurrent SDβ / SDβ′ NAHR breakpoint.** We investigated whether the localized recurrence of NAHR may be further nurtured by unusually high sequence similarity between SDβ and SDβ′ at this locus which may make meiotic mis-pairings more likely. We considered this both population-wide and specifically in the three parents. Contrary to expectations, we found elevated divergence between SDβ and SDβ′ in the 2-kb window around the ‘hot’ PRDM9 motif both in HPRC control genomes (3.50-fold more mismatches over the SDβ-wide median; 14 vs 4 mismatches per 2 kbp) and the three parents, albeit less pronounced in the latter (2.00-, 2.00-, and 2.75-fold enriched over median), initially indicating protection against NAHR. We also noticed, however, that SDβ′ copies among each other likewise show a divergence peak at this locus of comparable size, and particularly strongly in the parents (2.75-, 4.25-, and 3.75-fold enriched for Parents of T2, T3, and T4 over the median of HPRC samples) (Supplementary Figure 12). We did not find variants affecting the PRDM9 motif directly (Supplementary Figure 67).

#### Supplementary Data


Supplementary Data 1. Phenotypic characteristics of patients and parents
Supplementary Data 2. Sample overview and sequencing statistics
Supplementary Data 3. 15q13.3 haplotypes and mechanisms
Supplementary Data 4. CNV Breakpoint statistics
Supplementary Data 5. Number of gene copies in control and patient samples
Supplementary Data 6. Population predispositions
Supplementary Data 7. PRDM9-Alleles
Supplementary Data 8. Additional variants in patients
Supplementary Data 9. HiFi assembly statistics
Supplementary Data 10. Comparison of assembly- and read based copy numbers

Supplementary Data 11. SVs on reads-to-assembly mapping called by Sniffles
Supplementary Data 12. k-mer based confirmation of breakpoints
Supplementary Data 13. Run statistics of Optical Genome Mapping data created for 8 samples
Supplementary Data 14. Summary of duplicons used in phylogenetic tree analysis
Supplementary Data 15. Selected best direct references for each Inv-γ haplotype

### Supplementary Figures

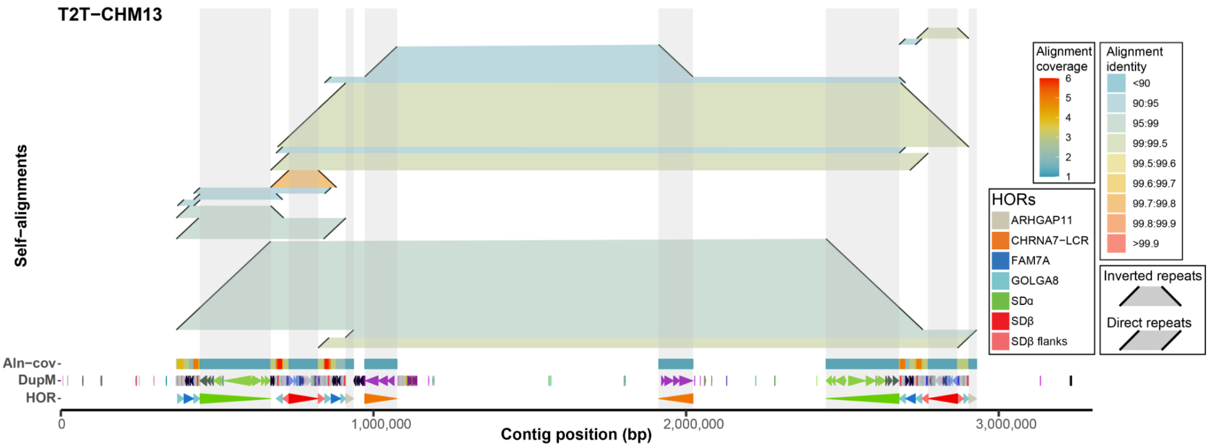


**Supplementary Figure 1. Definition of higher order structure at 15q13.2-13.3** A “horizontal dotplot” visualization showing self-alignments within the T2T-CHM13 reference at the 15q13.3 region. Alignments (diagonally oriented black lines) are connected by horizontal ribbons colored by percentage of matched bases. Below there is a heatmap summarizing coverage of overlapping self-alignments. Further below there is a DupMasker annotation of T2T-CHM13 colored by a unique duplicon IDs. Last there is a higher order repeat (HOR) annotation shown as direction pointing arrowheads.

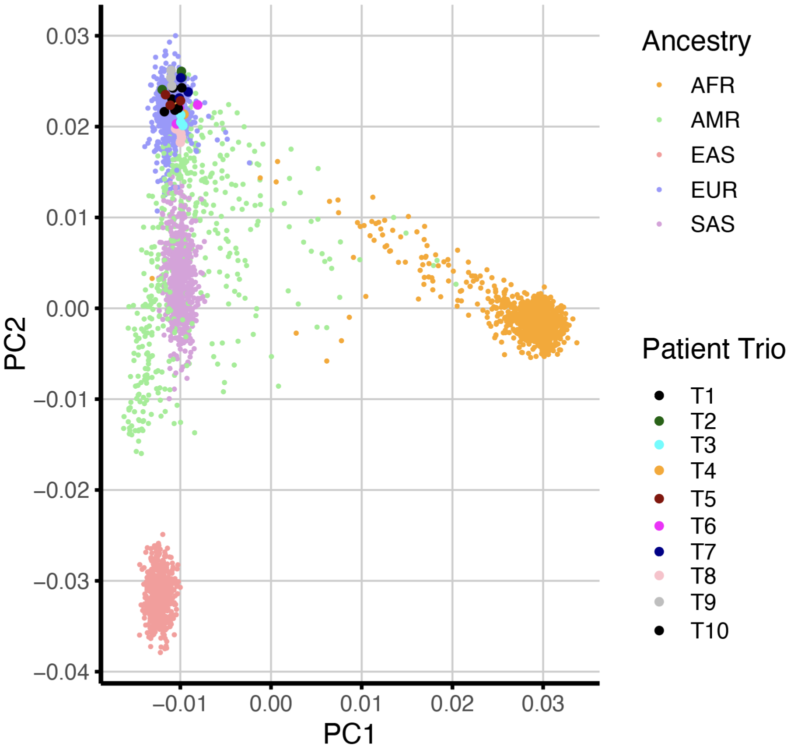

**Supplementary Figure 2**. A PCA plot of chr21 SNPs from the 1000 genomes project[1] and the 15q13.3 samples sequenced in this study. All Trios cluster closest with self-identified european individuals from the 1kGP.

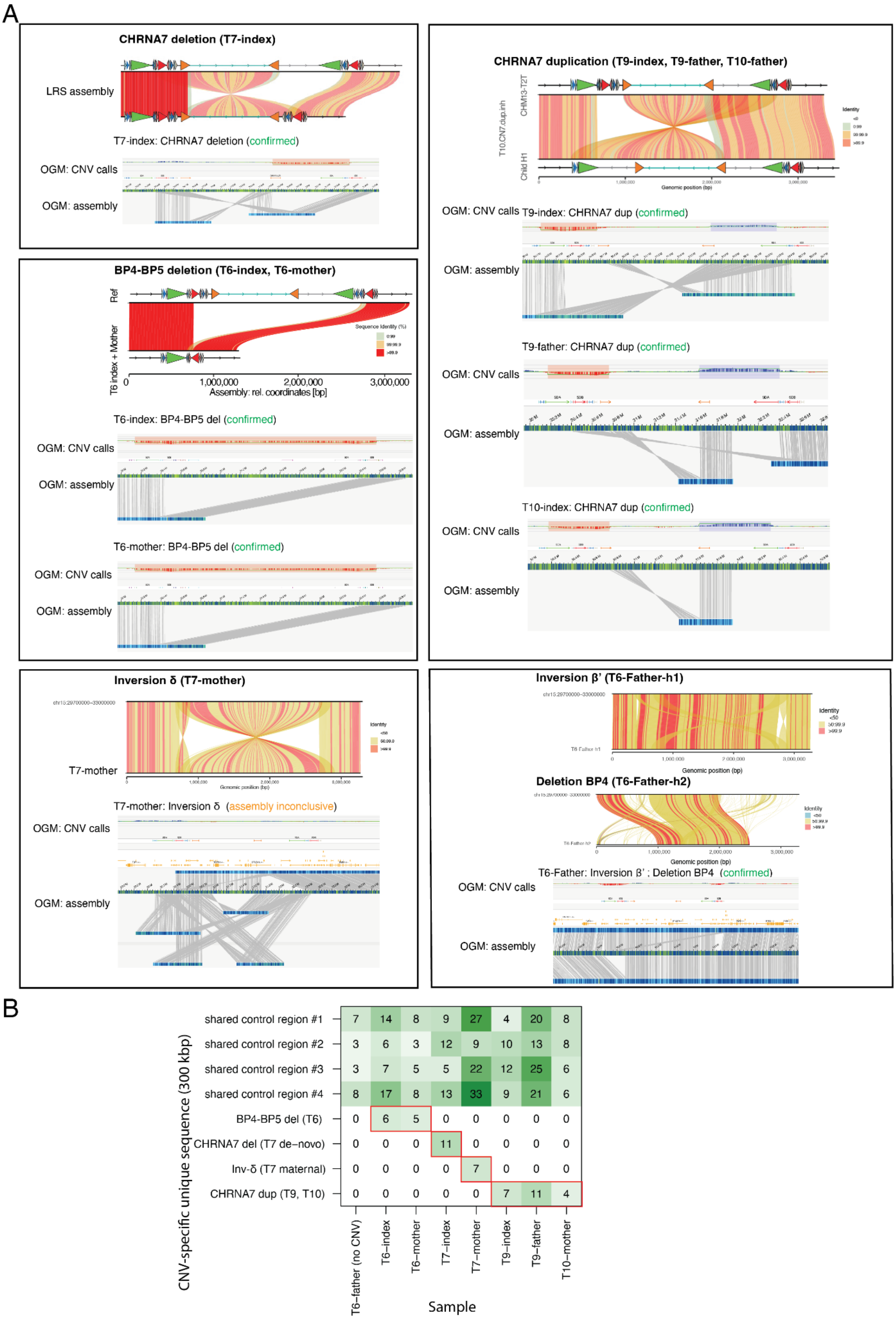

**Supplementary Figure 3. Orthogonal validation of LRS-based assemblies with Optical Genome Mapping (OGM).** A) OGM validation of CNVs in 8 individuals. For each sample, the top panel shows the LRS-based assembly aligned to GRCh38, indicating expected non-reference junctions. Below, OGM-derived CNV calls and OGM consensus assemblies are shown aligned to the same reference, independently confirming the predicted structural configuration. All tested CNVs were confirmed: *CHRNA7* deletion (T7-index), *CHRNA7* duplication (T9-index, T9-father, T10-mother), BP4-BP5 deletion (T6-index, T6-mother), Inv-β′ with BP4 deletion (T6-father). For T7-mother (Inv-δ), the OGM assembly was inconclusive due to the exceptionally long SDα-β-α configuration in this sample (~790 kbp with no unique sequence between SDα copies); this sample was validated using the assembly-free approach shown in (B). B) Assembly-free OGM validation. CNV-specific unique sequences (300 kbp) spanning structurally diagnostic junctions were extracted from LRS assemblies, converted to predicted OGM label patterns, and matched against raw OGM molecules (Methods). Numbers indicate individual molecules unambiguously spanning each CNV-specific sequence per sample. Red borders indicate expected positive cells based on the LRS assemblies. Four shared control regions present in all samples serve as positive controls. All expected CNV-specific molecules were detected exclusively in the predicted samples, including Inv-δ in T7-mother following correction of the upstream misassembly (Supplementary Note 1).

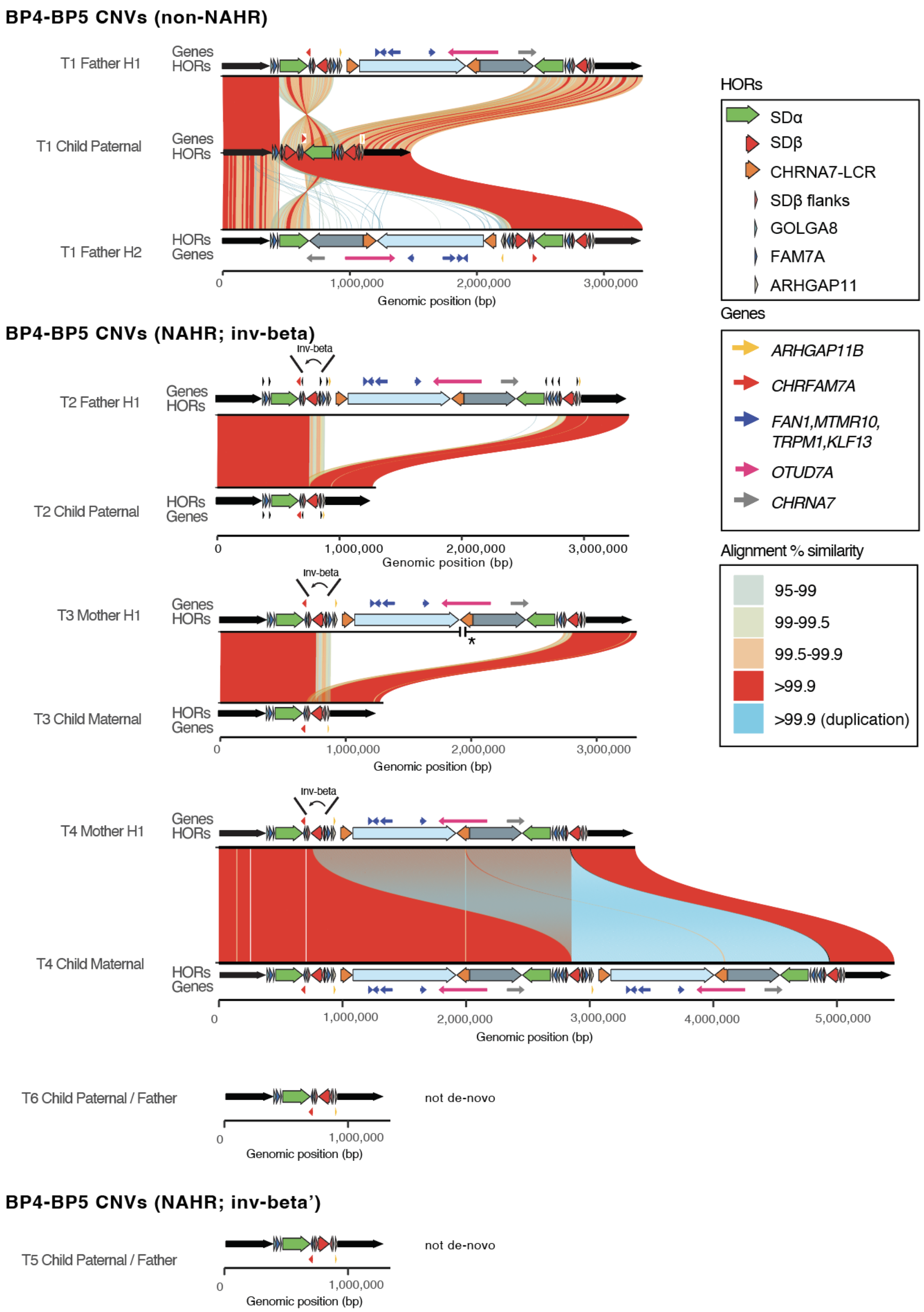


**Supplementary Figure 4. Miropeats plots showing the structure of all resolved BP4-BP5 CNVs.** HOR structures (thick arrows) and genes (thin arrows) are overlaid over all assemblies. CNV alleles which were not de novo (T5, T6) are displayed without parental alleles. Samples are grouped by structure and putative mechanism of the CNV.

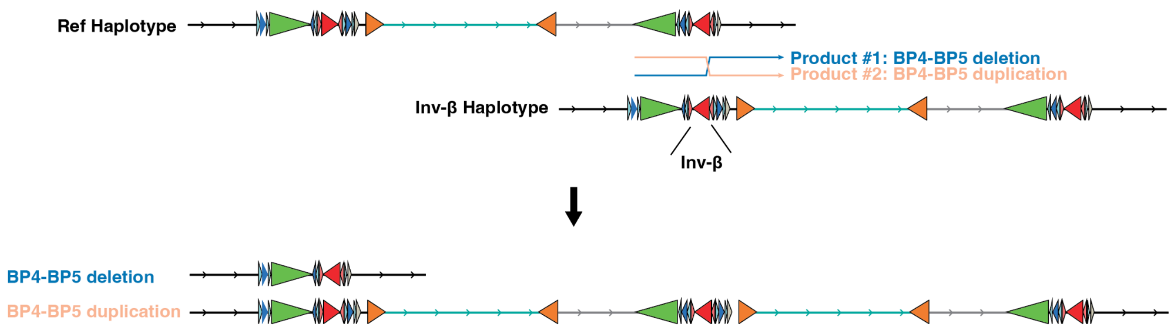


**Supplementary Figure 5. Inferred mechanism of the recurrent BP4-BP5 deletion mediated by Inv-β.** A simplified schematic of a hypothetical 15q13.3 configuration with a reference and an Inv-β carrying haplotype. Illustrated is an interchromosomal rearrangement involving the segment affected by Inv-β (SDβ; red triangles). An interchromosomal rearrangement creates a reciprocal pair of duplication and deletion. Although not illustrated here, NAHR can also appear between sister chromatids (in that case, two copies of the Inv-β haplotype would recombine, again creating a deletion/duplication pair) or within the same molecule (with the SDβ repeat pair folding back to recombine within the same molecule, creating a deletion). These mechanisms are explained in more detail e.g. in [2].

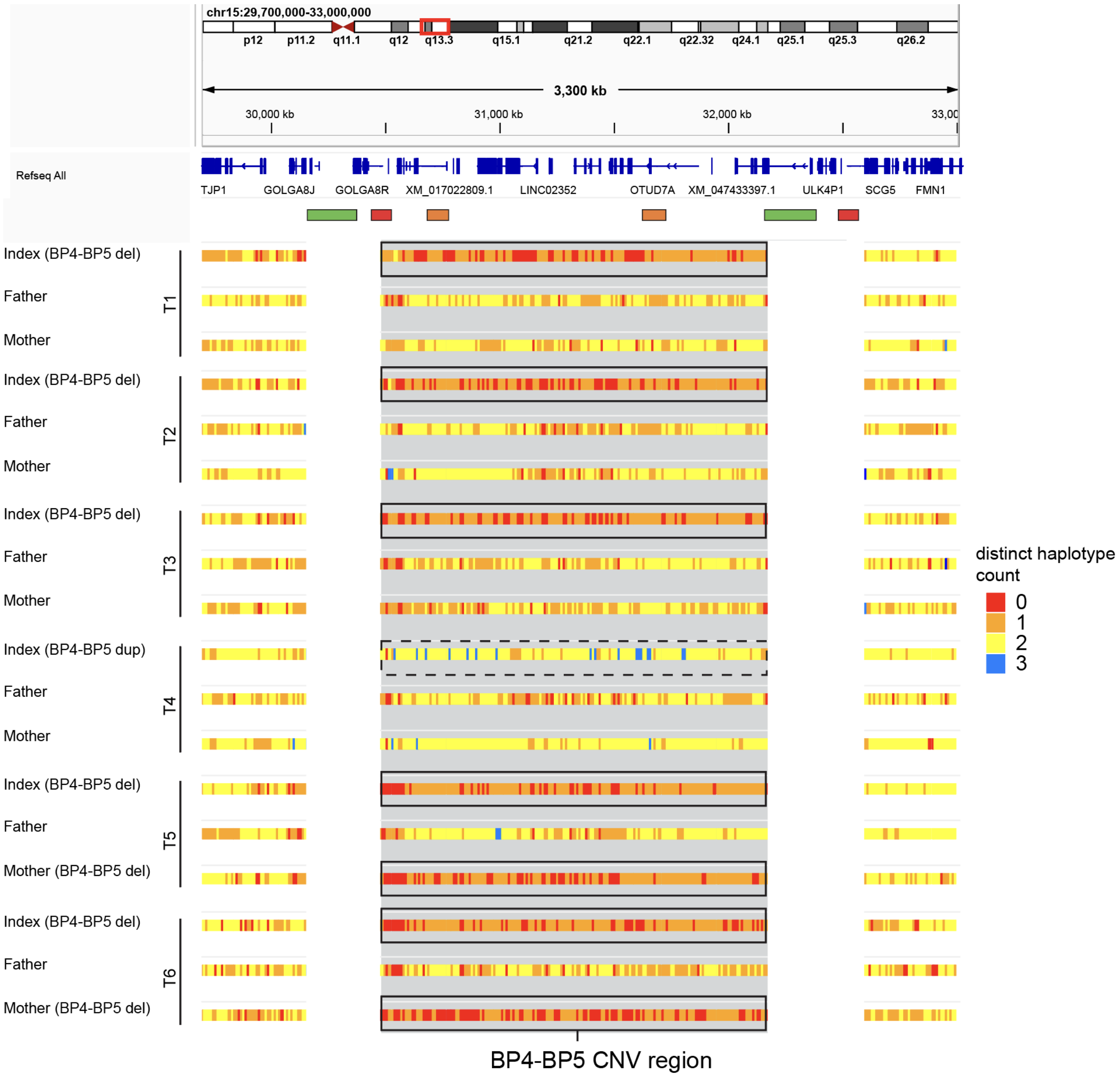


**Supplementary Figure 6. Estimation of the number of distinct alleles in BP4-BP5 Trio samples.** We independently invoked the devider program at periodic intervals of 10kbp on aligned reads of all members of BP4-BP5 CNV patient families. Devider is a long-read phasing tool which dynamically determines haplotypes per genomic position. A haplotype count '0' can either indicate that no reads were available, or that phasing was impossible, usually because no reads carried any SNVs - in that case, the haplotype count is interpreted as '1'. BP4-BP5 deletion carrying individuals are highlighted with black boxes, BP4-BP5 duplication individuals are highlighted with dotted black boxes. Aside from local phasing errors, the duplication-carrying samples carry only 2 distinct haplotypes, indicating that no inter-chromosomal duplication has taken place which would result in three distinct haplotypes. Instances of apparent 'three' haplotypes, as observed e.g. in T4-index, were manually verified to be artifacts caused by local sequencing errors or ambiguous mapping in repeat regions.

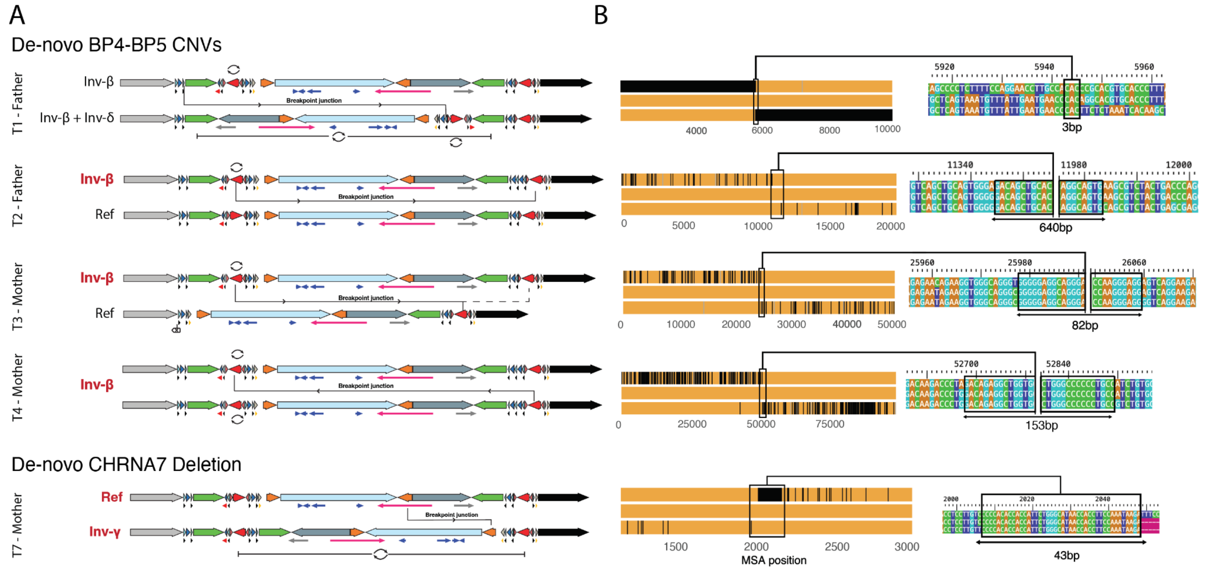


**Supplementary Figure 7. Breakpoints of de-novo 15q13.3 CNVs. A**) Parental assemblies on which de novo CNVs arose, with higher-order repeats overlaid, and inferred breakpoint positions indicated. **B)** SNP-based views of the respective breakpoints and microhomologies. Left: Three-way alignment between the parent's first breakpoint regions and the recombined sequence in the child. Mismatches are shown with thick black lines. Right: Zoom-in into the respective breakpoint regions.

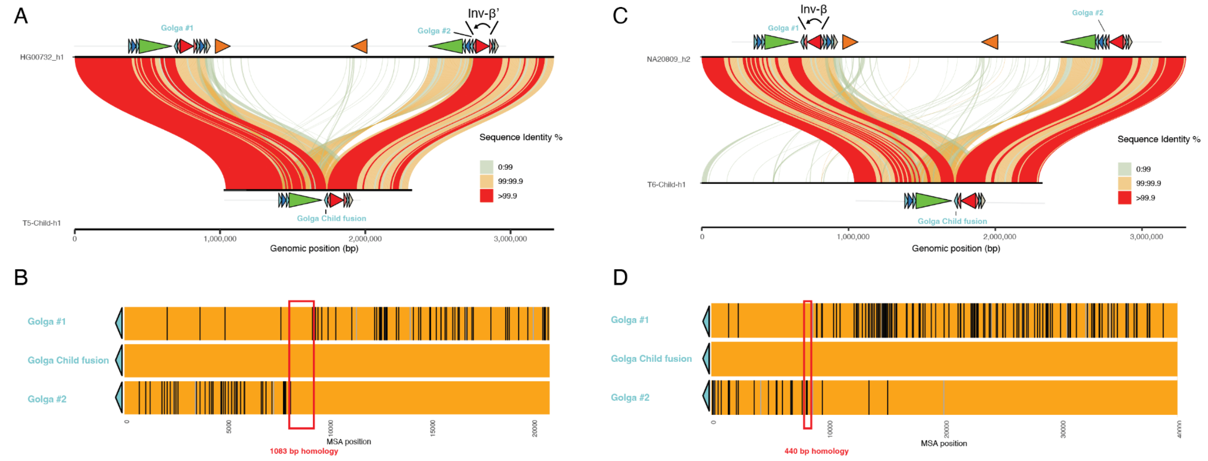


**Supplementary Figure 8. Mechanisms and structures inferred for inherited BP4-BP5 CNVs in T5 and T6.** A) An SvbyEye plot mapping the CNV in T5 to the best match from the population cohort, HG00732_h1. The latter carries an Inv- β′ which explains the predisposition to NAHR and the unique direction of SDβ in the CNV. B) A three-way alignment between two GOLGA8 elements in HG00732_h1 and the derived likely fusion in the Child (middle). A putative fusion breakpoint is visible and carries a long homology. C) A similar plot as A), this time with NA20809_h2 as the best-matching population sample to T6_child. A likely breakpoint region is visible between SDα and a GOLGA8 element D) see B).

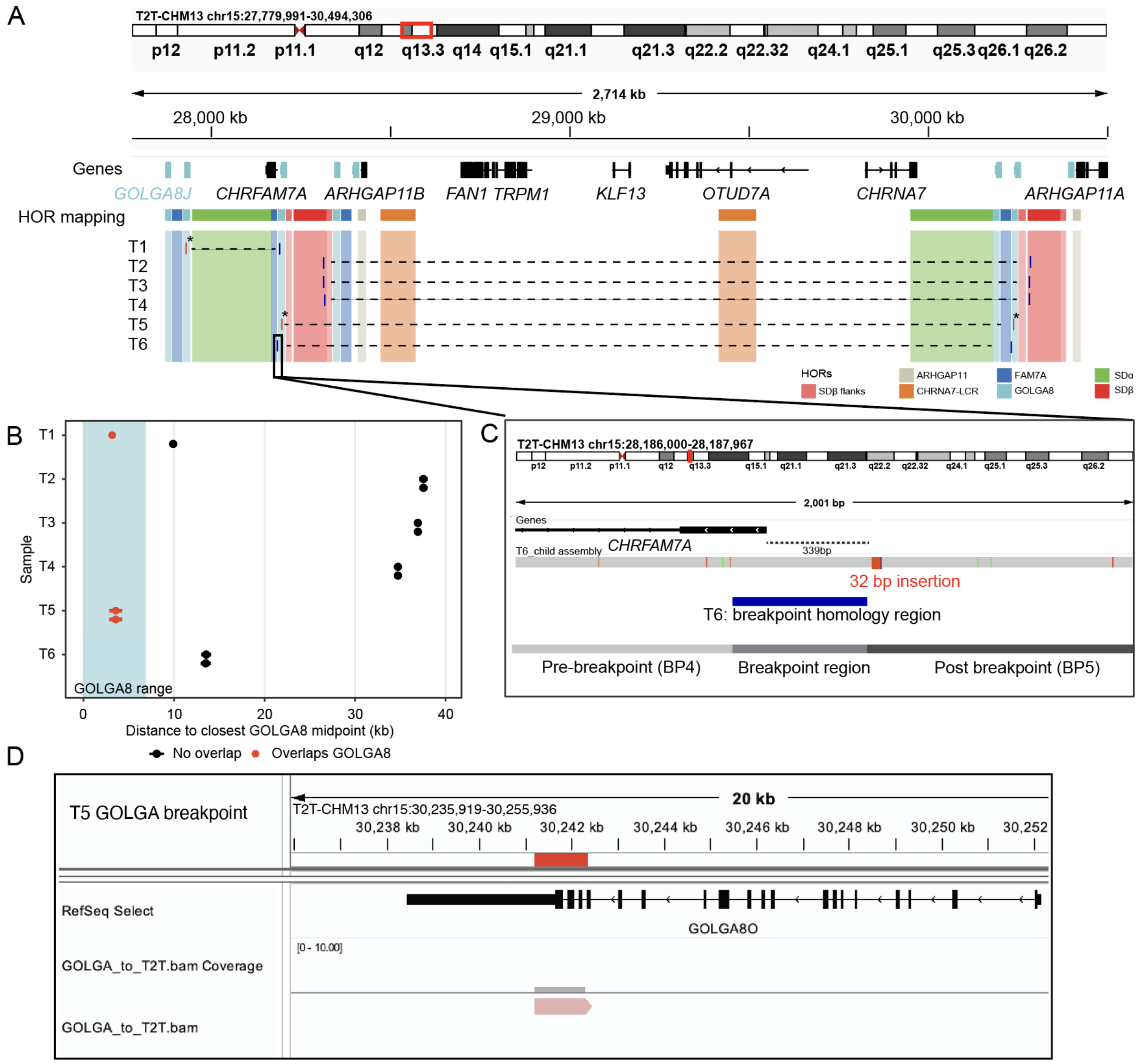


**Supplementary Figure 9. Detailed view of the breakpoint positions of BP4-BP5 CNVs in T1-T6.** A) IGV plot indicating genes and HOR annotations in the 15q13.3 region, overlaid with positions of the CNV breakpoints mapped onto T2T-CHM13. Breakpoints overlapping with a *GOLGA8* copy (T1: one breakpoint; T5: both breakpoints) are highlighted with an asterisk (*). B) Scatter plot of the distance from each breakpoint to the midpoint of the closest *GOLGA8* copy. Horizontal whiskers indicate breakpoint ambiguity arising from homology. Breakpoints overlapping with a *GOLGA8* copy are highlighted in red. C) Zoomed-in view into the breakpoint in T6. The mapped child assembly is indicated below the gene. The breakpoint homology region overlaps the first exon of *CHRFAM7A*. While the gene sequence itself remains unchanged, the region directly upstream is exchanged to a homologous partner sequence harboring e.g. a 32 bp insertion which may affect gene function. D) Visualization of the breakpoint region in T5, overlaid on the *GOLGA8O* copy in T2T-CHM13 coordinates. The break overlaps the end of the actively transcribed region.

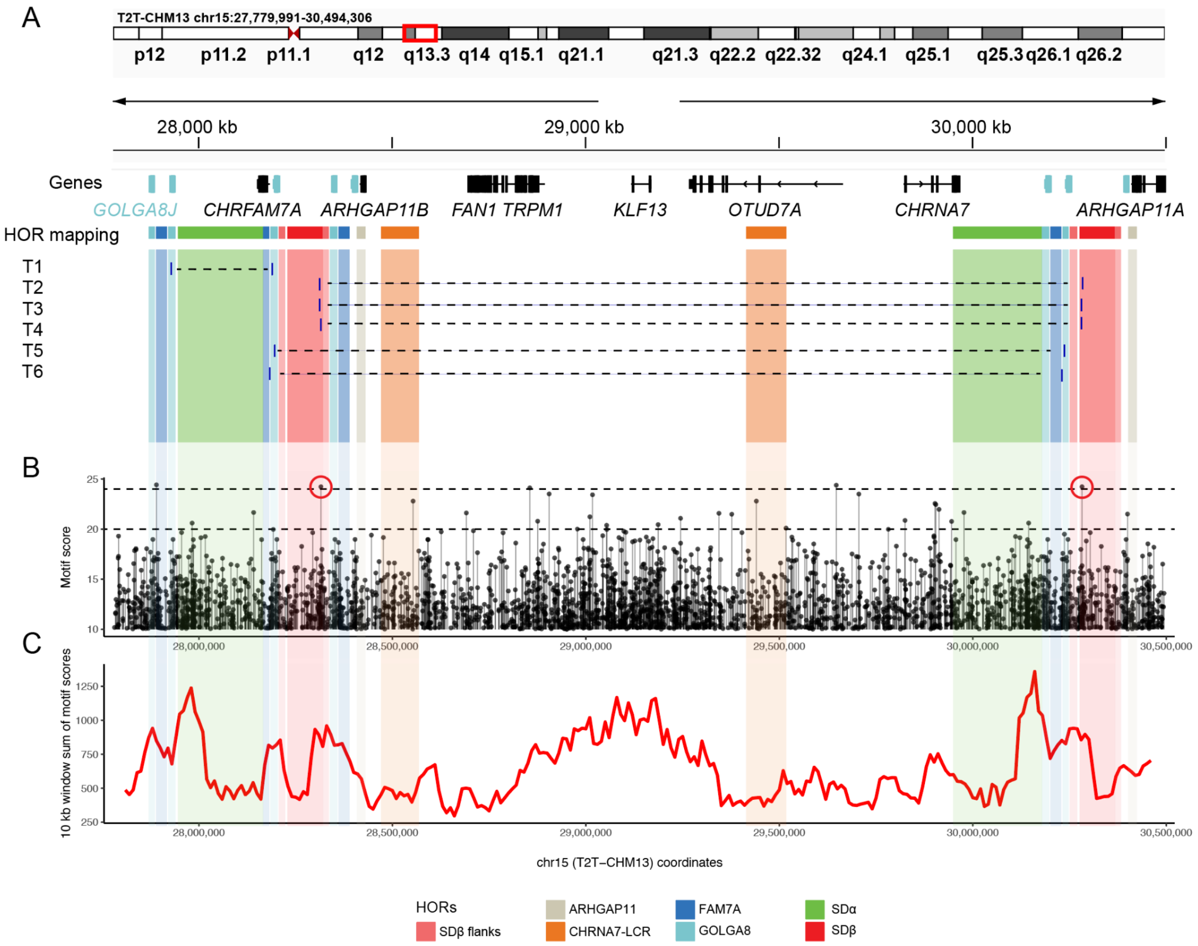


**Supplementary Figure 10. Strength and density of PRDM9 motifs in the T2T-CHM13 15q13.3 region.** A) IGV plot indicating genes and HOR annotations in the 15q13.3 region, overlaid with positions of the CNV breakpoints mapped onto T2T-CHM13. B) Strength and position of PRDM9 motif mapping score by FIMO (methods). Red circles indicate the individual strong motif associated with recurrent CNV formation in T2-T4. C) Running sum of PRDM9 mapping scores in a sliding window of 10 kbp size.

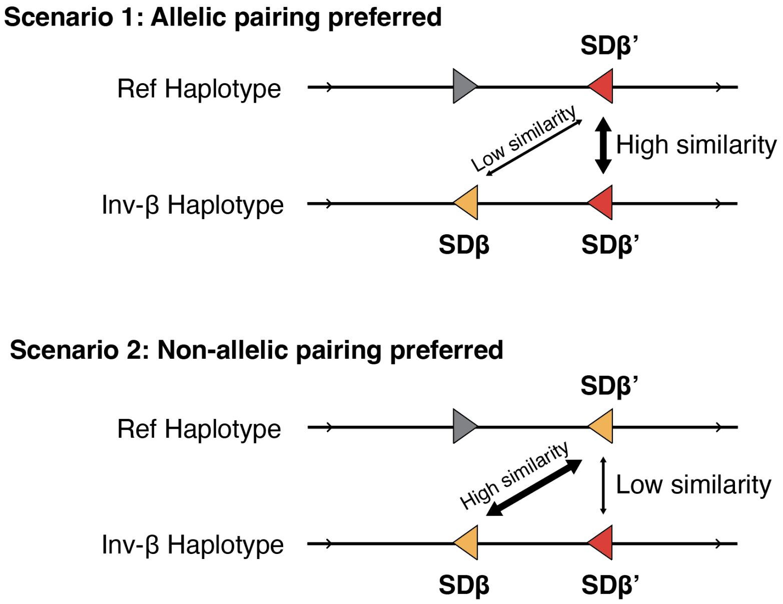

**Supplementary Figure 11. Relative Homology Advantage (RHA) principle.** Illustration of the concept of recombination competition, or relative homology advantage (RHA) between allelic, canonical homologous recombination (SDβ′ - SDβ′) and the non-allelic alternative enabled by Inv-β bringing SDβ in direct orientation with SDβ′. Top: A combination primed against NAHR may have a high similarity of the canonical pairing and a low similarity of the non-allelic pairing. Bottom: The opposite concept where SDβ′ is locally more similar to SDβ than to the allelic partner SDβ′, leading to a higher chance for mispairing. In this manuscript, we define a relative homology advantage (RHA) which quantifies the relative similarity of a non-allelic compared to an allelic pairing. If we take into account intra-chromosomal, inter-chromosomal and inter-chromatidal events, any SDβ′ instance has multiple pairing options, each with their own RHA values.

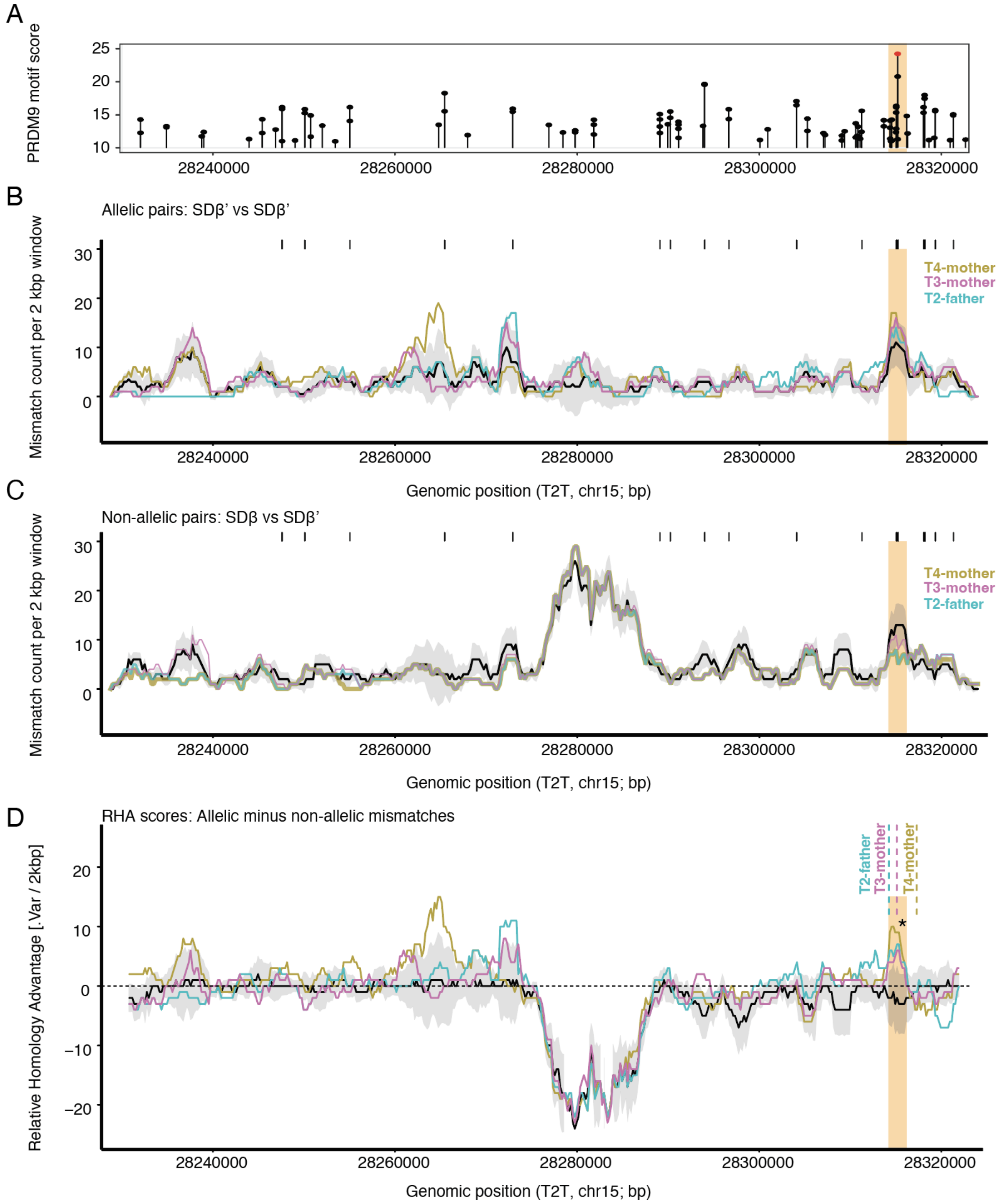


**Supplementary Figure 12. Sequence similarities along SDβ between allelic and non-allelic copies in controls and parents of de novo CNVs.** A) PRDM9 binding site scores on SDβ′ predicted by Fimo. B) 2 kbp sliding window indicating the number of variant positions between every (allelic) SDβ′-SDβ′ pair (black line / grey ribbon: median / std. dev. of all controls). The pairings that led to NAHR in the three patients are indicated in pink, blue and yellow. The 2 kbp window centered around the highest scoring PRDM9 element is indicated by an orange highlight C) Same as B, but measuring the dissimilarity between non-allelic SDβ-SDβ′ pairs. Parent samples are indicated by lines of different thickness to help distinguish overlapping tracks D) Relative Homology Advantage (RHA) scores calculated as the values from the B minus C, per pairing. A positive score indicates that for this pair and window, the non-allelic pairing has a higher similarity than the allelic pairing. For plots B-D, the black line and grey band show the median and 2.5th-97.5th percentile range across n=612 RHA scores from 153 population samples from HPRC and HGSVC (Methods).

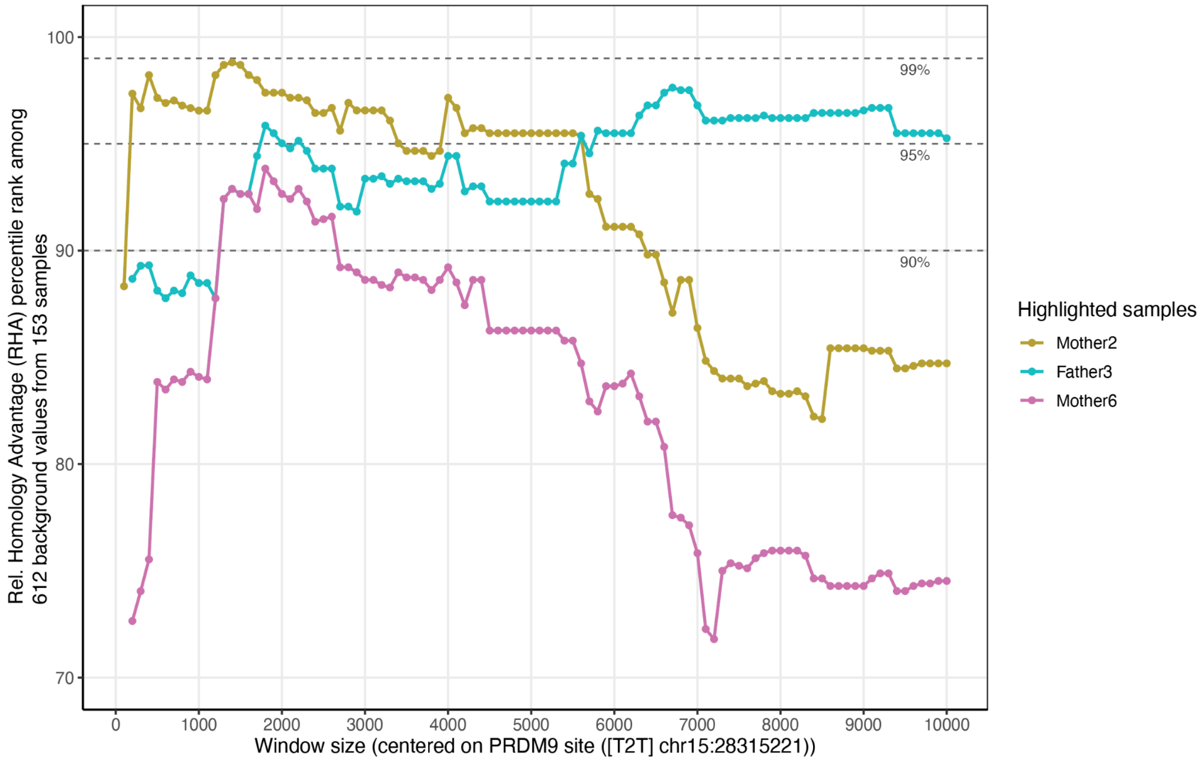


**Supplementary Figure 13. The rank Relative Homology Advantage (RHA) scores of the three parent samples among 612 control RHAs from 153 samples, as a function of window size around the PRDM9 motif.** RHA is not highest directly at the PRDM9 motif but peaks in a window of ~1100 - 2500 bp, indicating the RHA principle works on these size ranges. See Methods for RHA calculation and sample inclusion.

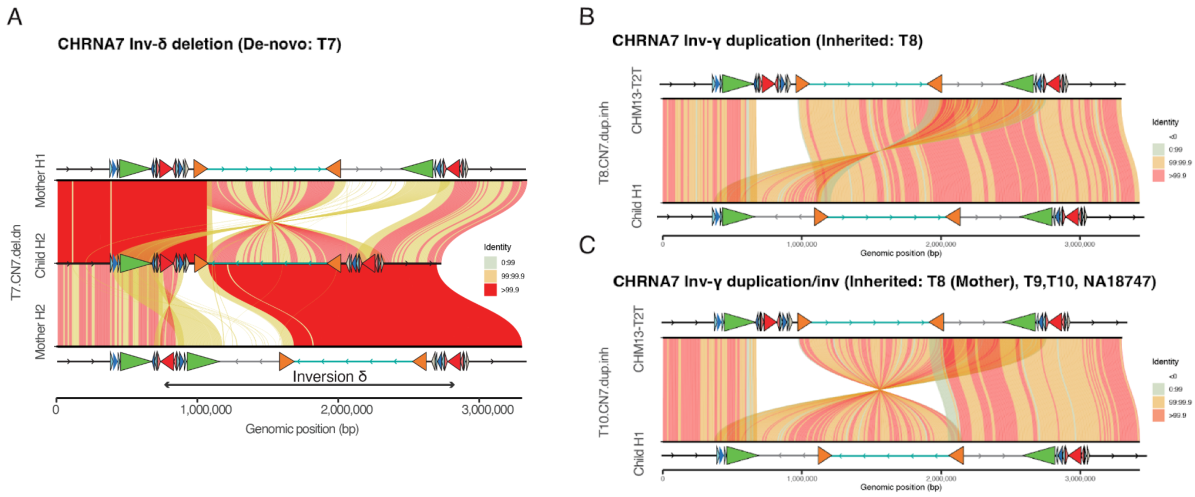


**Supplementary Figure 14. Assembly visualization of representations of all resolved CHRNA7 CNVs.** A) T7 carries a de novo CHRNA7 deletion, mediated by Inv-δ. HORs are overlaid. see Figure 3A. B) A CHRNA7 duplication found in the father and index of T8, aligned to T2T-CHM13. The allele carries an inverted duplication of the segment between CHRNA7-LCR′ (orange triangle) and SDα (green triangle), but also lacks the region between SDα and CHRNA7-LCR; including a copy of SDβ (red triangle). The CNV configuration is perfectly explained by interchromosomal NAHR between CHRNA7-LCR and CHRNA7-LCR′ situated on a direct and opposite haplotype, as illustrated in Figure 3C. C) A second CHRNA7 duplication allele is observed in 4 out of 5 samples (also counting NA18747), where the central segment between CHRNA7-LCR and CHRNA7-LCR′ is also inverted. Again, this configuration is consistent with interchromosomal NAHR (Figure 3C).

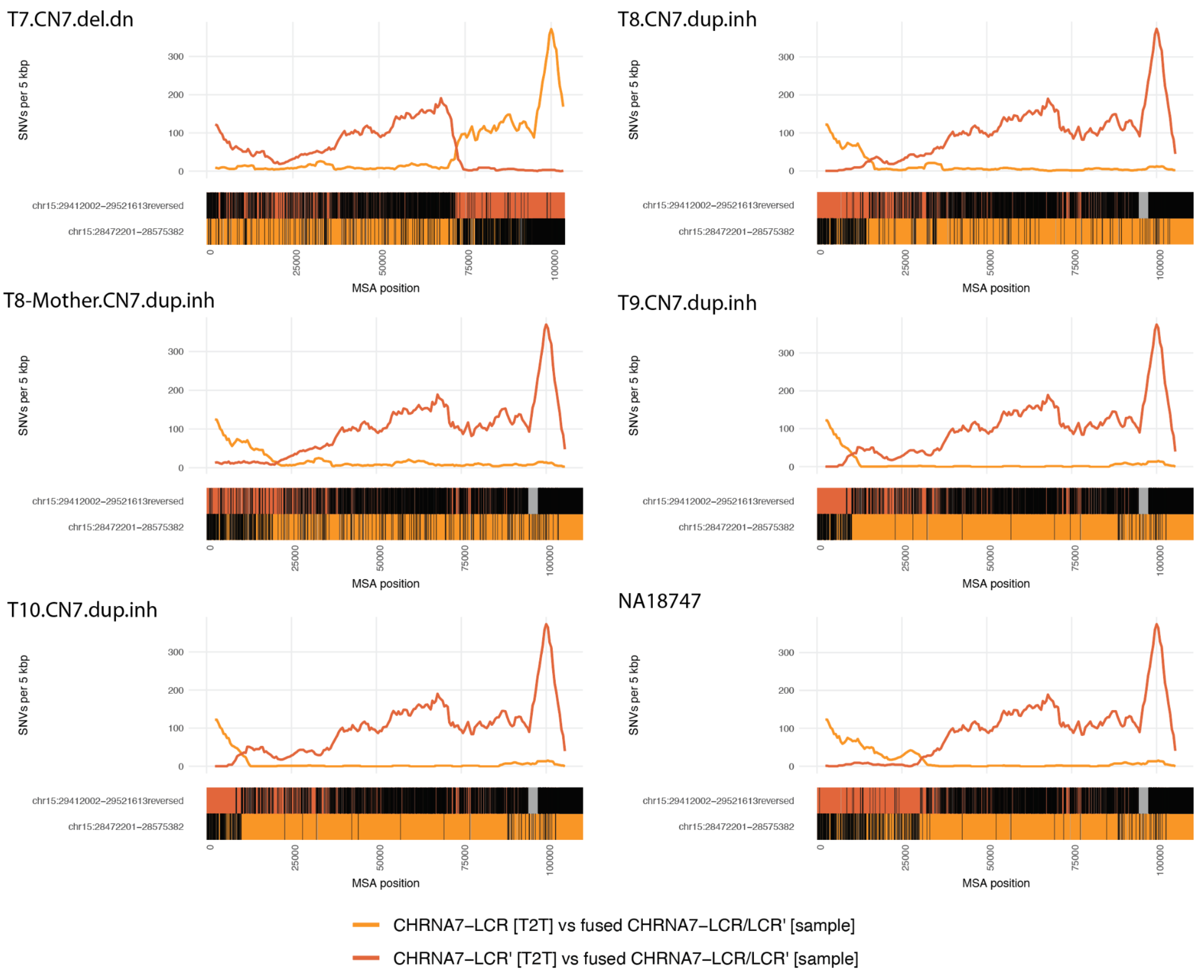


**Supplementary Figure 15. Sequence-based inference of ancestral CHRNA7 CNV breakpoints.** Three-way alignments between (presumed) fused CHRNA7-LCR/CHRNA7-LCR′ segments in CNV alleles, and [T2T-CHM13] CHRNA7-LCR (orange) and [T2T-CHM13] CHRNA7-LCR′ (red). Mismatches are indicated in black. The approximate ancestral breakpoint positions emerge as a ‘switch’ from preferred match between the reference alleles. All CHRNA7-CNVs show signs of ancestral recombination as predicted by our model. The CNVs in T9 and T10 likely originate from the same event, while T8 (child, father) and T8 (mother) have arisen as separate events.

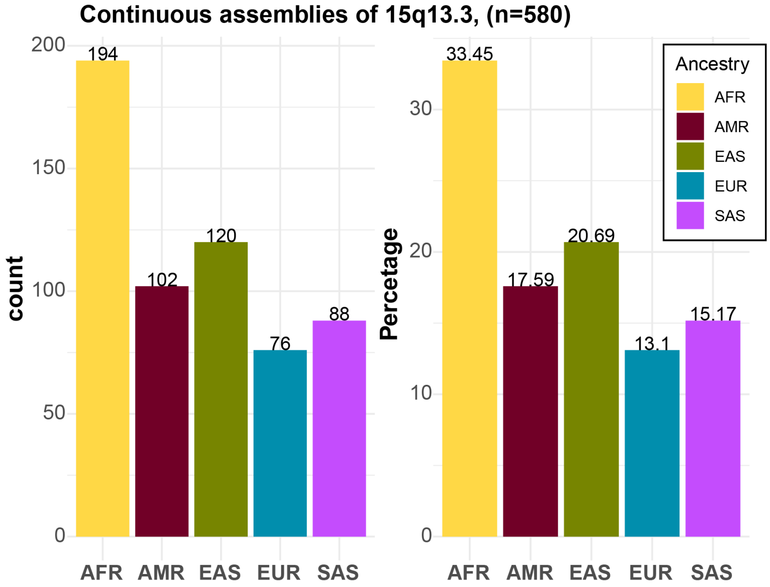


**Supplementary Figure 16. Ancestry summary of all continuous assemblies of 15q13.3** Ancestry representation of n=580 (excluding T2T-CHM13 reference) fully assembled haplotypes among HPRC and HGSVC phased genome assemblies. The left barplot shows total counts while right one percentages of total sample count. Each bar is colored by corresponding ancestry (AFR - African, AMR - American, EAS - East Asian, EUR - European, and SAS - South Asian).

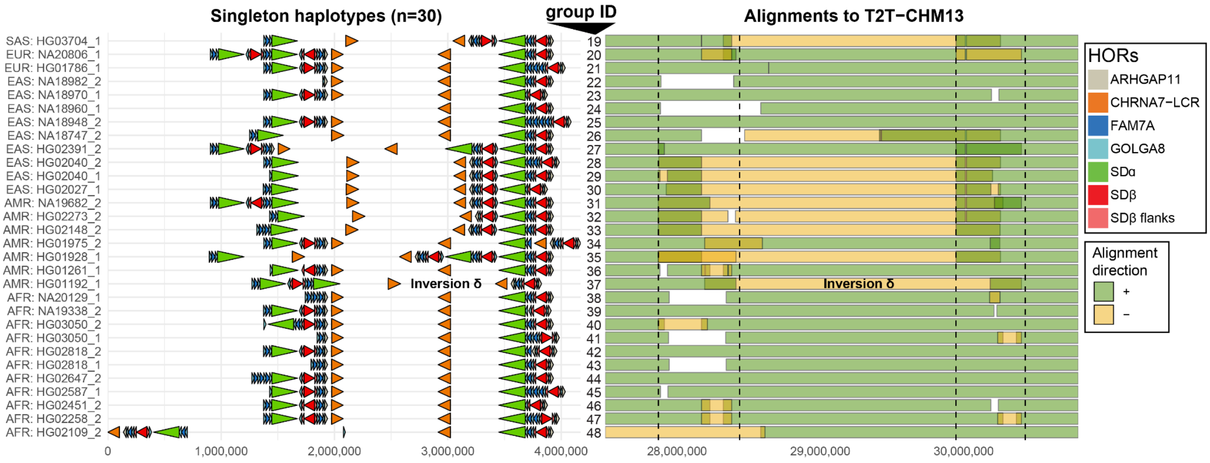


**Supplementary Figure 17. 15q13.3 HOR structures observed only in a single haplotype. A)** HOR annotation of the 30 haplotype structures (haplotype group 19-48) observed only in a single haplotype. Each HOR structure is defined by direction pointing arrowheads colored by the HOR unit (see legend). **B)** Alignment directionality of all singleton haplotypes with respect to T2T-CHM13. Forward oriented alignments are shown in green while oppositely oriented alignments in yellow (designating large-scale inversions and inverted duplications). Vertical dashed lines highlight positions of SD blocks where CNV and inversion breakpoints (BP4-left and BP5-right) reside.

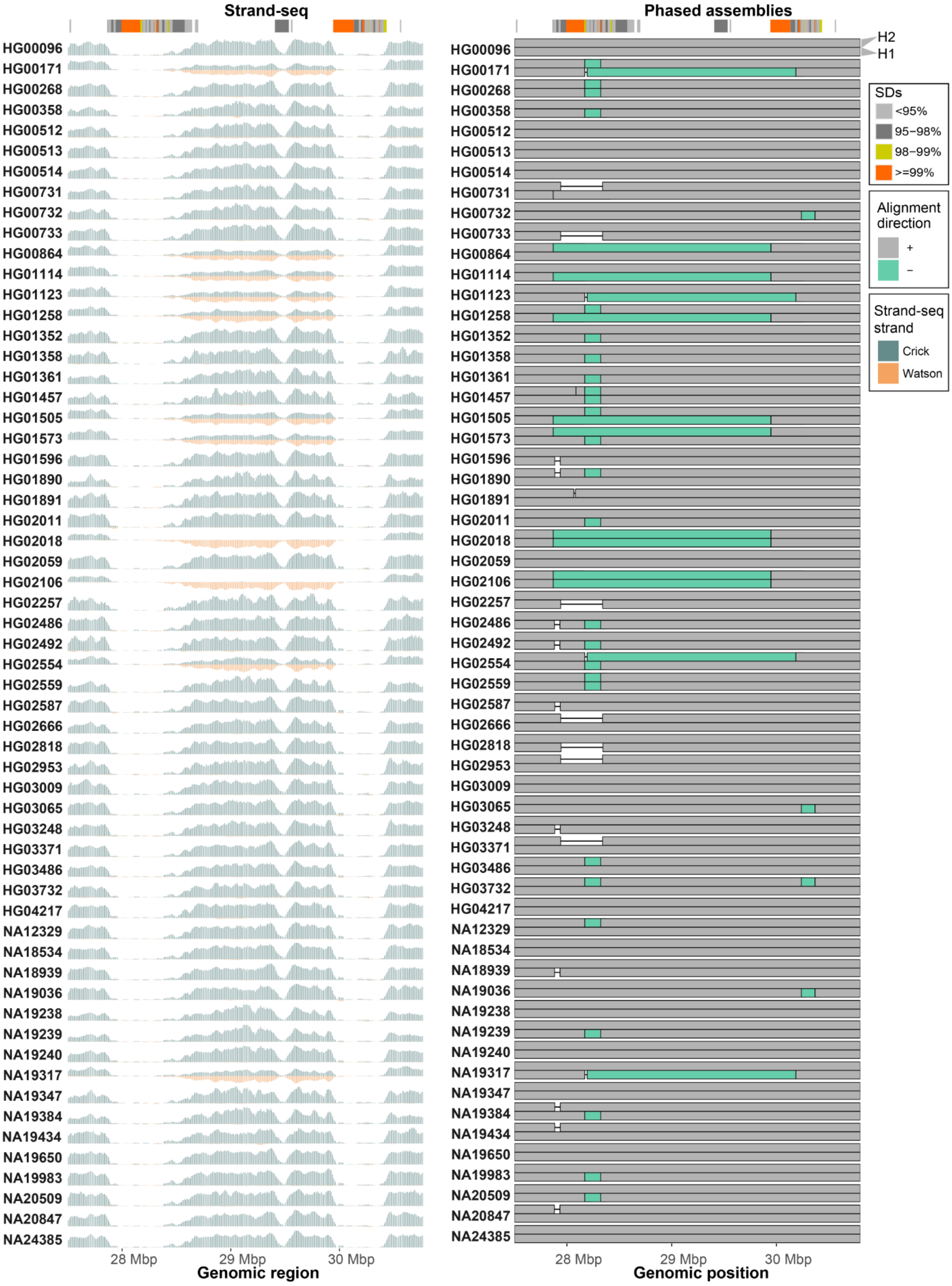


**Supplementary Figure 18. Assembly validation using Strand-seq.** Left: Read-coverage profiles of Strand-seq data over the 15q13.3 region summarized as binned (bin size: 50 kbp step size: 10 kbp) read counts represented as bars above (teal; Crick read counts) and below (orange; Watson read counts) the midline. The inversion polymorphism lies between the SD blocks (colored by sequence identity) marked at the top of the plot. Here, equal coverage of Watson and Crick counts represents a heterozygous inversion (e.g HG00171) as only one homologue is inverted with respect to the reference (T2T-CHM13). The homozygous inversion (e.g HG02018) is visible as complete switch from Crick to Watson reads over the inverted region. Right: Alignment directionality of fully assembled haplotypes (H1 - bottom and H2 top) for 59 samples (118 haplotypes) over the 15q13.3 region. Forward ('+') oriented alignments are shown in gray while reverse oriented alignments in light green ('-'). On top there is an SD annotation (SDs) colored by sequence identity.

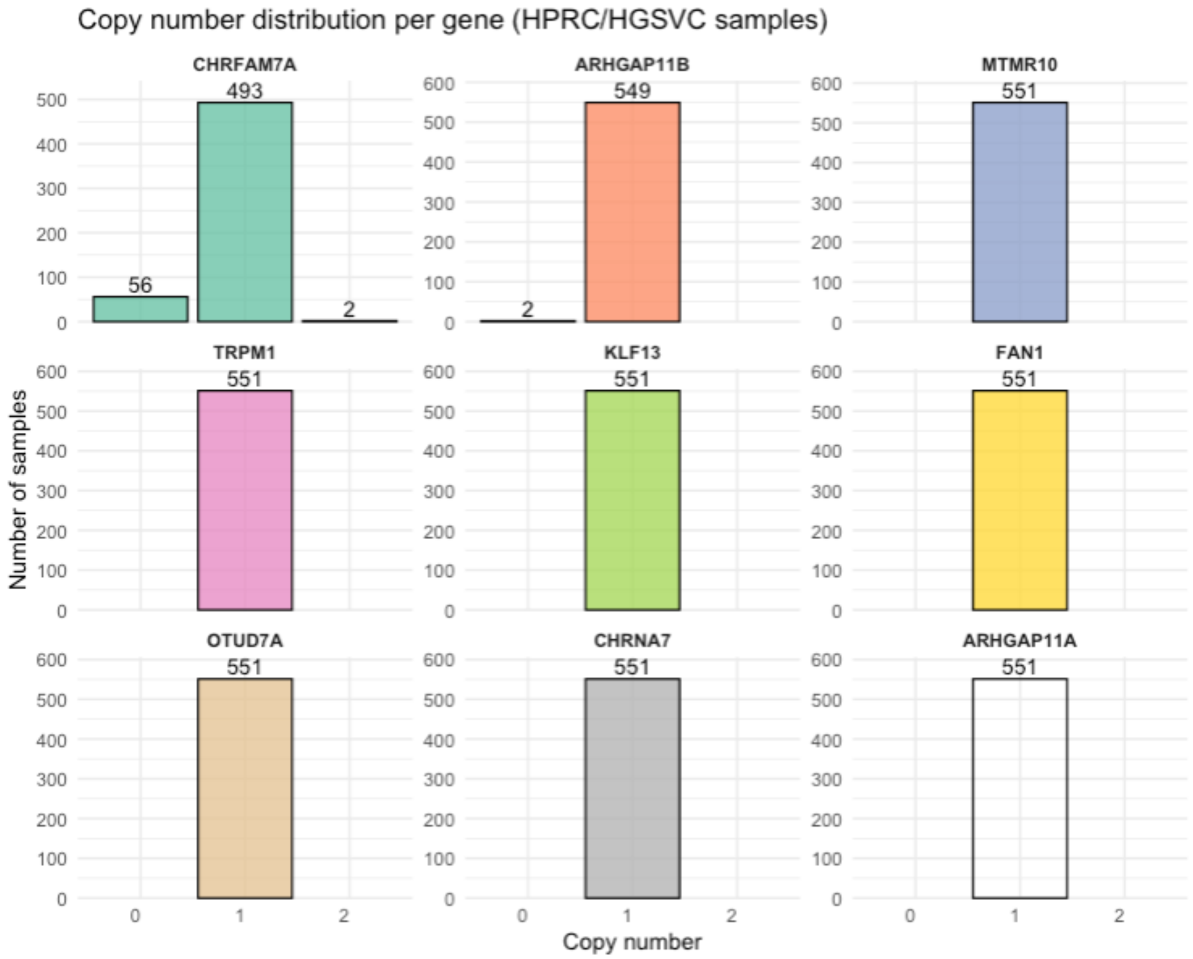


**Supplementary Figure 19. Copy numbers of major genes in the 15q13.3 region among 551 haplotypes representing the 18 major haplotype groups of healthy individuals from HPRC and HGSVC.** *CHRFAM7A* deletions are observed in ~10% of haplotypes, making those alleles potentially disease relevant as 'second-hits' modifying phenotypic severity of 15q13.3 CNVs.

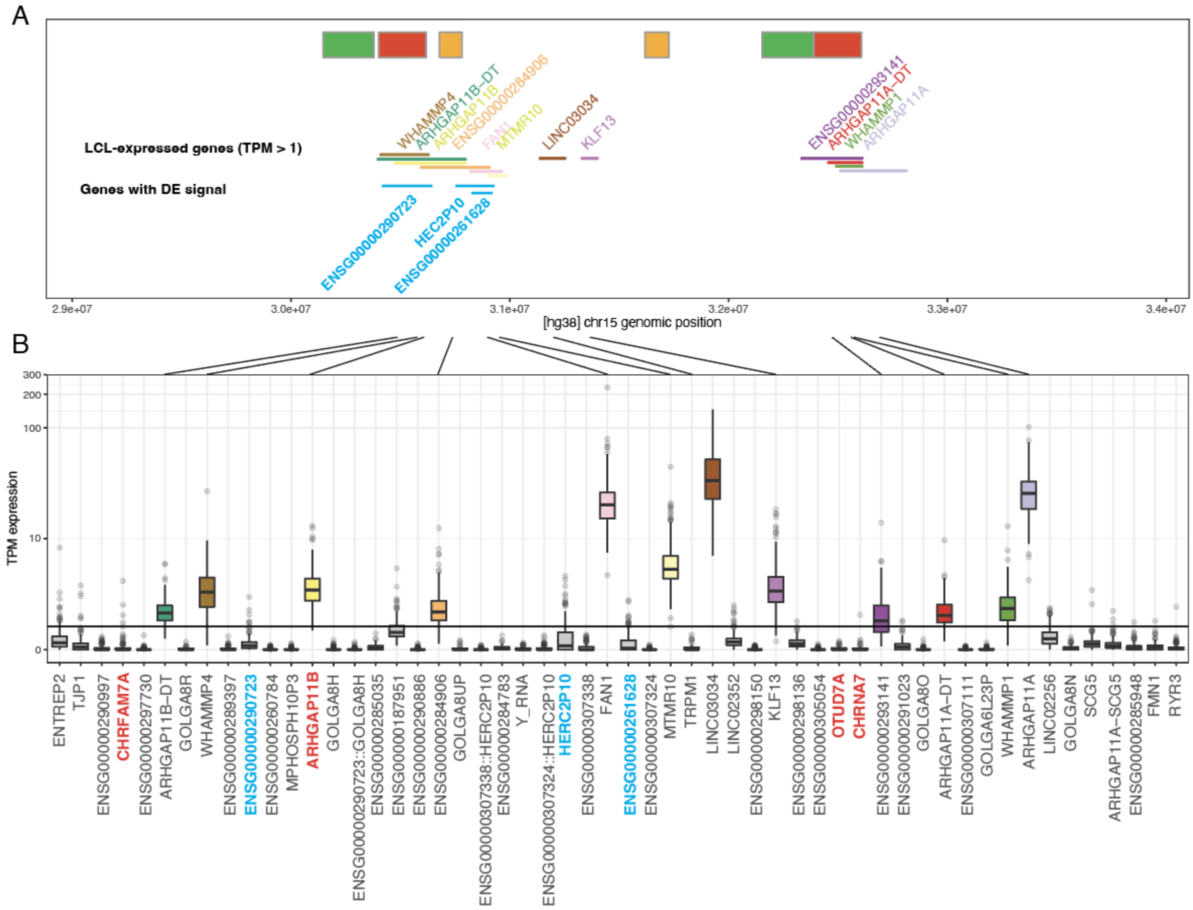

**Supplementary Figure 20. Location and expression of genes in the 15q13.3 region in lymphoblastoid cell lines (LCLs).** A) Visualization of genes with a median expression >1 TPM across n=206 long-read RNA-seq datasets. Major segmental duplication blocks are indicated above (green: SDα; red: SD-β; orange: CHRNA7-LCR). Genes whose expression is significantly correlated with at least one haplogroup are highlighted in blue. B) Boxplots showing TPM expression per sample and gene from the same n=206 samples. Among genes of interest for the incomplete penetrance of 15q13.3 copy number variants (*CHRFAM7A, ARHGAP11B, OTUD7A,* and *CHRNA7*, highlighted in red), only *ARHGAP11B* shows significant expression in LCLs. Genes with stable expression (median >1 TPM) are color-coded. Arrows link genes between panels A and B to indicate correspondence.

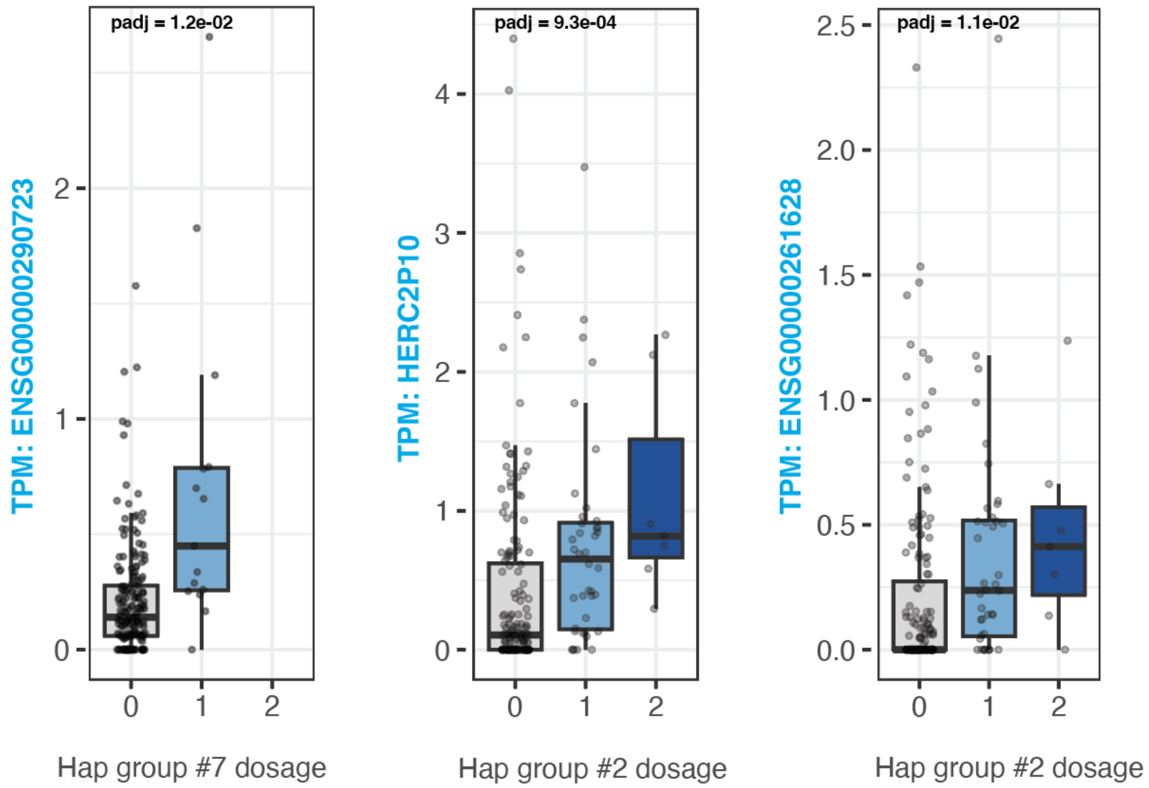


**Supplementary Figure 21. The three identified significant gene/haplogroup correlations.** Boxplots show expression levels for genes stratified by dosage of a particular haplogroup (7, 2 and 2 for ENSG00000290723, HERC2P10 and ENSG00000261628, respectively). Dosage values (0, 1, 2) indicate the number of copies of a given haplogroup present in each sample (0 = absent, 1 = heterozygous presence, 2 = homozygous presence). Data is based on n=206 samples (Methods).

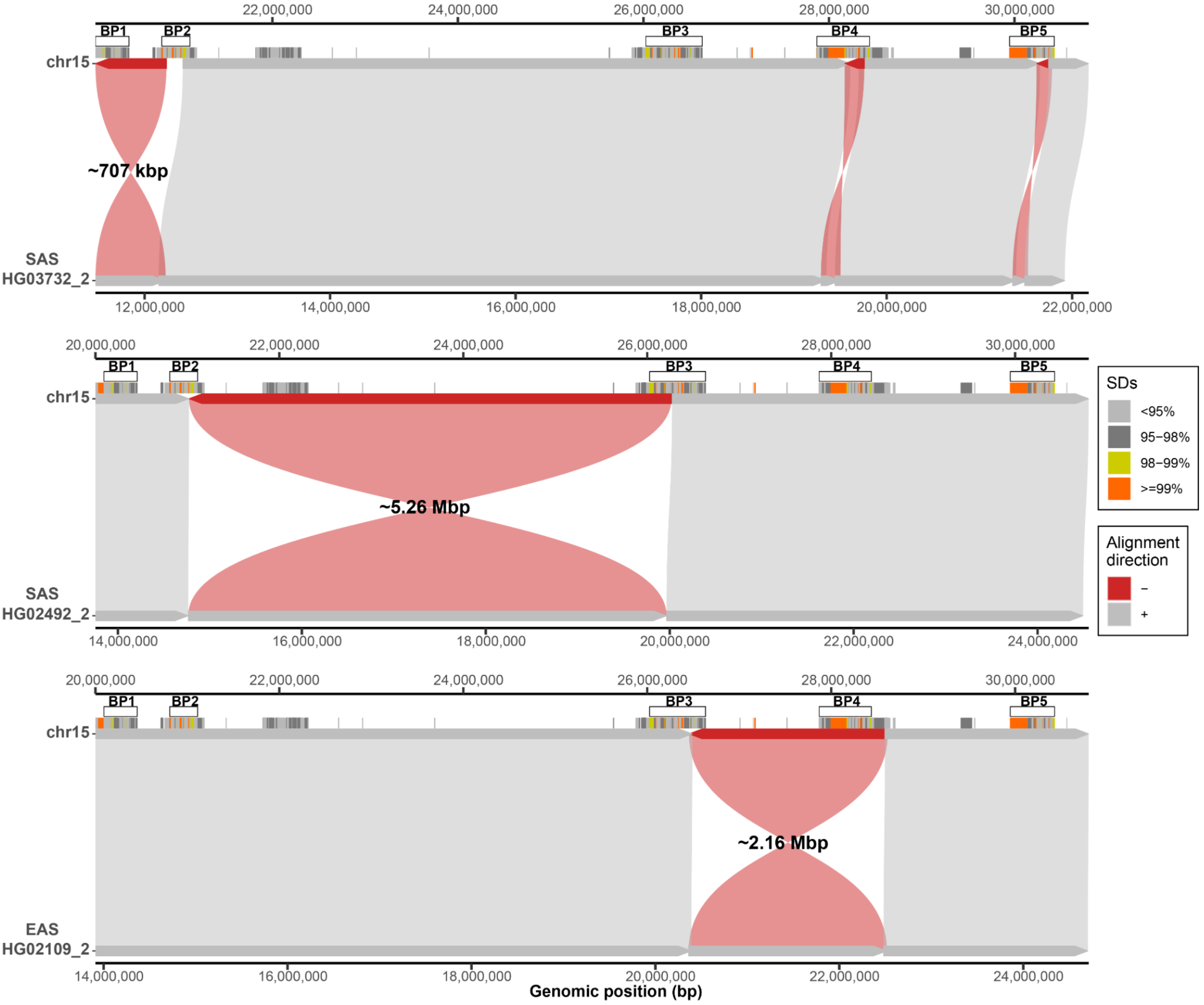


**Supplementary Figure 22. Synteny plots showing rare inversions between BP1-BP2, BP2-BP3, and BP3-BP4.** The direct (T2T-CHM13 reference) haplotype is shown on the top and inverted haplotype at the bottom. On top there is segmental duplication annotation colored by their sequence identity along with marked positions of BP1-BP5. Direct ('+') alignments are shown gray and inverted ('-') alignments in red.

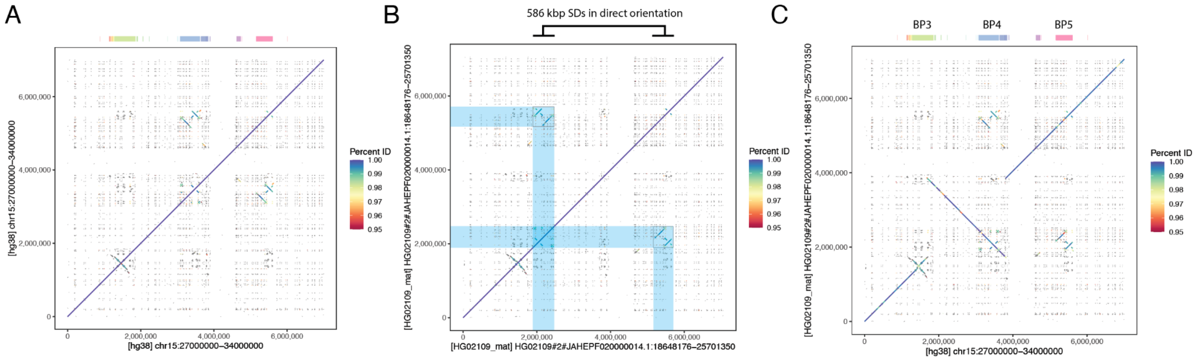


**Supplementary Figure 23. Dotplot view of the wider BP3-BP5 region in a HPRC sample carrying a BP3-BP4 inversion.** A) a dotplot of the reference sequence [HG38:27,000,000-34,000,000] plotted against itself visualizes the repeat structure (off-diagonal alignments) of BP3, 4 and 5. B) a dotplot of the same region in the inversion-carrying sequence of HG02109 (maternal haplotype) plotted against itself. 586 kbp of directly oriented SDs (highlighted in blue) likely make the structure highly susceptible to NAHR. C) A dotplot of the reference sequence (Panel A) versus the variant sequence (Panel B) visualizes the large inversion between BP3 and BP4.

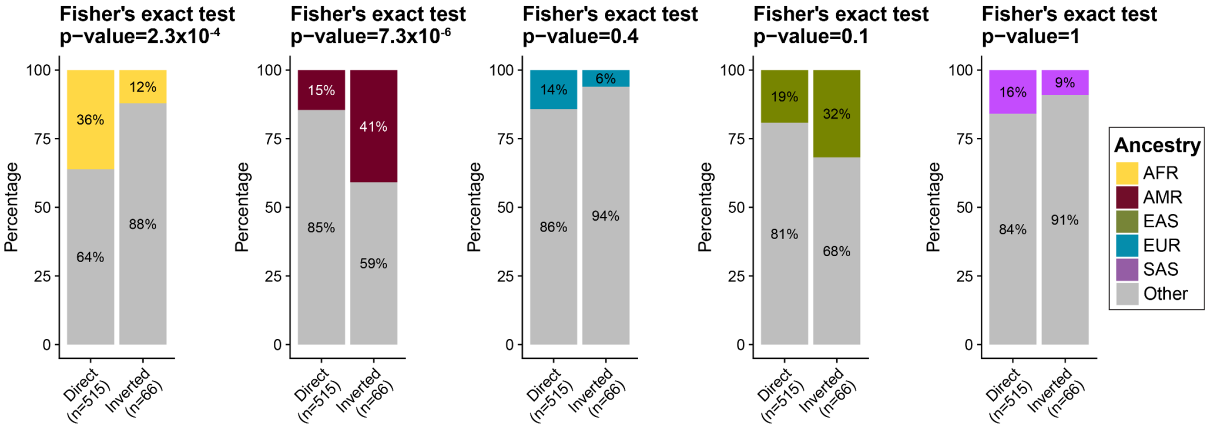


**Supplementary Figure 24. Enrichment of BP4-5 inversion among different ancestries.** Evaluation of the proportions of each ancestry (AFR - African, AMR - American, EUR - European, EAS - East Asian, and SAS - South Asian) among haplotypes defined as either direct (no inversion) and inverted between BP4 and BP5 with respect to the rest of the ancestries ('Other' - gray). We used Fisher's exact test (two-sided) to test the significance of the difference between these proportions. Above each barplot is a reported p-value after Bonferroni correction. Sample numbers are indicated below each plot.

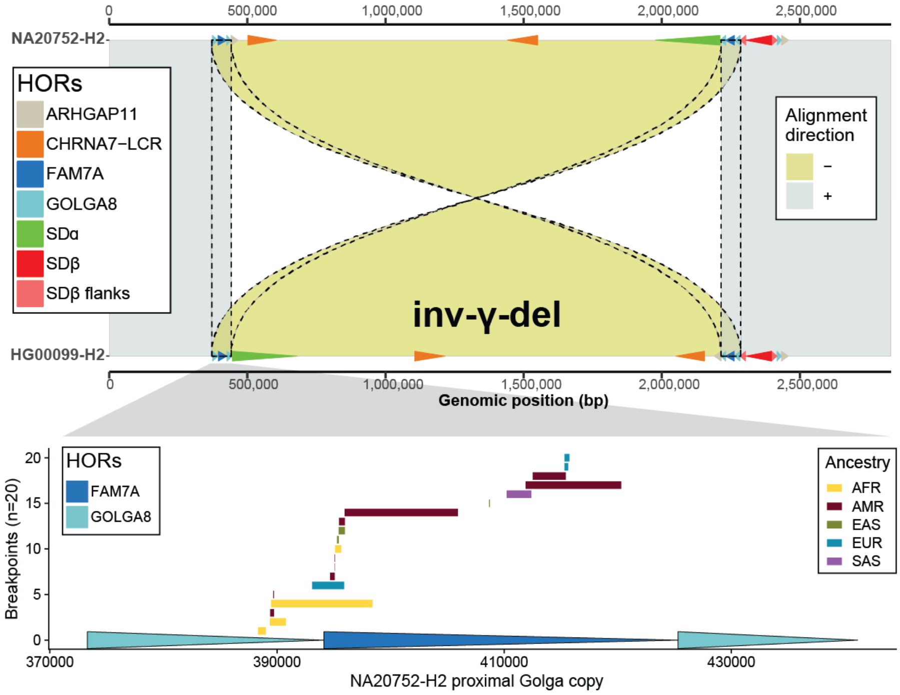


**Supplementary Figure 25. Inversion breakpoint mapping for Inv-γ-del.** A synteny plot showing an example Inv-γ-del. The direct (reference) haplotype is shown on the top and inverted haplotype at the bottom. Each haplotype has HOR annotation shown on top. Direct ('+') alignments are shown gray and inverted ('-') alignments in yellow. Redundant alignments at the inverted repeats implicated in inversion formation are highlighted by dashed outline. Below there are mapped inversion breakpoints for Inv-γ-del. Further below there is annotation of inverted HORs implicated in inversion formation. Inversion breakpoints are shown with respect to a proximal repeat copy. Breakpoints are reported with respect to single reference haplotype NA20752-H2.

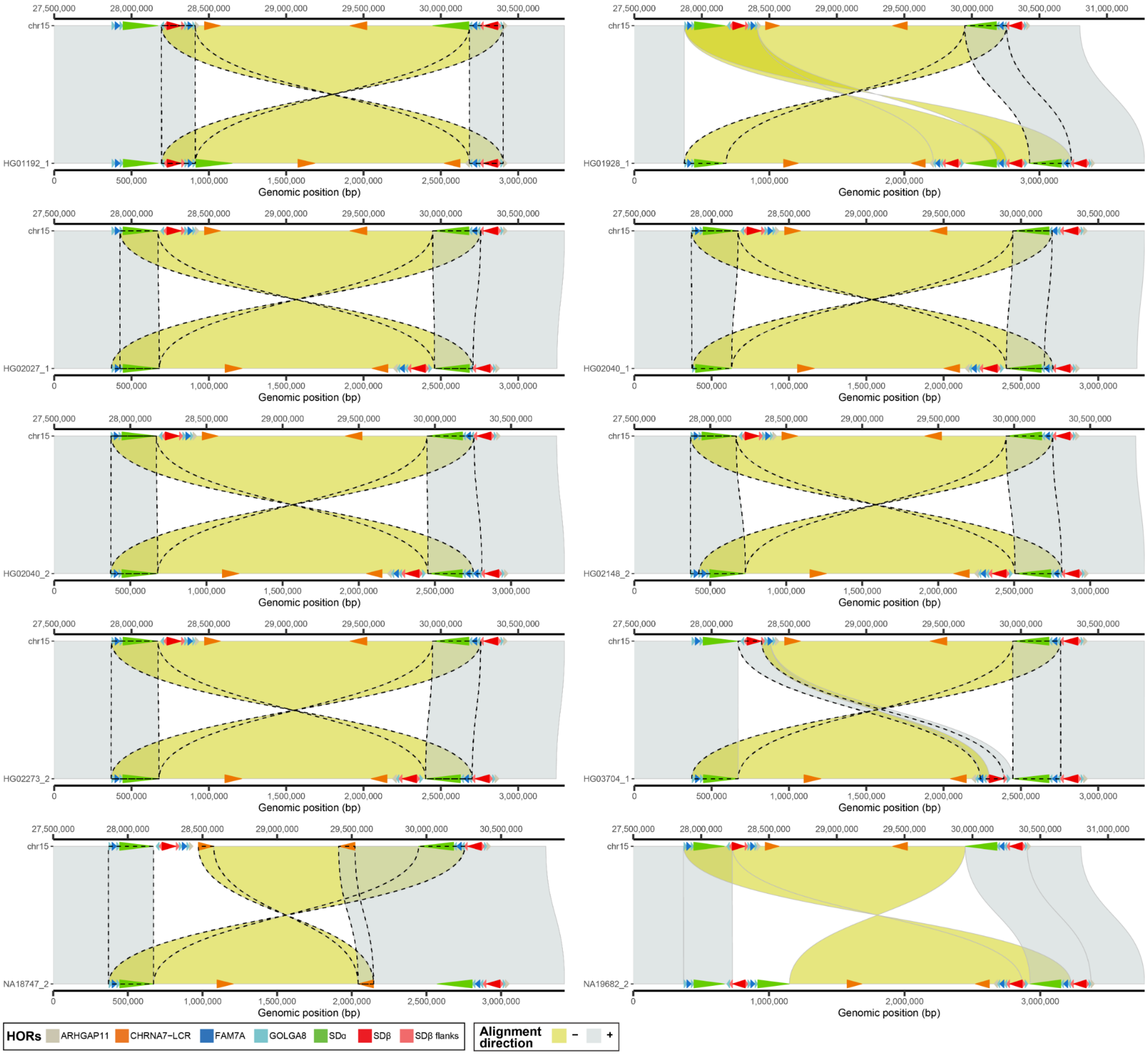


**Supplementary Figure 26. Synteny plots for 10 singleton inversions.** Synteny plots showing additional rare haplotypes with inversion between BP4 and BP5. The direct (T2T-CHM13 reference) haplotype is shown on the top and inverted haplotype at the bottom. Each haplotype has HOR annotation shown on top. Direct ('+') alignments are shown in gray and inverted ('-') alignments in yellow. Redundant alignments at the inverted repeats implicated in inversion formation are highlighted by dashed outline.

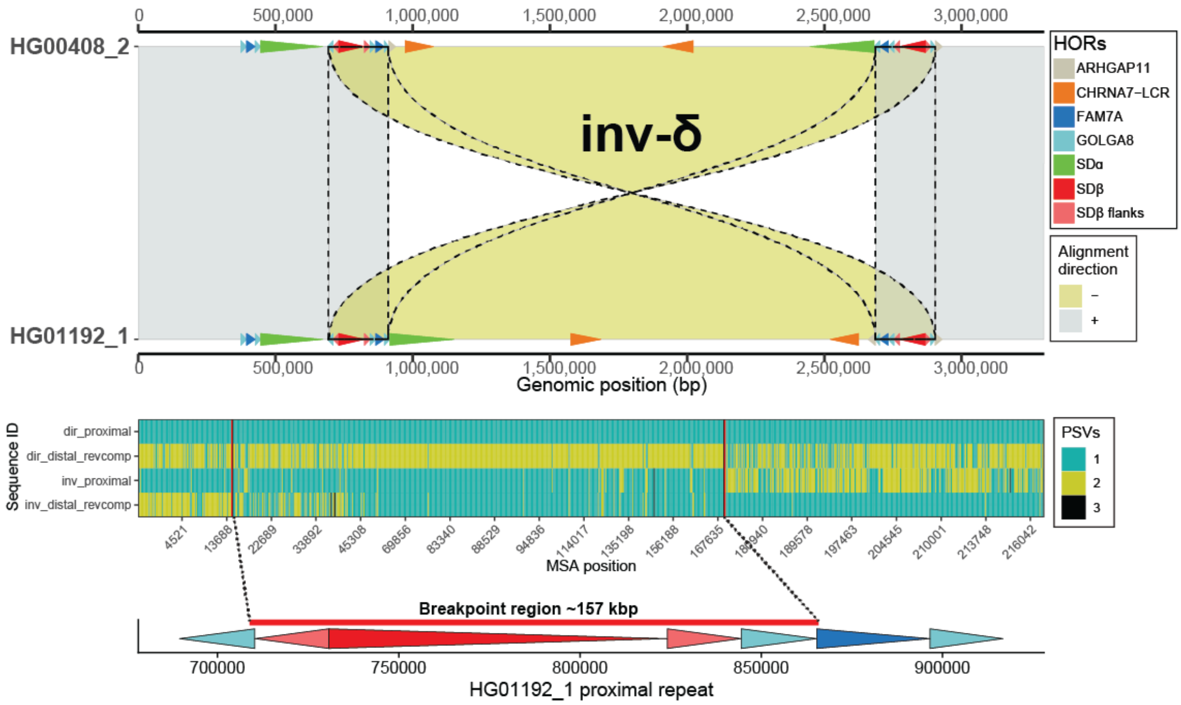


**Supplementary Figure 27. Inversion breakpoint mapping for Inv-δ.** A synteny plot showing an example Inv-δ. The direct (reference) haplotype is shown on the top and inverted haplotype at the bottom. Each haplotype has HOR annotation shown on top. Direct ('+') alignments are shown gray and inverted ('-') alignments in yellow. Redundant alignments at the inverted repeats implicated in inversion formation are highlighted by dashed outline. Below there is a visualization of the multiple sequence alignment showing the transition between paralog specific alleles observed in inverted haplotype. Last we show the mapped inversion breakpoint with respect to HOR structure of the proximal repeat.

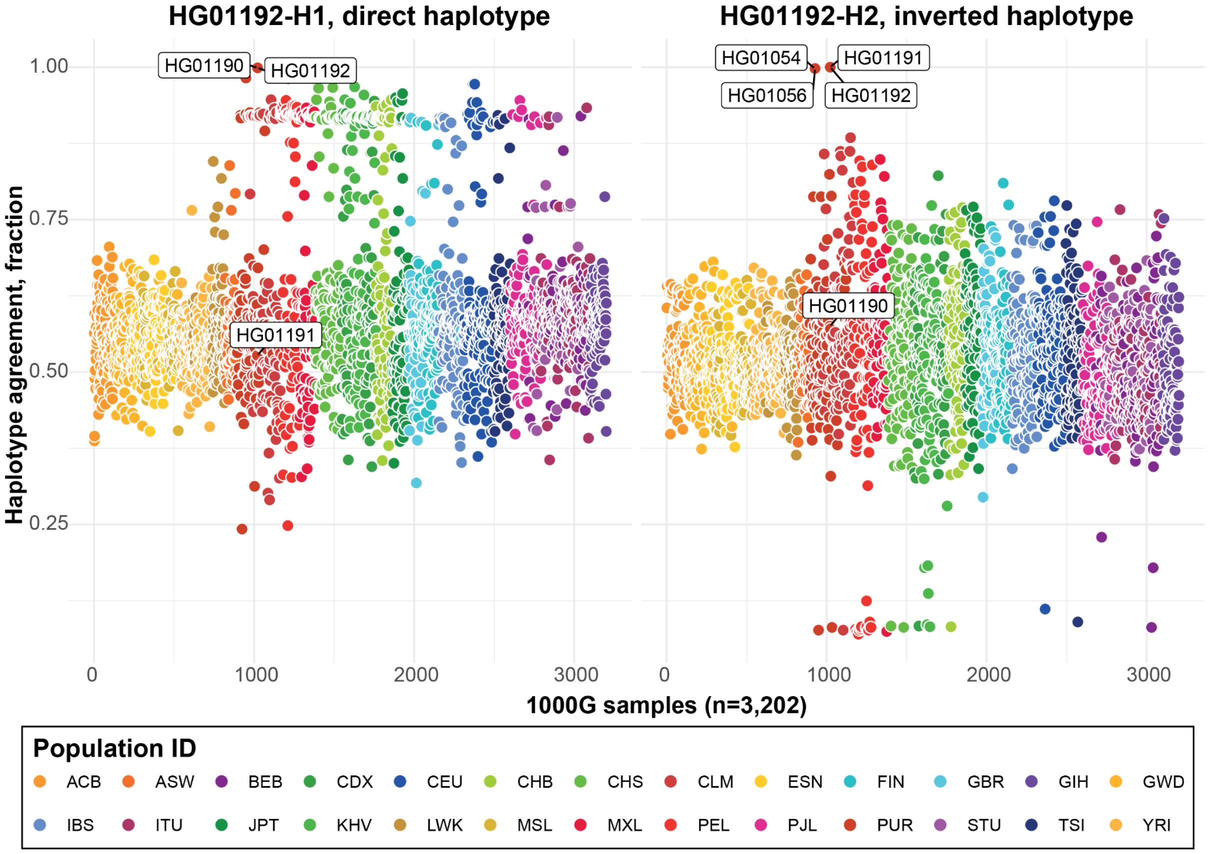


**Supplementary Figure 28. Genotyping of Inv-δ in the 1KG panel.** Haplotype agreement (y-axis, fraction of matching alleles) between inverted haplotype with respect to 1KG sample panel (n=3,202, T2T-CHM13 coordinates) [3]. Each dot represents a single sample colored by a population ID of five major ancestries (African - shades of orange, American - shades of red, East Asian - shades of green, European - shades of blue, and South Asian - shades of purple). Given the observed haplotype agreement between inverted haplotypes and 1KG samples, we report that inverted haplotype (H2) in HG01192 was inherited from the mother (HG01191). We also observe an additional family duo that carries the given inversion haplotype (HG01056 - child and HG01054 - father).

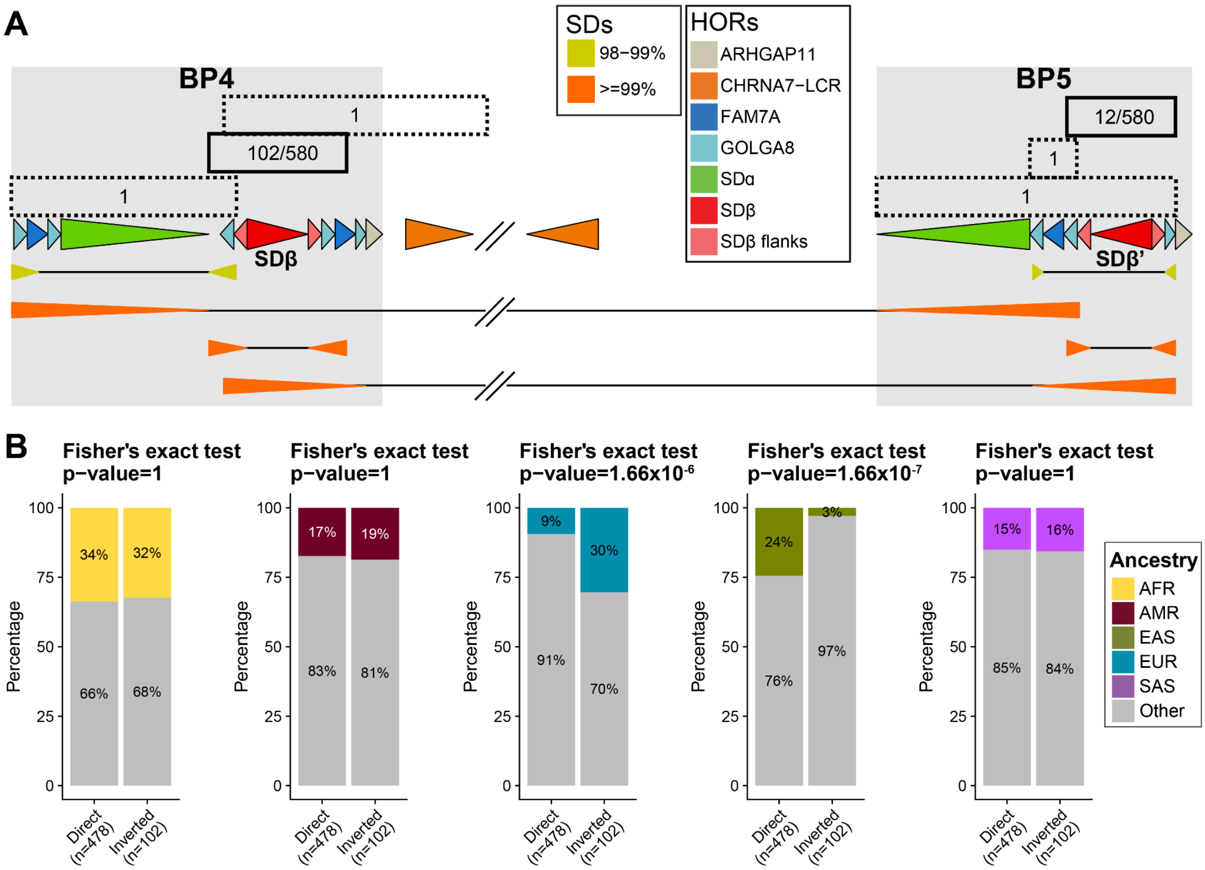


**Supplementary Figure 29. Inverted duplications and enrichment of SDβ inversion among different ancestries.** A) Visualization of the most frequent (solid lines) and singleton inverted duplications (dashed lines) within proximal (BP4) and distal (BP5) breakpoints in T2T-CHM13 coordinates. Below there is an annotation of HOR units and SD annotation showing the most identical inverted repeats in this region. B) Evaluation of the proportions of each ancestry (AFR - African, AMR - American, EUR - European, EAS - East Asian, and SAS - South Asian) of SDβ (102 out of 580 haplotypes) among haplotypes defined as either direct (no inversion) and inverted between BP4 and BP5 with respect to the rest of the ancestries (‘Other’ - gray). We used Fisher’s exact test (two-sided) to test the significance of the difference between these proportions. Above each barplot is a reported p-value after Bonferroni correction.

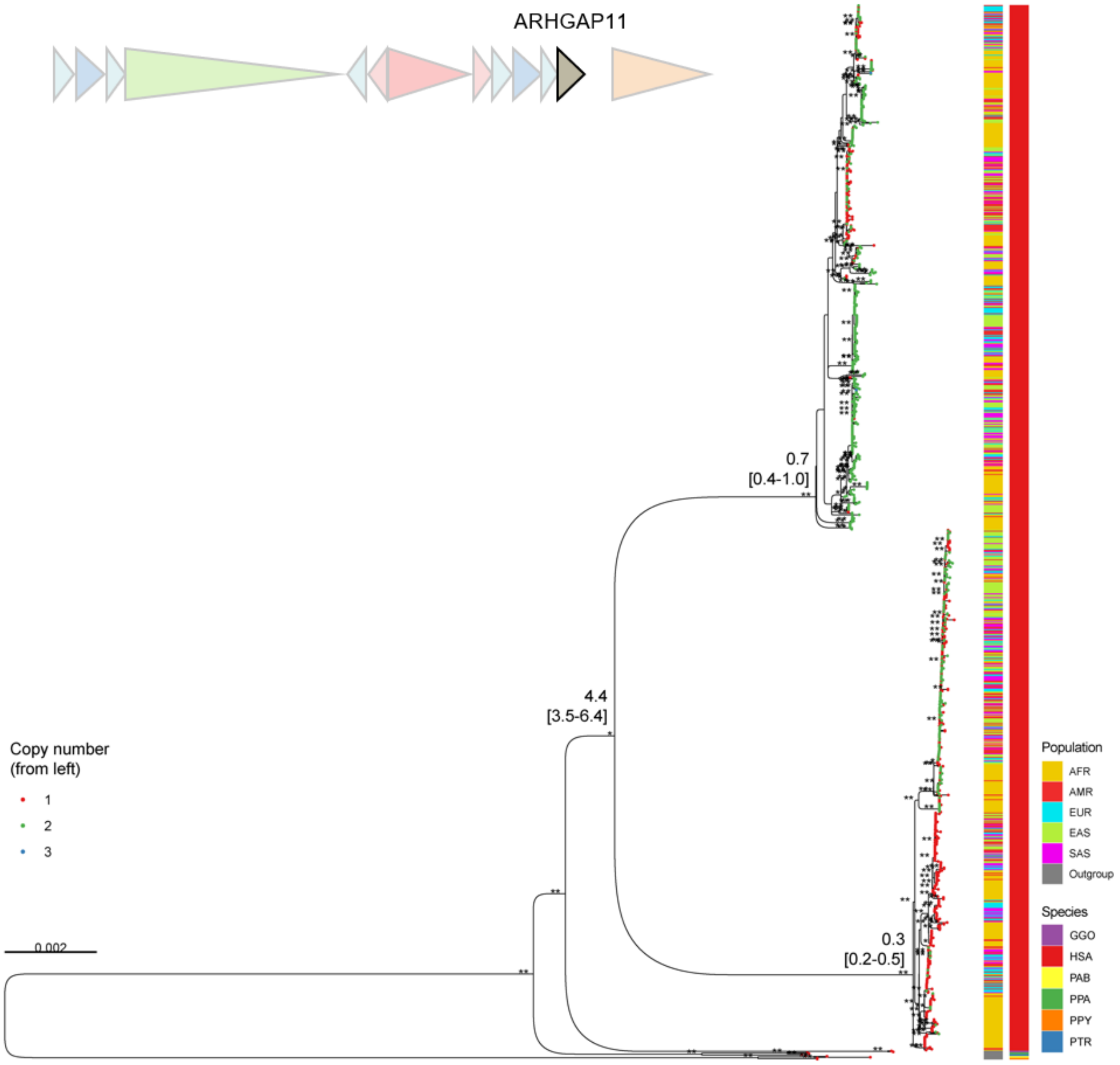


**Supplementary Figure 30. Maximum likelihood phylogenetic tree of the *ARHGAP11* duplicon.** The tree includes 1174 + 1 macaque outgroup sequences with conserved alignment length of 25 kbp. The tree was rooted with the ortholog copy from a macaque genome (not displayed due to long branch length). In addition, the tree also includes ape orthologs from diploid assemblies of chimpanzee (PTR), bonobo (PPA), gorilla (GGO), Bornean orangutan (PPY), and Sumatran orangutan (PAB). Bootstrap support was indicated by ** (>95) and * (>90). The average divergence times are indicated along with the confidence interval from 100 replicates below. From the 5′ the copy number was counted and indicated by different color of tip nodes. Ancestries of human haplotypes are indicated on the right as well as the species codes for nonhuman genomes.

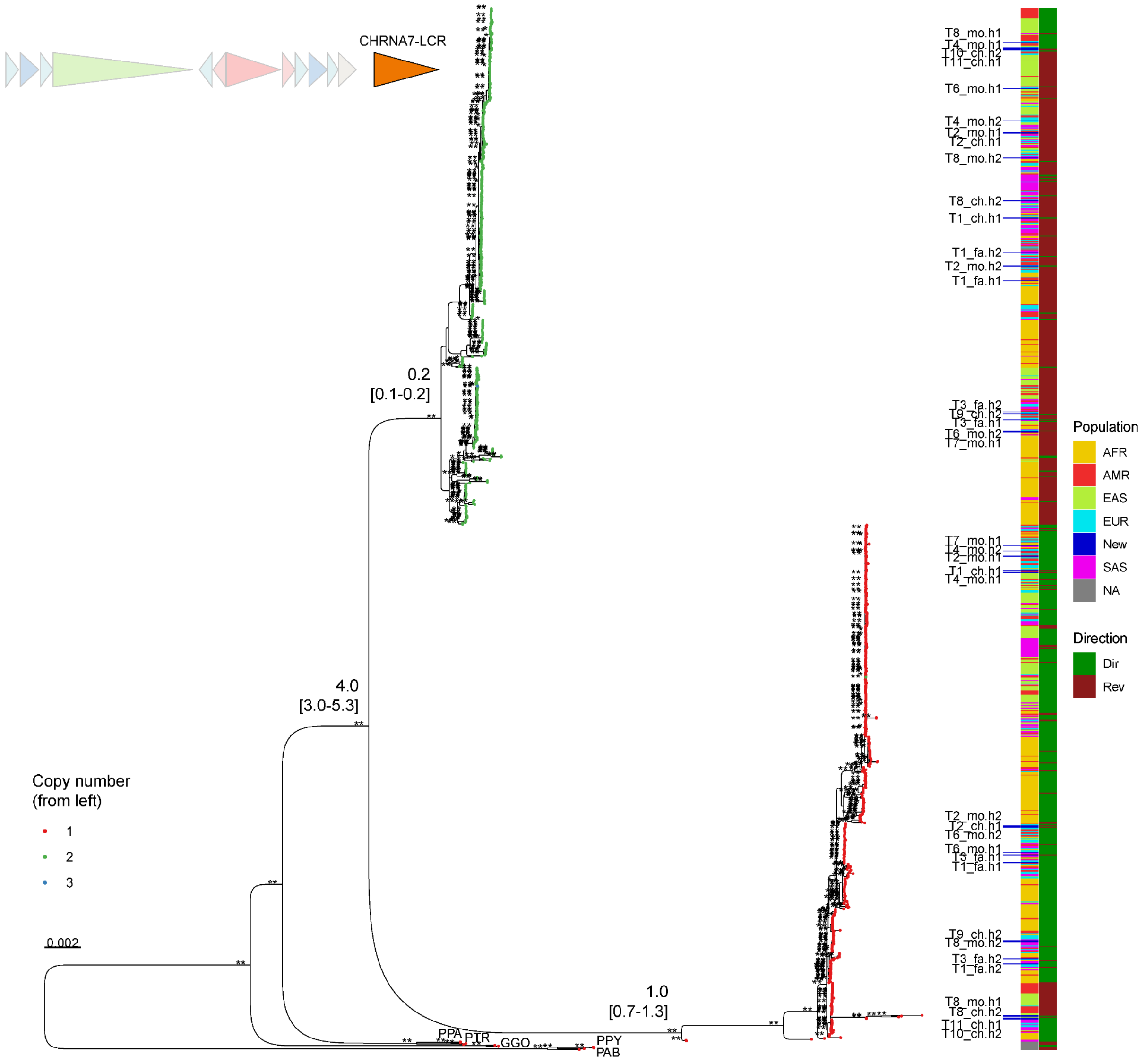


**Supplementary Figure 31. Maximum likelihood phylogenetic tree of the CHRNA7-LCR duplicon.** The tree includes 1214 (including trios sequenced in the current study) + 1 macaque outgroup sequences with conserved alignment length of 104 kbp. The tree was rooted with the ortholog copy from a macaque genome (not displayed due to long branch length). In addition, the tree also includes ape orthologs from diploid assemblies of chimpanzee (PTR), bonobo (PPA), gorilla (GGO), Bornean orangutan (PPY), and Sumatran orangutan (PAB). Bootstrap support was indicated by ** (>95) and * (>90). The average divergence times are indicated along with the confidence interval from 100 replicates below. From the 5′ the copy number was counted and indicated by different color of tip nodes. Ancestries of human haplotypes are indicated on the right with trio samples represented by blue. The direction of the duplicon sequence is indicated by green (direct) or red (opposite).

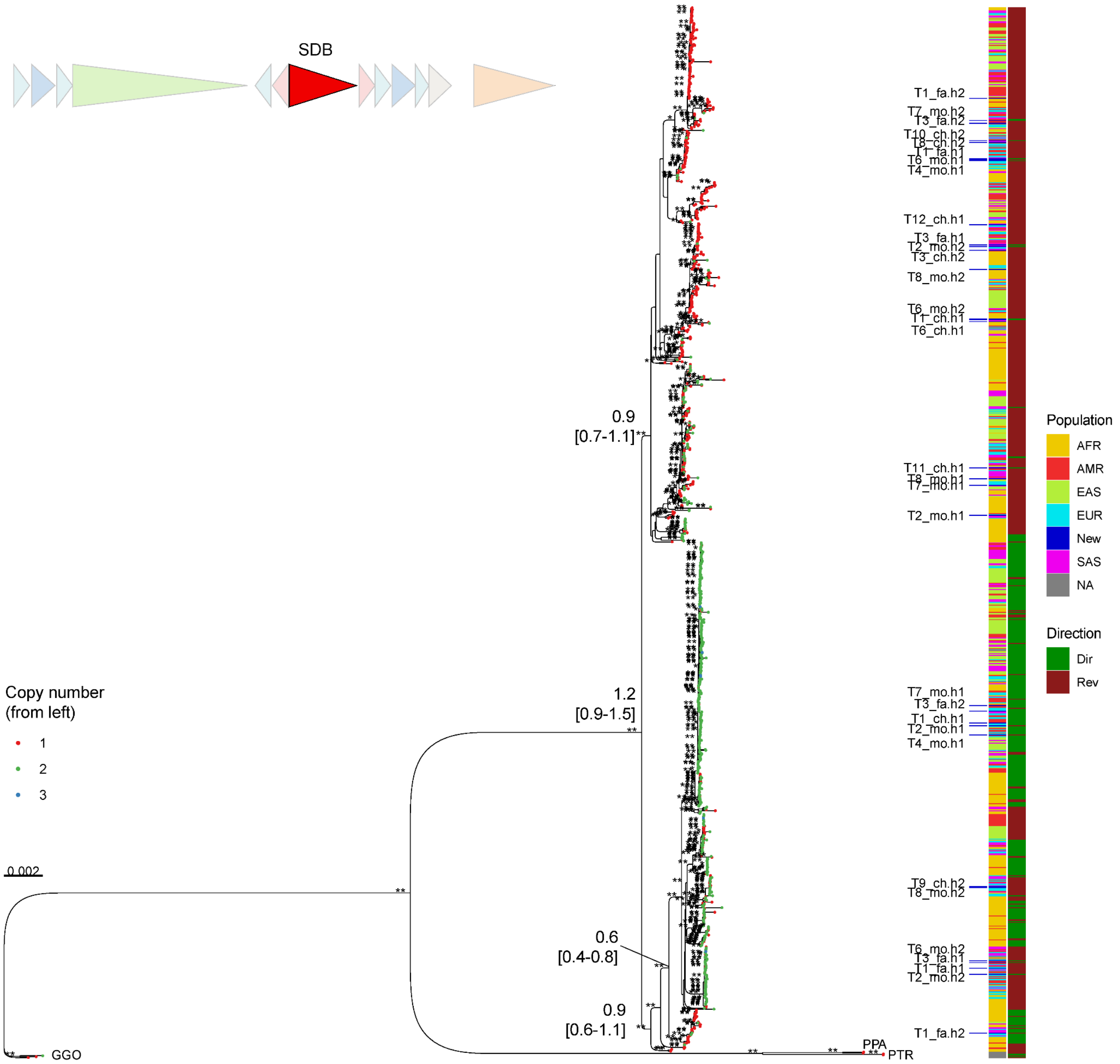


**Supplementary Figure 32. Maximum likelihood phylogenetic tree of the SDβ duplicon.** The tree includes 1149 (including trios sequenced in the current study) sequences with conserved alignment length of 93 kbp. The tree was rooted with the ortholog copy from gorilla (GGO) genome. In addition, the tree also includes ape orthologs from diploid assemblies of chimpanzee (PTR) and bonobo (PPA). Bootstrap support was indicated by ** (>95) and * (>90). The average divergence times are indicated along with the confidence interval from 100 replicates below. From the 5′ the copy number was counted and indicated by different color of tip nodes. Ancestries of human haplotypes are indicated on the right with trio samples represented by blue. The direction of the duplicon sequence is indicated by green (direct) or red (opposite).

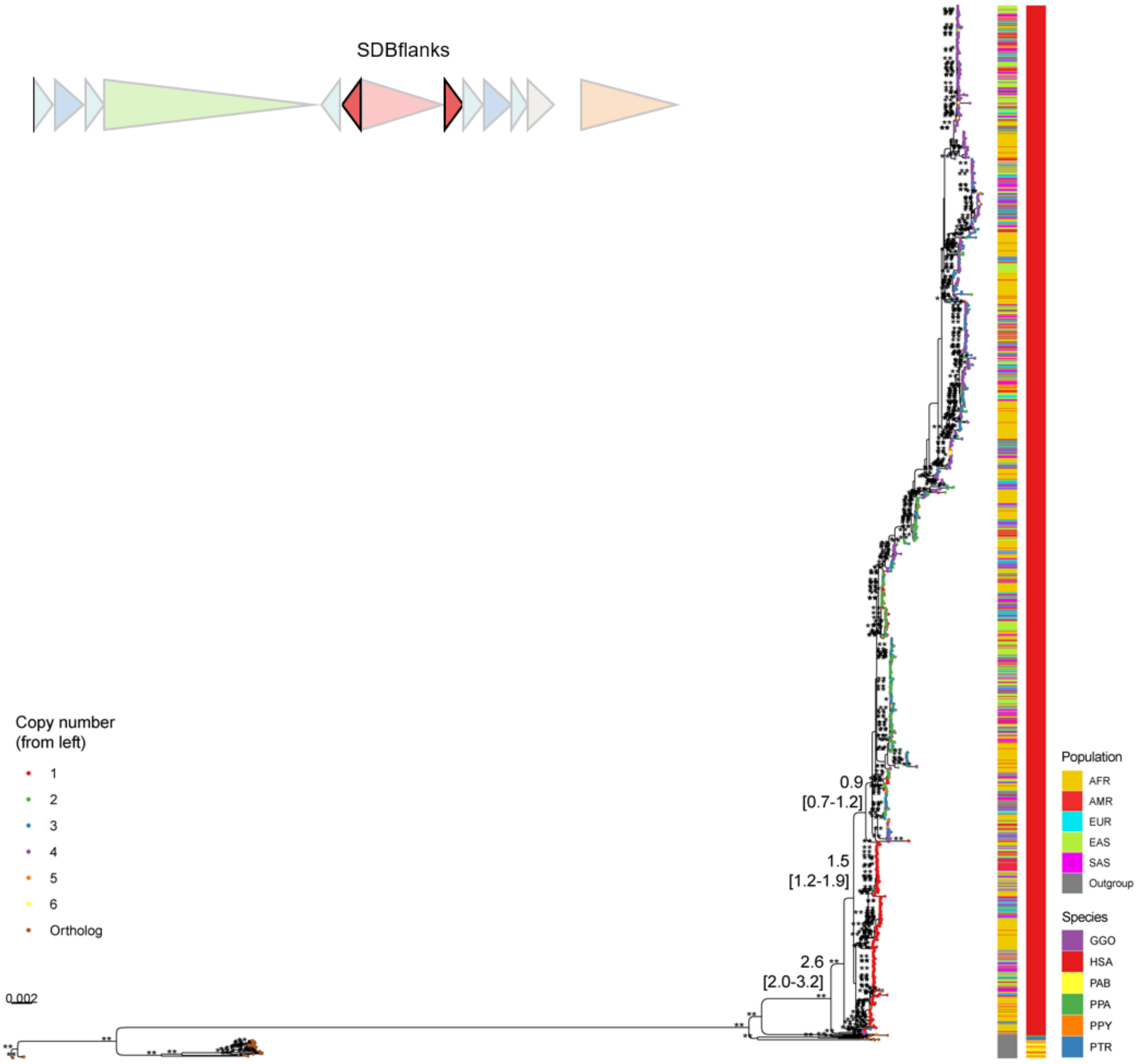


**Supplementary Figure 33. Maximum likelihood phylogenetic tree of the SDβ flanks duplicon.** The tree includes 2272 sequences with conserved alignment length of 20 kbp. The tree was rooted with the ortholog copies from the Sumatran orangutan (PAB) genome. In addition, the tree also includes ape orthologs from diploid assemblies of chimpanzee (PTR), bonobo (PPA), gorilla (GGO) and Bornean orangutan (PPY). Bootstrap support was indicated by ** (>95) and * (>90). The average divergence times are indicated along with the confidence interval from 100 replicates below. From the 5′ the copy number was counted and indicated by different color of tip nodes. Ancestries of human haplotypes are indicated on the right as well as the species codes for nonhuman genomes.

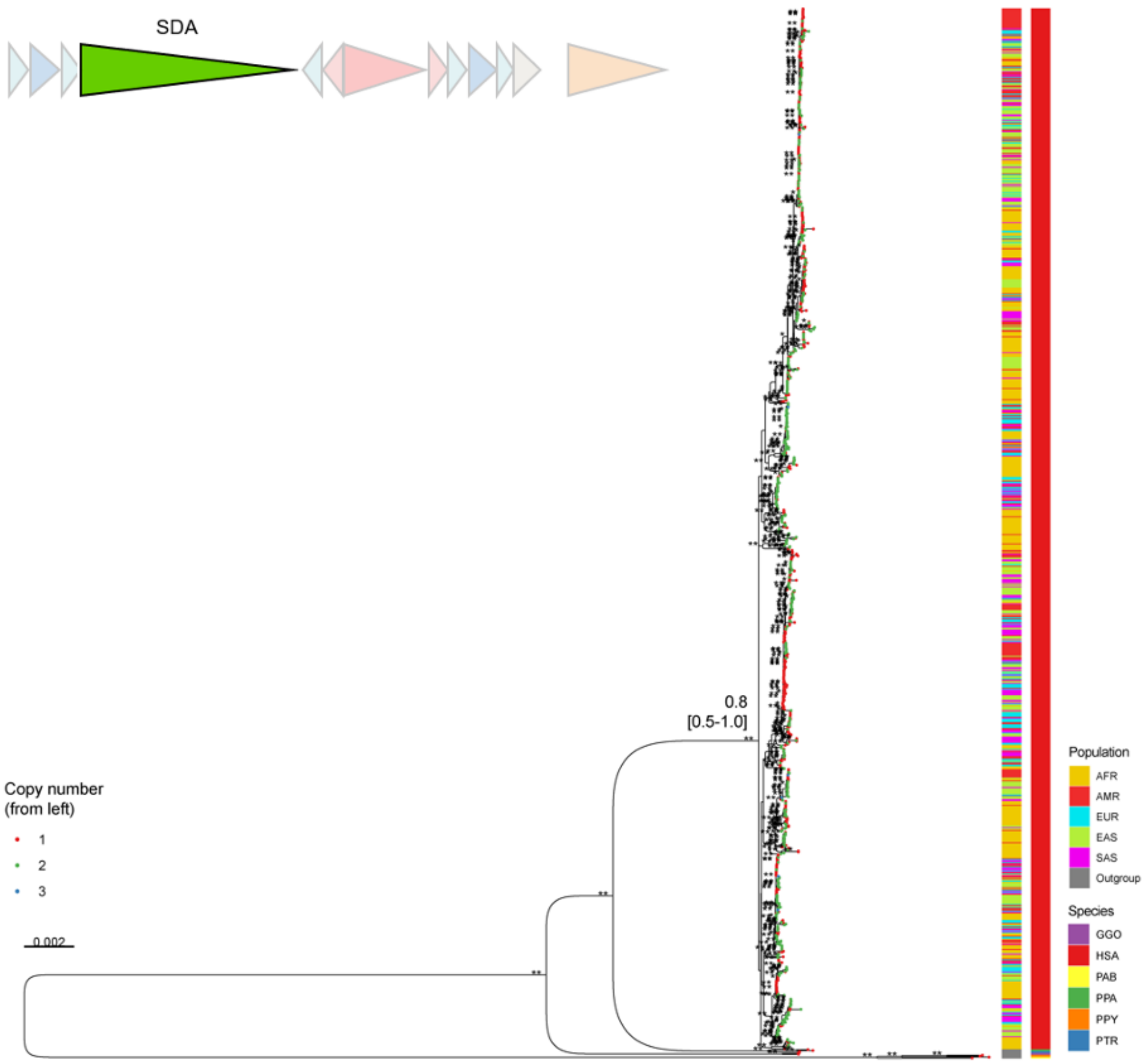


**Supplementary Figure 34. Maximum likelihood phylogenetic tree of the SDα duplicon.** The tree includes random 196 subset sequences including macaque outgroup with conserved alignment length of 49 kbp. The tree was rooted with the ortholog from a macaque genome (not displayed due to long branch length). In addition, the tree also includes ape orthologs from diploid assemblies of chimpanzee (PTR), bonobo (PPA), gorilla (GGO), Bornean (PPY), and Sumatran orangutans (PAB). Bootstrap support was indicated by ** (>95) and * (>90). The average divergence times are indicated along with the confidence interval from 100 replicates below. From the 5′ the copy number was counted and indicated by different color of tip nodes. Ancestries of human haplotypes are indicated on the right as well as the species codes for nonhuman genomes.

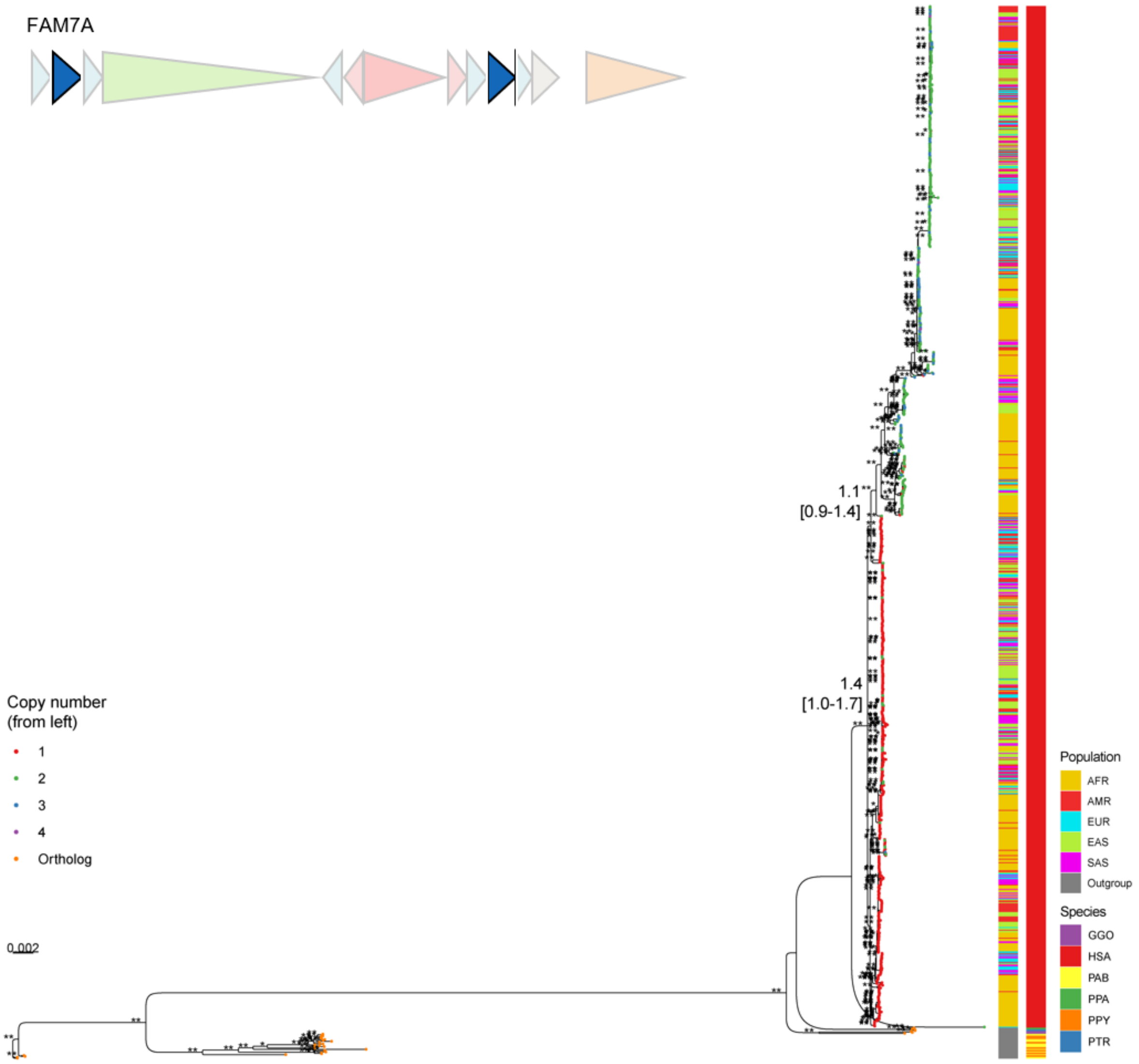


**Supplementary Figure 35. Maximum likelihood phylogenetic tree of the FAM7A duplicon.** The tree includes 1184 sequences with conserved alignment length of 31 kbp. The tree was rooted with the ortholog copy from the Sumatran orangutan (PAB) genome. In addition, the tree also includes ape orthologs from diploid assemblies of chimpanzee (PTR), bonobo (PPA), gorilla (GGO) and Bornean orangutan (PPY). Bootstrap support was indicated by ** (>95) and * (>90). The average divergence times are indicated along with the confidence interval from 100 replicates below. From the 5′ the copy number was counted and indicated by different color of tip nodes. Ancestries of human haplotypes are indicated on the right as well as the species codes for nonhuman genomes.

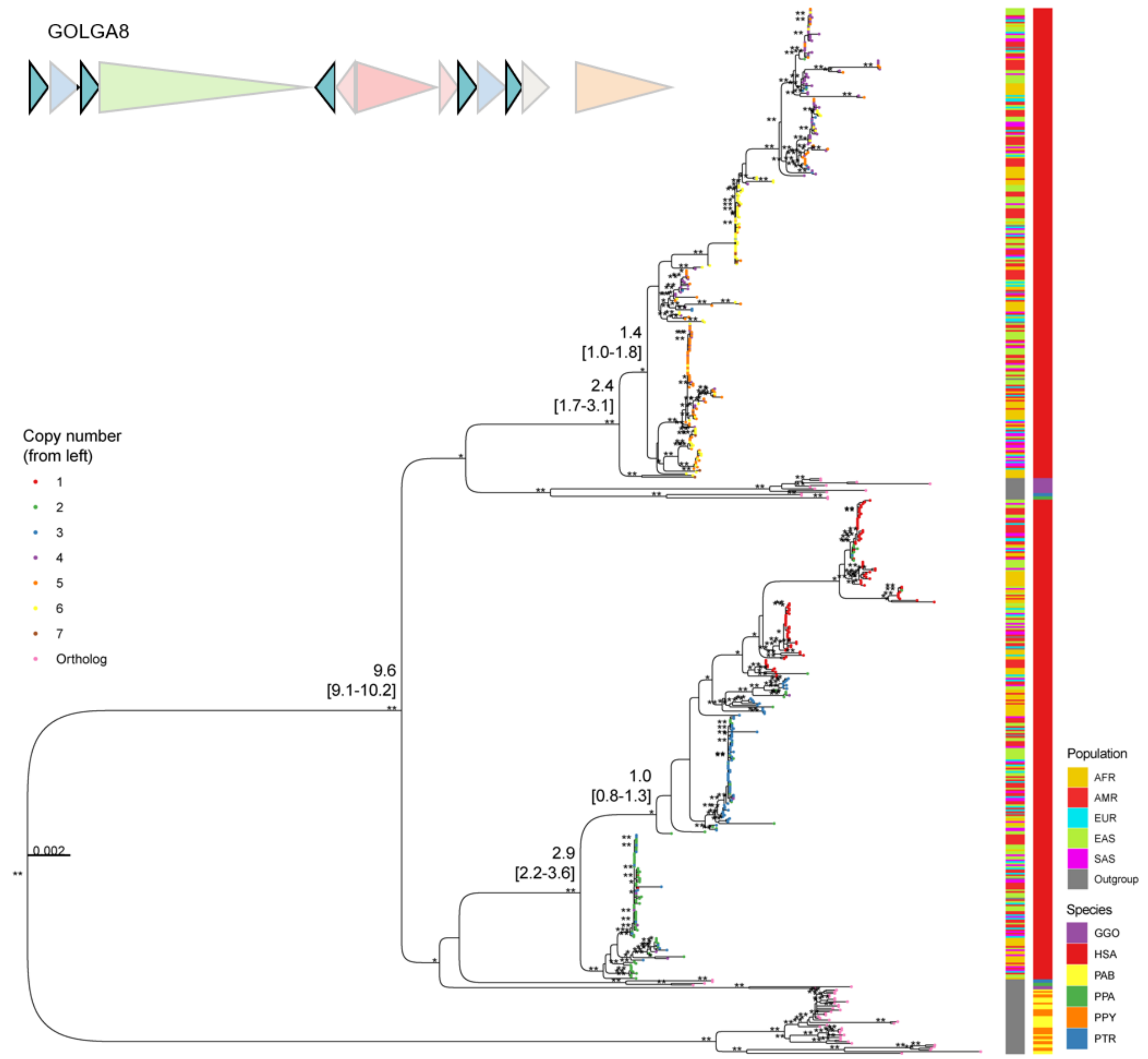


**Supplementary Figure 36. Maximum likelihood phylogenetic tree of the GOLGA8 duplicon.** The tree includes random 630 subset sequences including macaque outgroup sequences with conserved alignment length of 12 kbp. The tree was rooted with the ortholog copies from a macaque genome (not displayed due to long branch length). In addition, the tree also includes ape orthologs from diploid assemblies of chimpanzee (PTR), bonobo (PPA), gorilla (GGO), Bornean orangutan (PPY), and Sumatran orangutan (PAB). Bootstrap support was indicated by ** (>95) and * (>90). The average divergence times are indicated along with the confidence interval from 100 replicates below. From the 5′ the copy number was counted and indicated by different color of tip nodes. Ancestries of human haplotypes are indicated on the right as well as the species codes for nonhuman genomes.

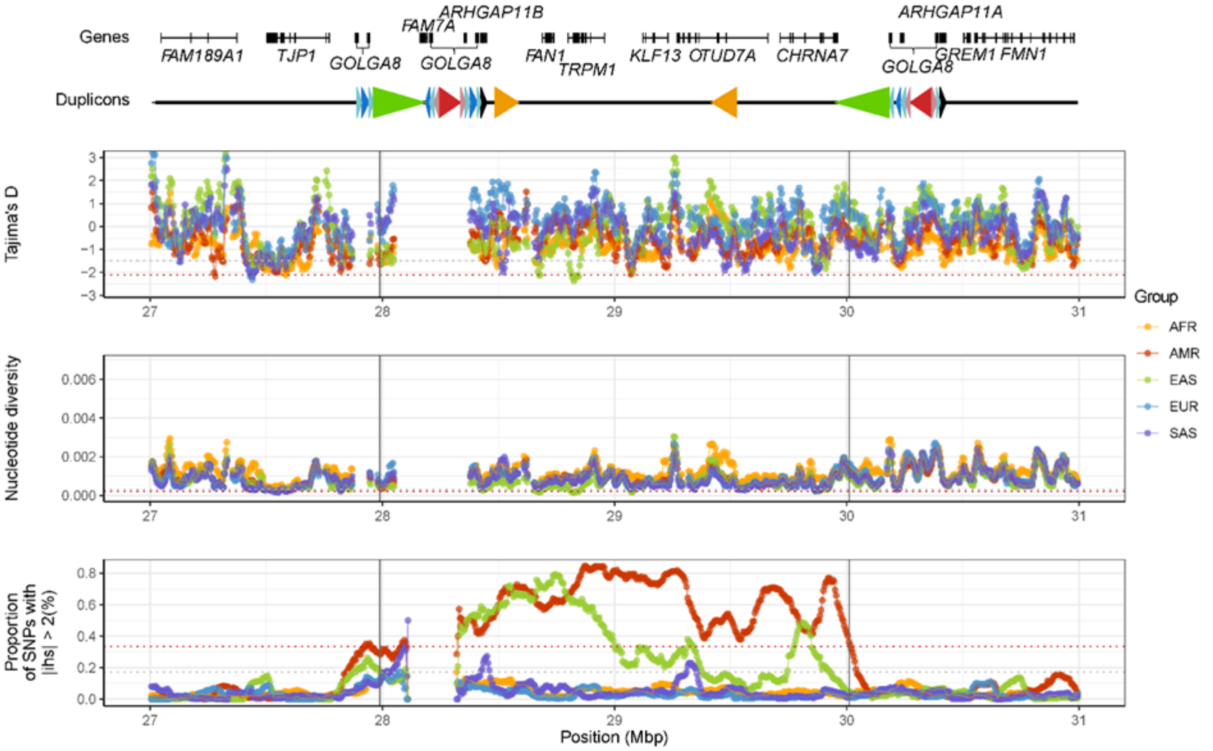


**Supplementary Figure 37. Signatures of selection at the 15q13 locus.** From the top, location of genes followed by duplicons and population genetics statistics are shown. Statistics include Tajima's D, nucleotide diversity (pi), and extended haplotypes (iHS). These are computed from 281 unrelated human genomes from Human Genome Structural Variation Consortium (HGSVC) release 3 and Human Pangenome Reference Consortium (HPRC) release 2 (89 AFR, 50 AMR, 60 EAS, 38 EUR and 44 SAS). For Tajima's D and nucleotide diversity, each dot indicates the statistics computed for 20 kbp bins, with the sliding window of 5 kbp. For iHS statistics, proportion of single-nucleotide substitution variants, with greater than |iHS| > 2 are indicated for the 100 kbp bins with a sliding window of 5 kbp. Red and gray dotted horizontal lines indicate top 1 and 5% of the statistics, respectively, observed in Chromosome 15 of the EAS continental group.

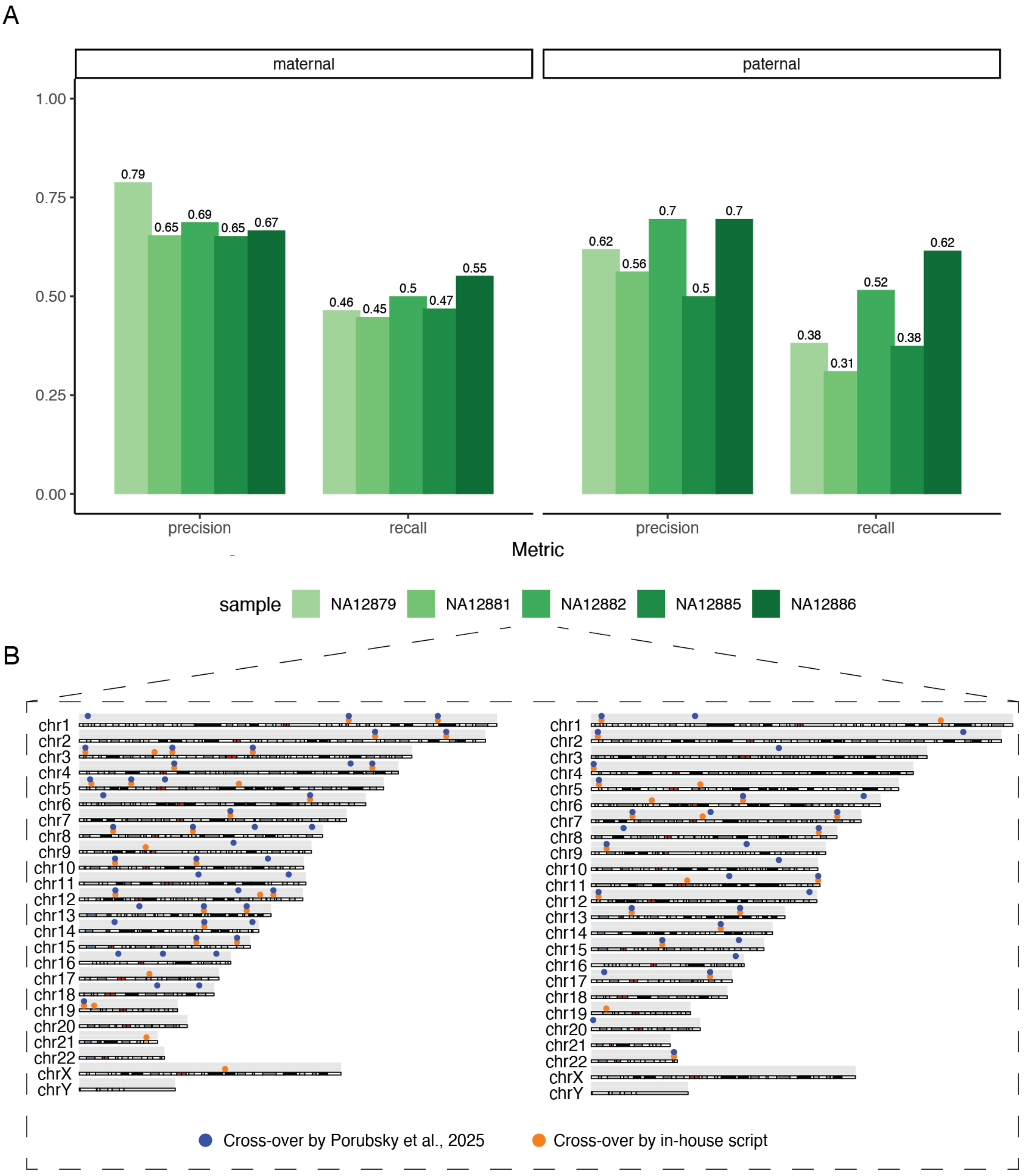


**Supplementary Figure 38. Performance of the Whatshap-based genome-wide crossover analysis** A) Precision and recall for five samples from the "G3" generation from the Platinum Pedigree Project [4]. B) Ideogram showing genomic positions of crossover events in sample NA12882 (left: maternal, right: paternal haplotype).

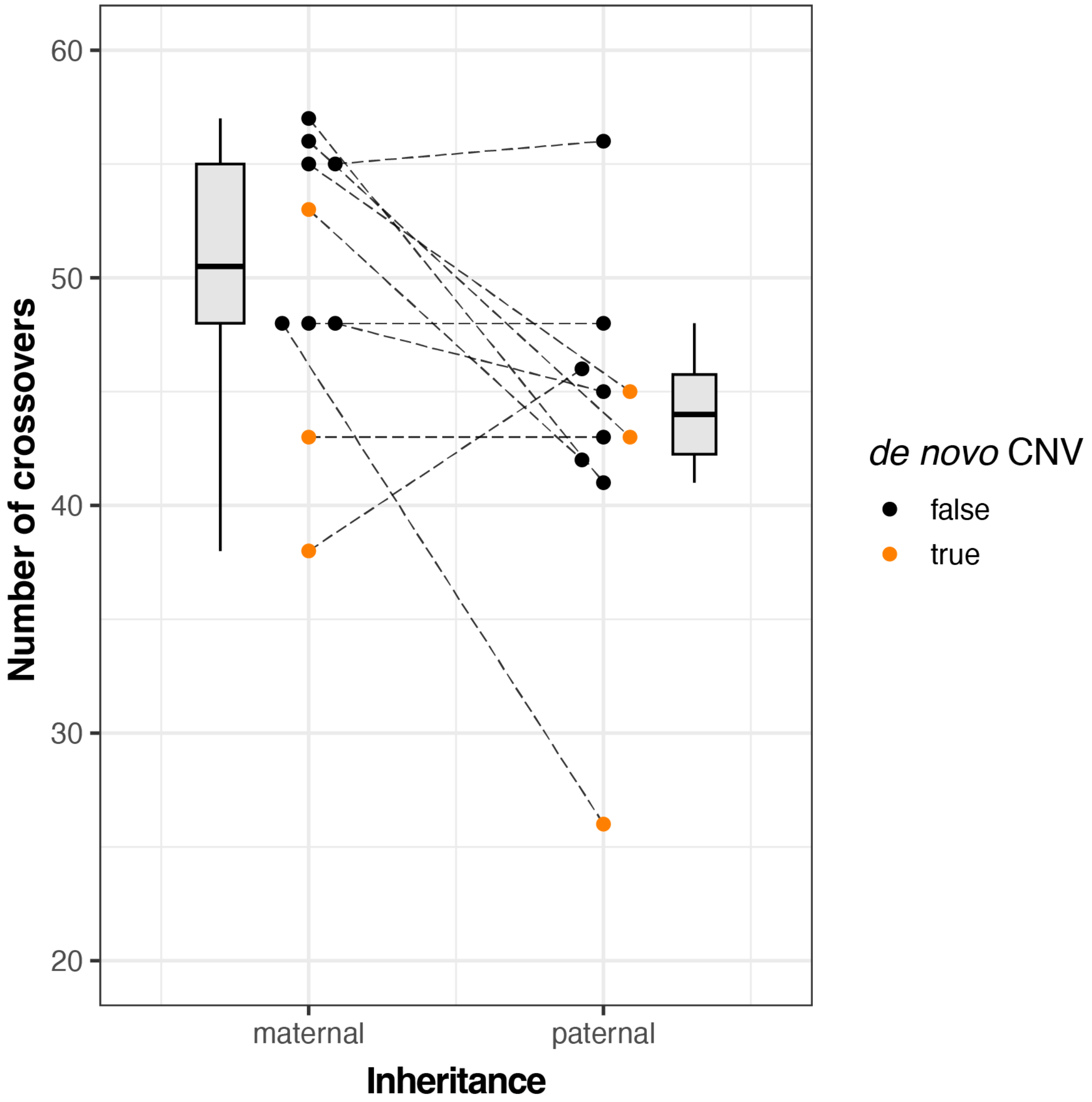


**Supplementary Figure 39. Number of genome-wide crossovers in index patients.** Connected scatterplot indicating the number of inferred genome-wide crossovers on the maternal and paternal alleles of index patients. Alleles carrying a de novo CNV (T1-T4, T7) are indicated in orange. Alleles carrying a de novo CNV do not show significantly increased recombination rates. One paternal outlier (T1) with particularly low crossovers was observed.

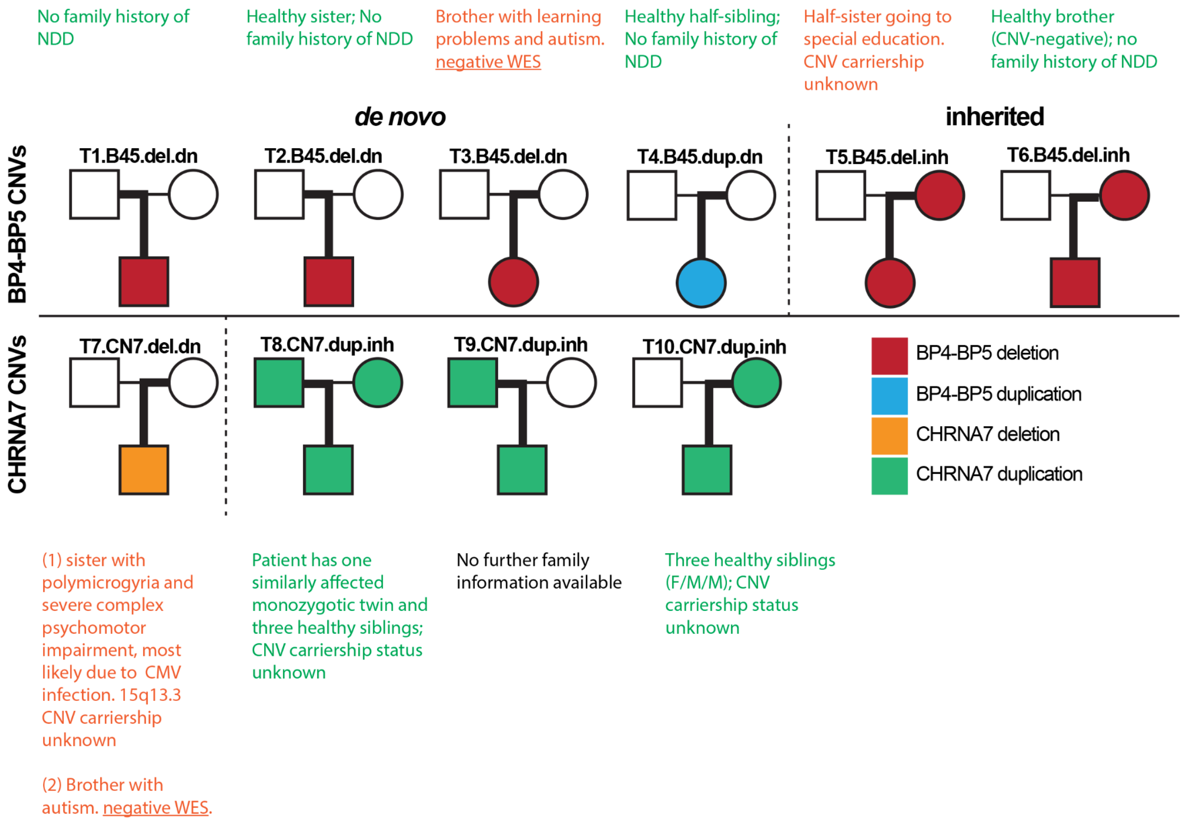


**Supplementary Figure 40. Overview of known cases of 15q13.3 CNVs and -associated phenotypes in the families of T1-T10 beyond the samples directly analysed in this study.** Six families have no known history of NDD (green text), three have additional family members affected by either 15q13.3 CNVs or associated phenotypes (red text). No further information could be obtained for T9 (black text).

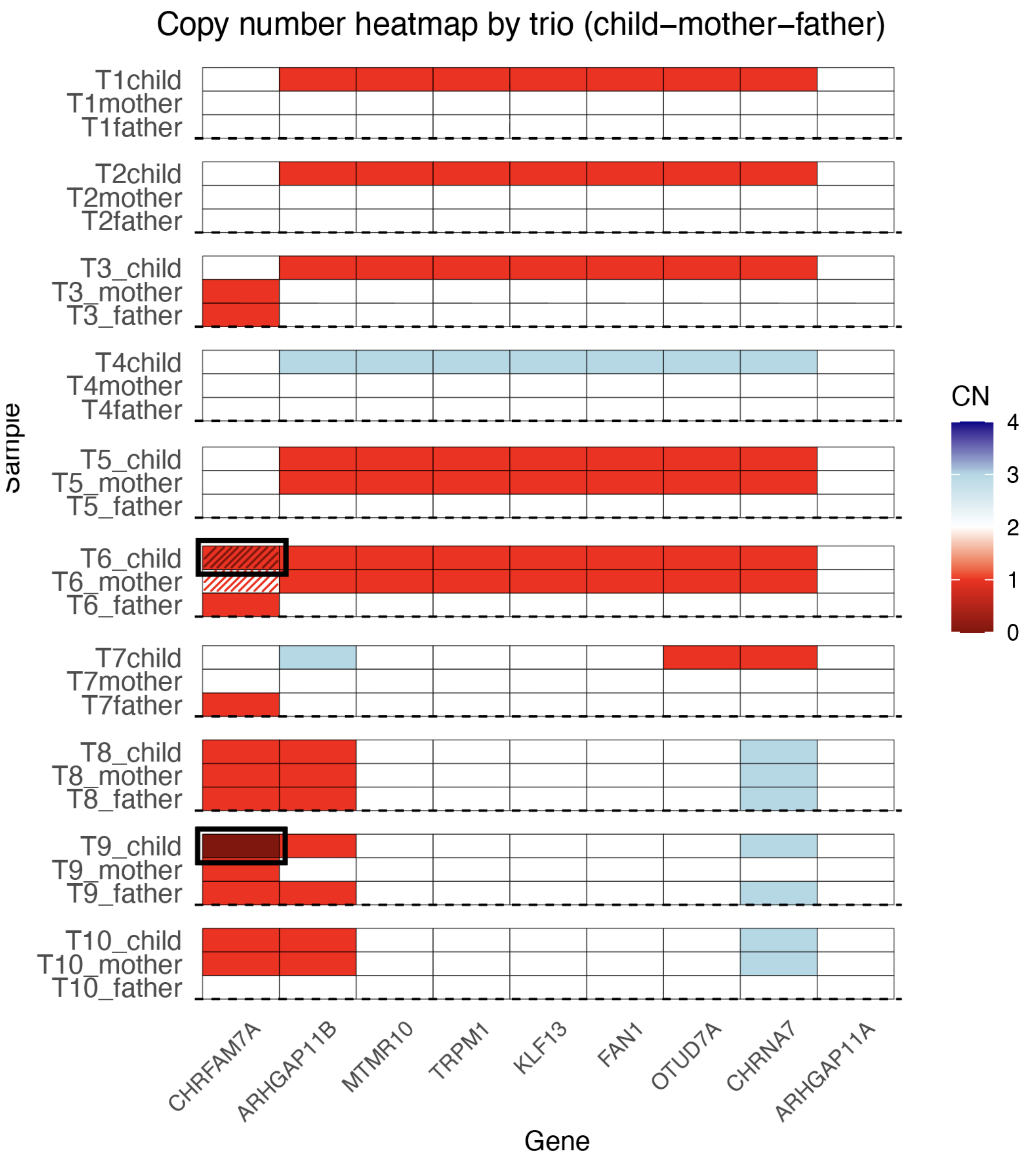


**Supplementary Figure 41. Copy-numbers of 15q13.3 genes in all patients and their parents.** In the case of T6, diagonal stripes indicate that the gene is present, but may be functionally affected due to a breakpoint overlapping the region directly upstream (See Supplementary Figure 9C). Instances where the index inherited copy-number variants from *both* parents are highlighted by black boxes. In both cases, the index inherited a *CHRFAM7A* deletion from one parent, additionally to the CNV inherited from the other parent. In case of T9, this leads to a homozygous deletion of *CHRFAM7A*, which may contribute to disease severity, particularly in conjunction with the duplication of *CHRNA7*.

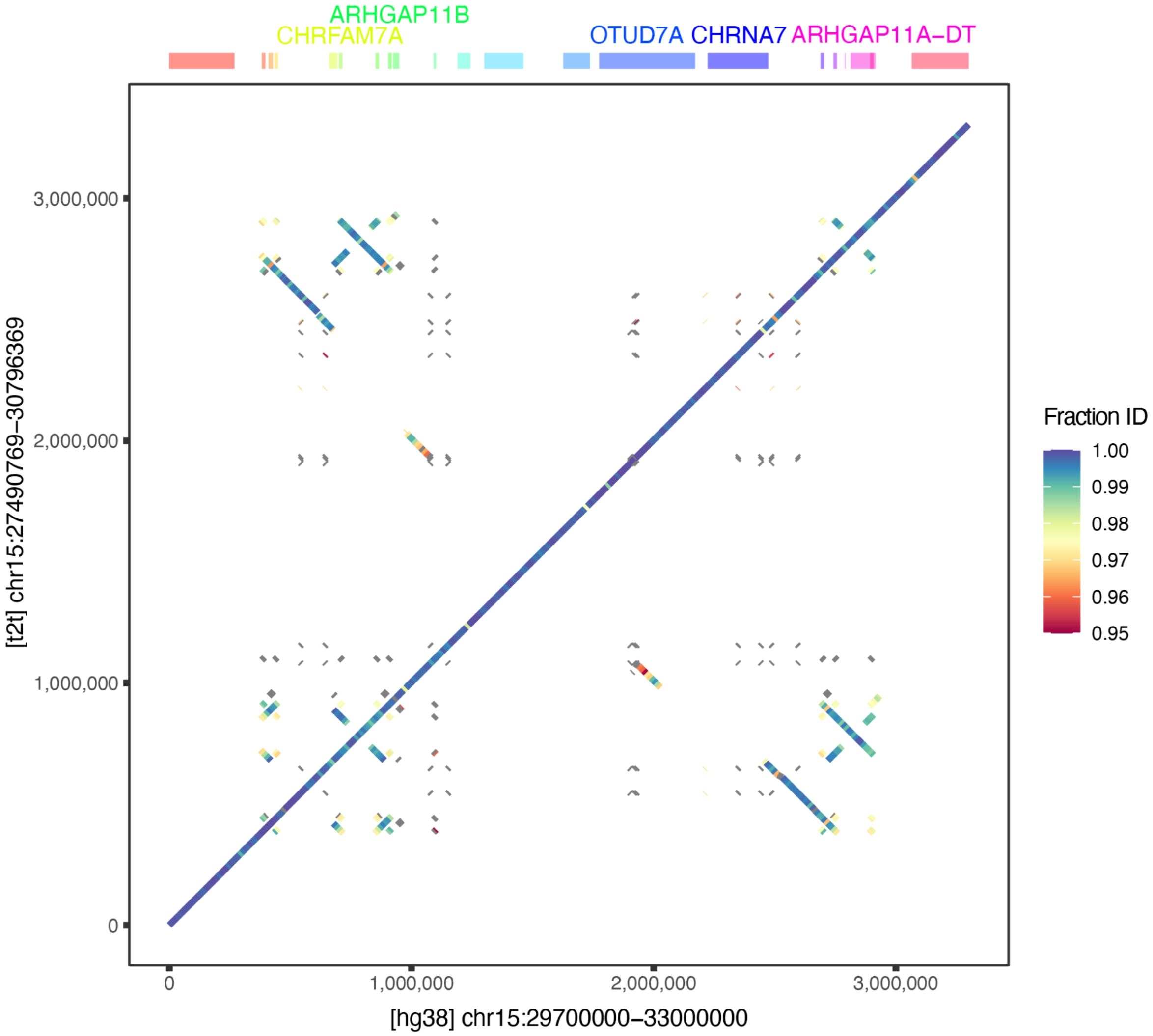


**Supplementary Figure 42. Dotplot comparing the 15q13.3 region in the GRCh38 and T2T-v2.0 genome assemblies.** Colored segments indicate high-similarity (>=95%) stretches of at least 1 kbp in size, with the color indicating the degree of similarity. The uninterrupted diagonal indicates an absence of SVs > 10 kbp between the two assemblies. Off-diagonal segments correspond to repeats or segmental duplications.

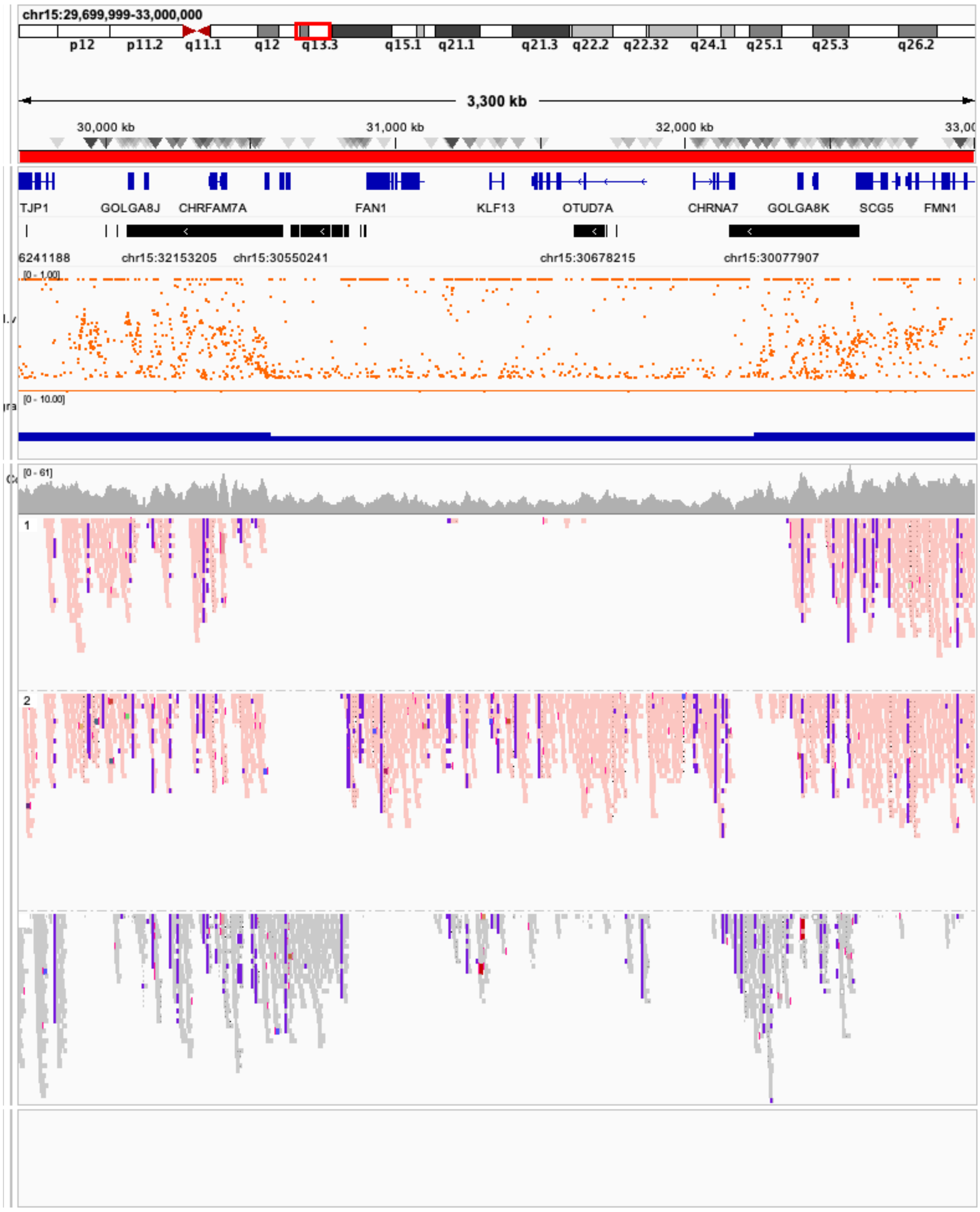


**Supplementary Figure 43. Visualization of aligned sequencing reads and derived metrics of the CNV-affected index patient of T1 (de novo BP4-BP5 Deletion).** Visualized region: [HG38:29,000,000-33,000,000]. Tracks from top to bottom: Refseq gene annotations (blue), Segmental duplications annotations (black; visible are the BP4, CHRNA7-LCR and BP5 blocks), b-allele frequency of SNVs (orange), HifiCNV-based copy-number estimate track (blue), read coverage (grey), Pedigree-phased aligned reads (first read track: paternal haplotype, second: maternal haplotype, third: unphased). The paternal deletion is easily detectable; however breakpoint locations and haplotype architecture can not be readily deduced.

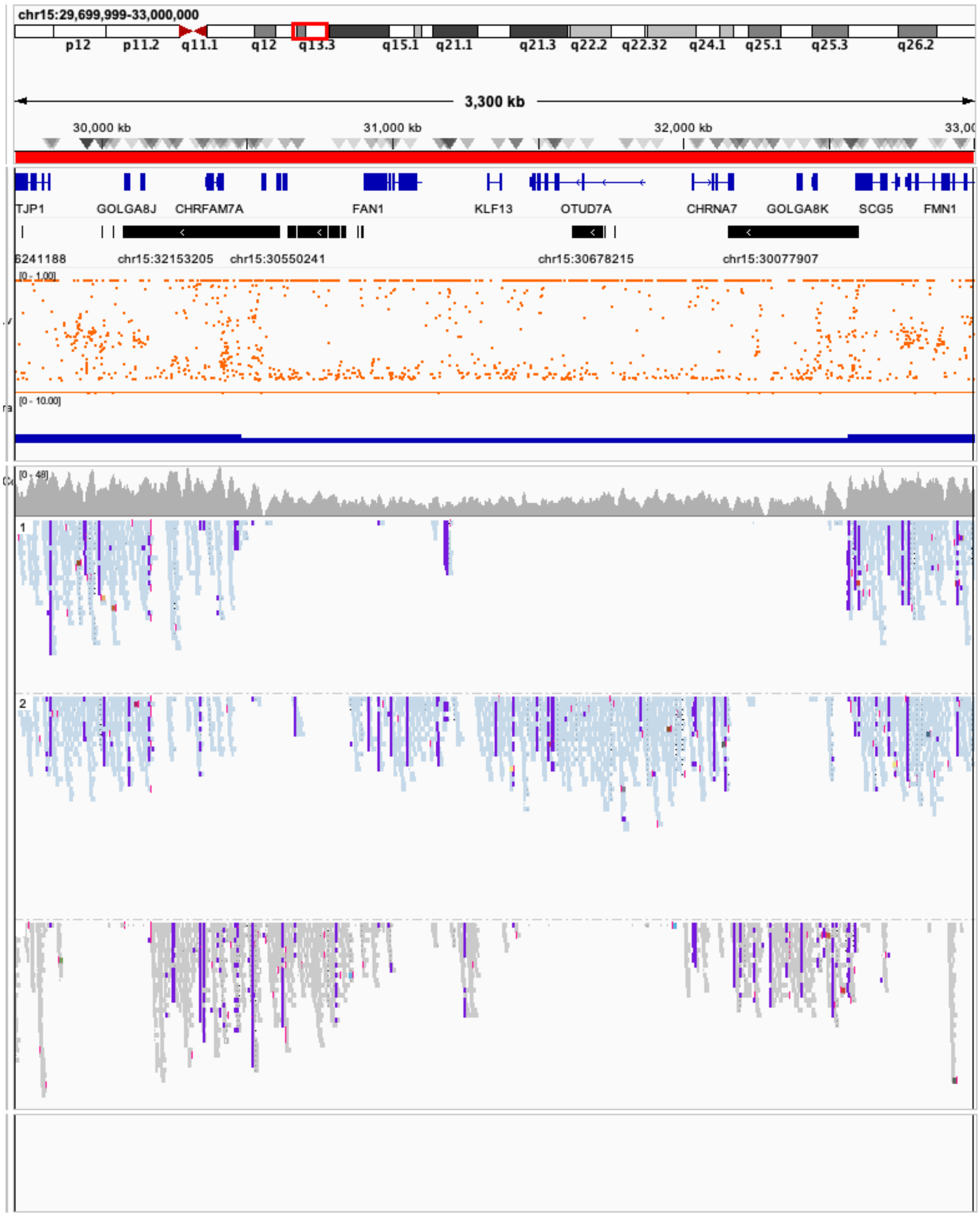


**Supplementary Figure 44. Sequencing reads from the index patient of T2 *(de novo BP4-B*P5 Deletion).**
Track description: see Supplementary Figure 43.

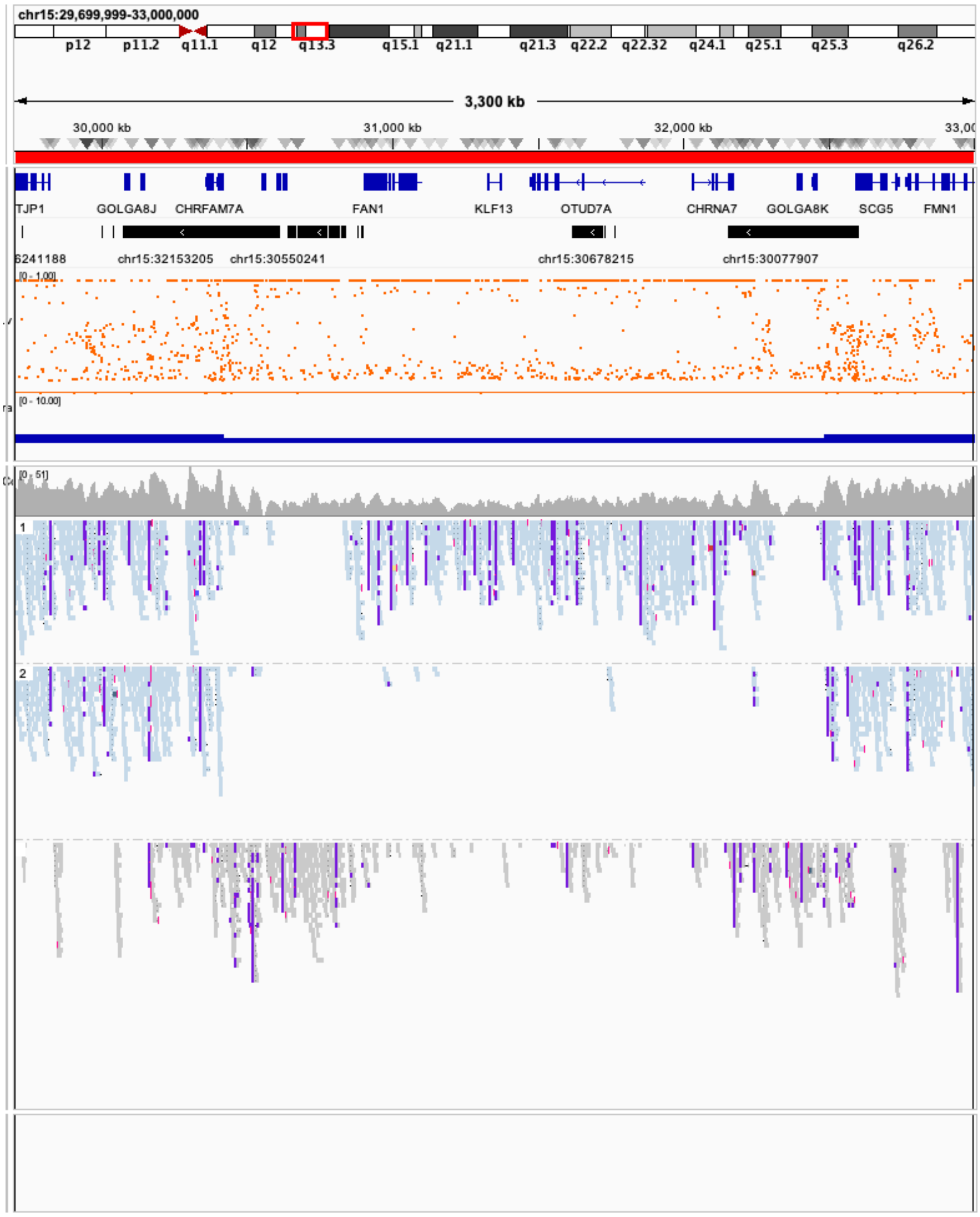


**Supplementary Figure 45. Sequencing reads from the index patient of T3 (de novo BP4-BP5 Deletion).**
Track description: see Supplementary Figure 43.

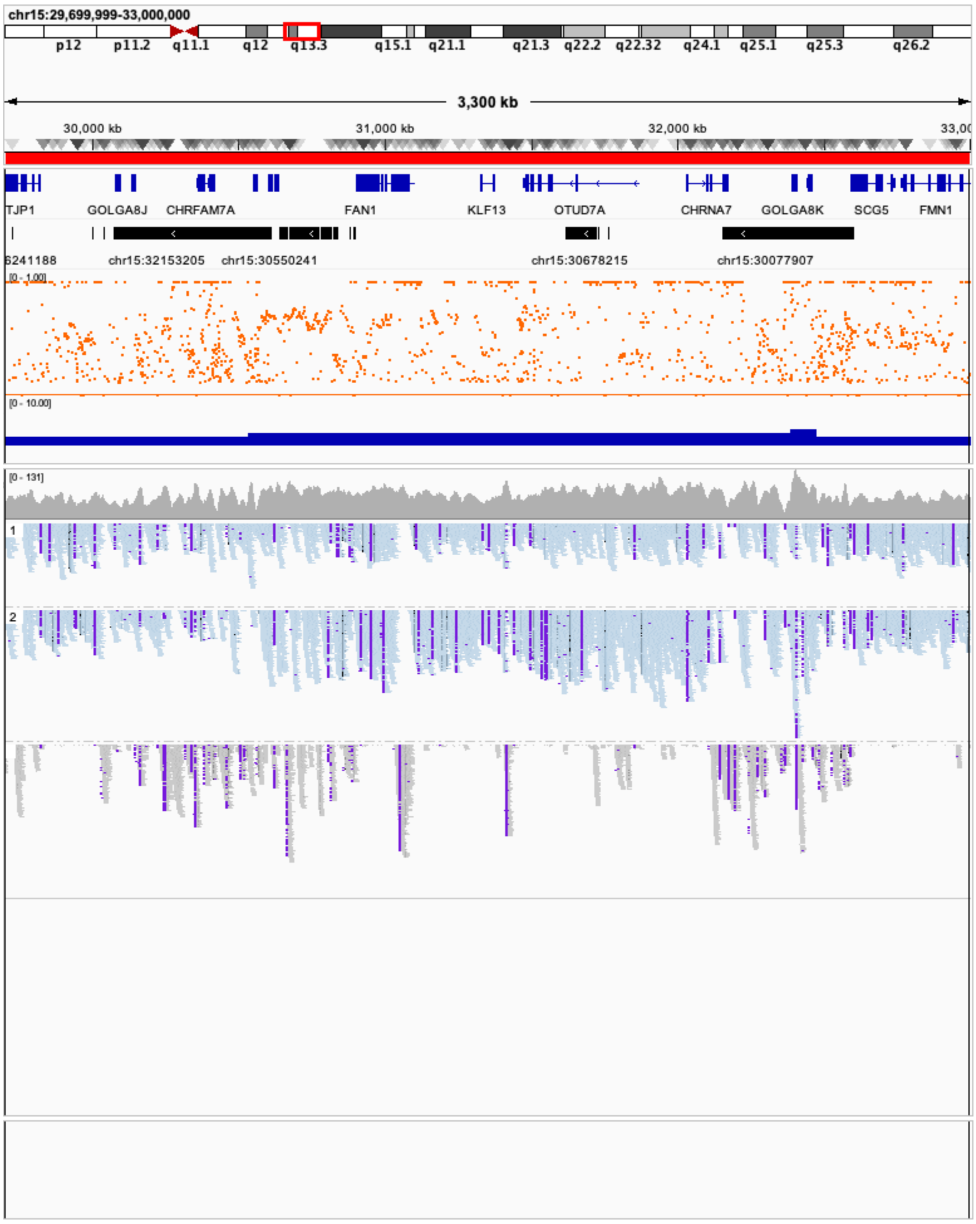


**Supplementary Figure 46. Sequencing reads from the index patient of T4 (de novo BP4-BP5 Duplication).** Track description: see Supplementary Figure 43.

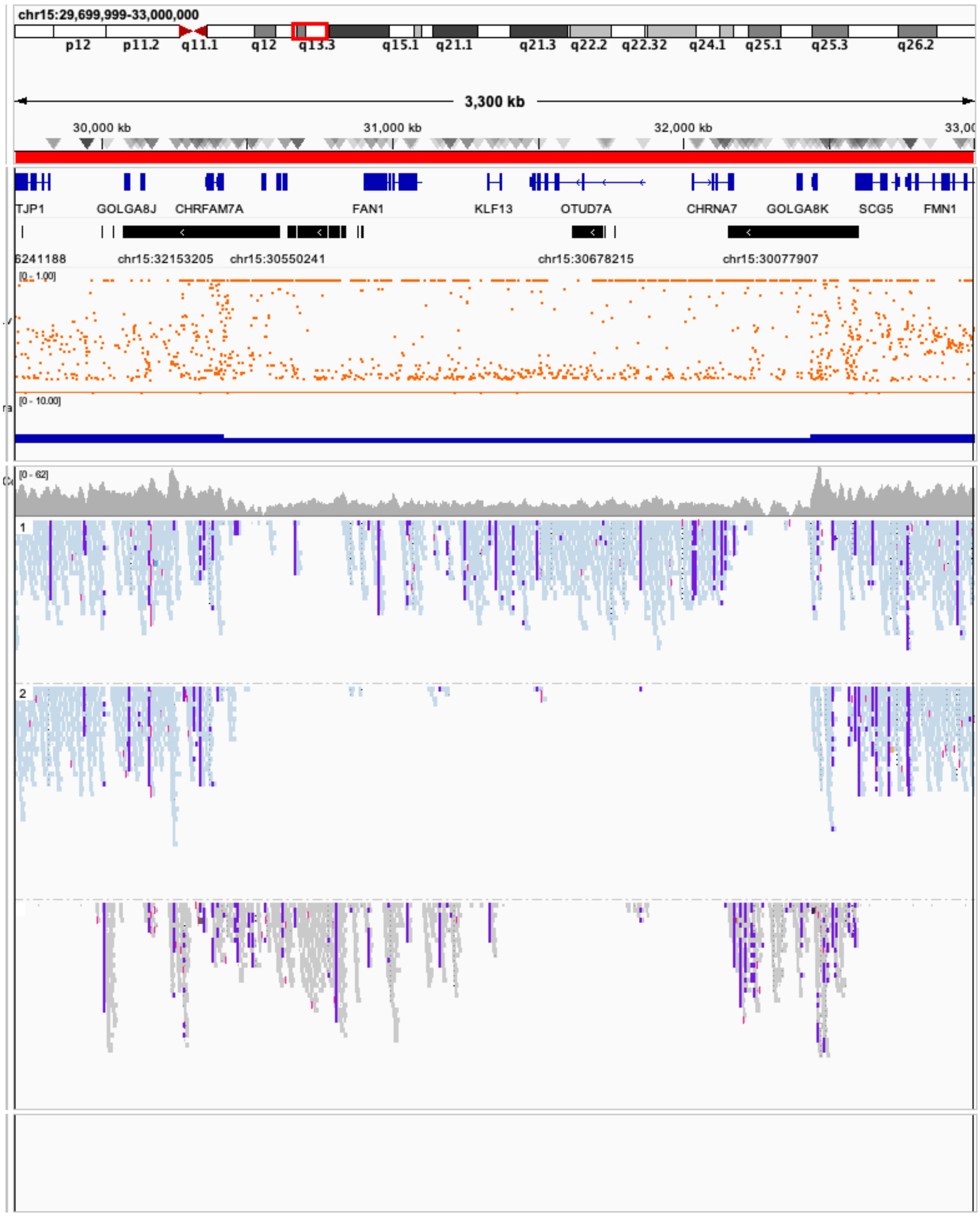


**Supplementary Figure 47. Sequencing reads from the index patient of T5 (inherited BP4-BP5 Deletion).** Track description: see Supplementary Figure 43.

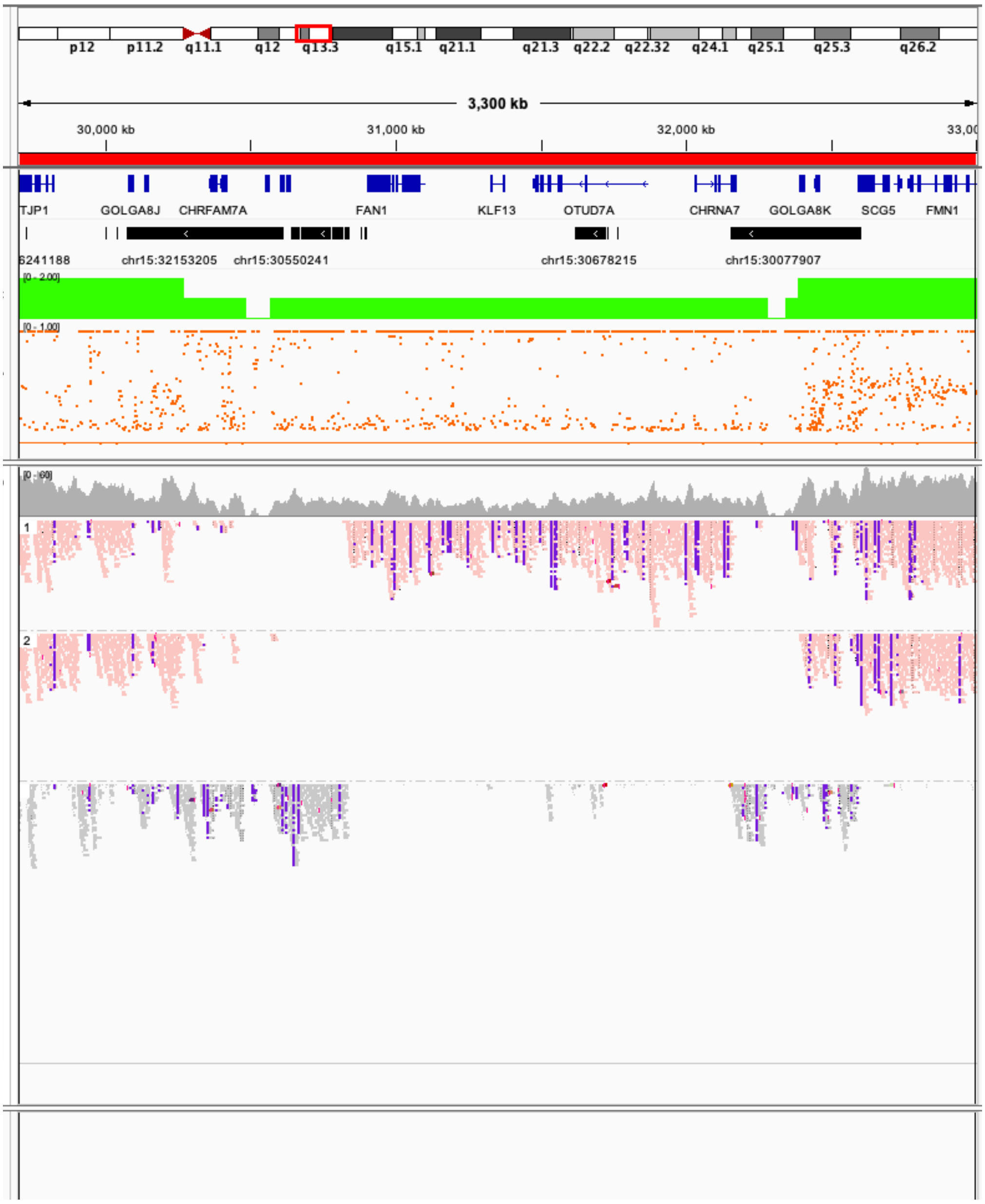


**Supplementary Figure 48. Sequencing reads from the index patient of T6 (inherited BP4-BP5 deletion).**
Track description: see Supplementary Figure 43.

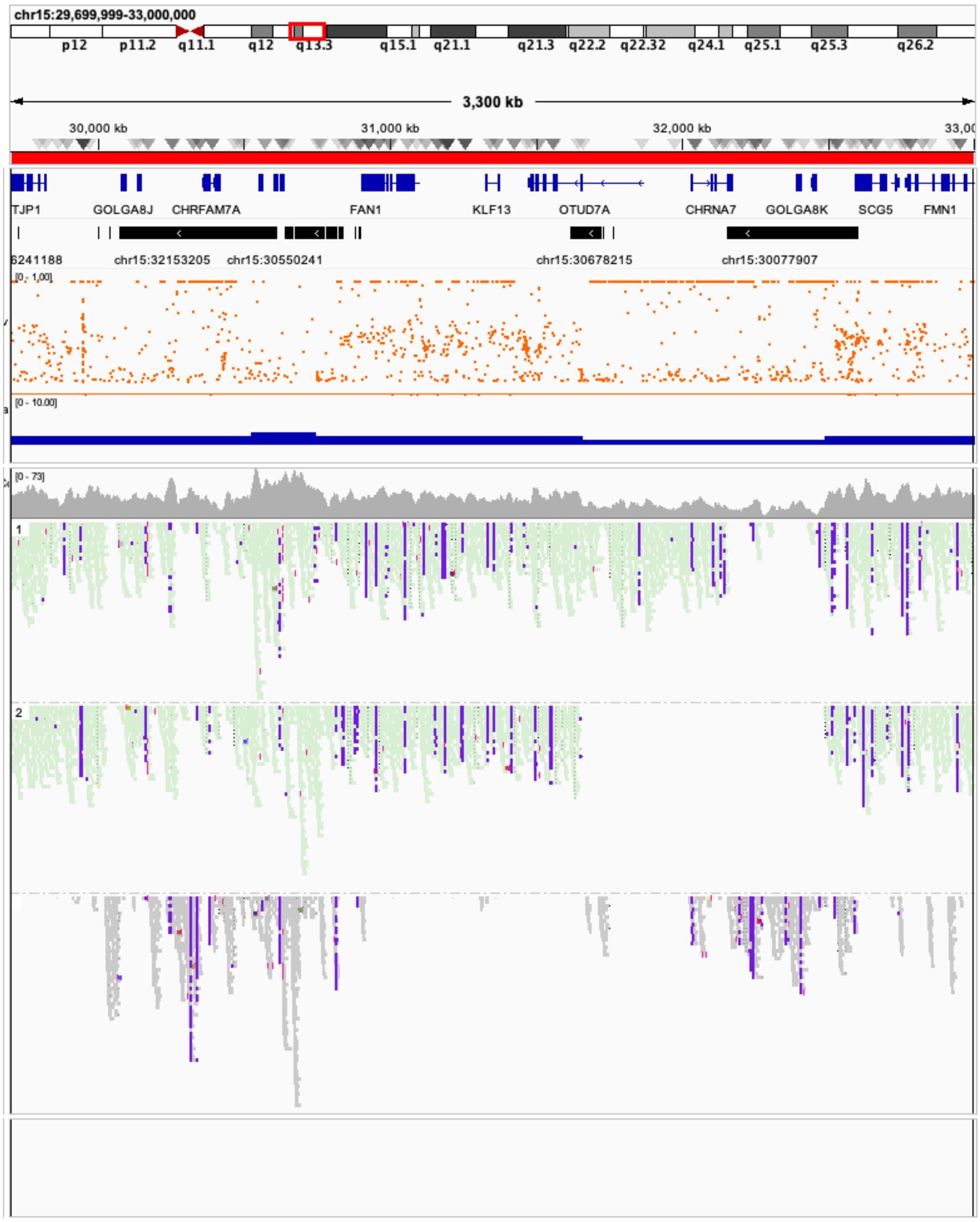


**Supplementary Figure 49. Sequencing reads from the index patient of T7 (de novo *CHRNA7* deletion).** Track description: see Supplementary Figure 43.

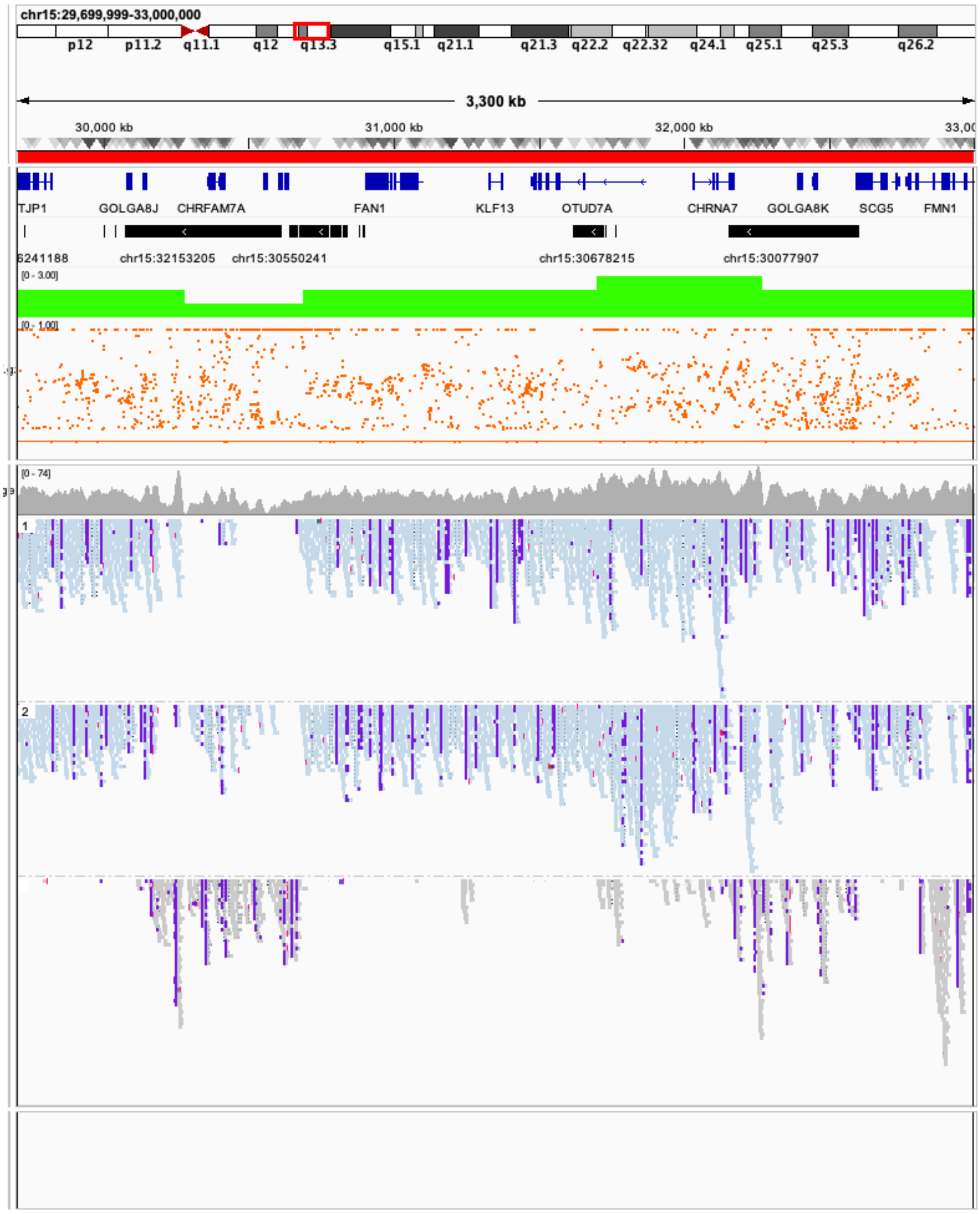


**Supplementary Figure 50. Sequencing reads from the index patient of T8 (inherited *CHRNA7* Duplication).** Track description: see Supplementary Figure 43.

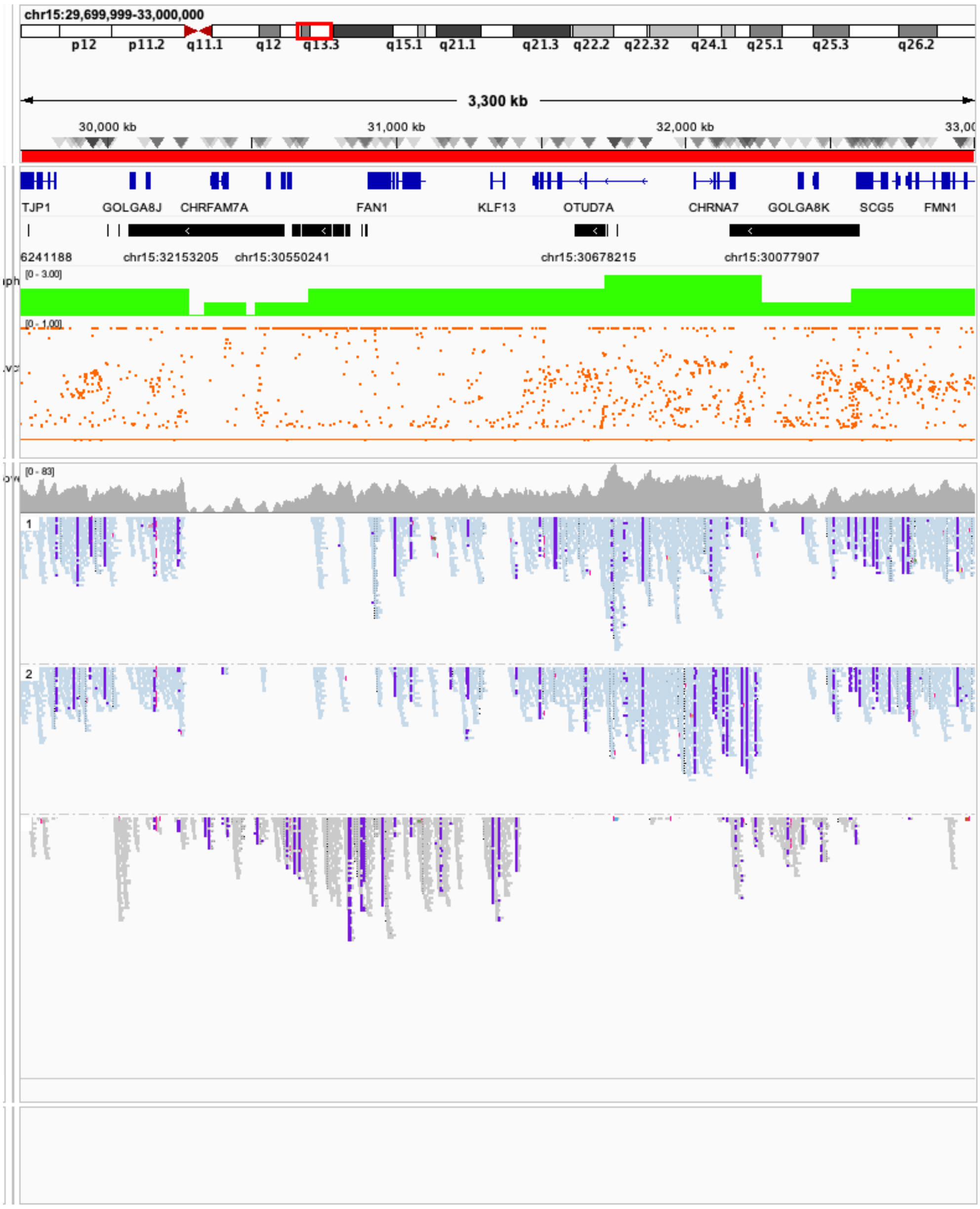


**Supplementary Figure 51. Sequencing reads from the index patient of T9 (inherited *CHRNA7* Duplication).** Track description: see Supplementary Figure 43.

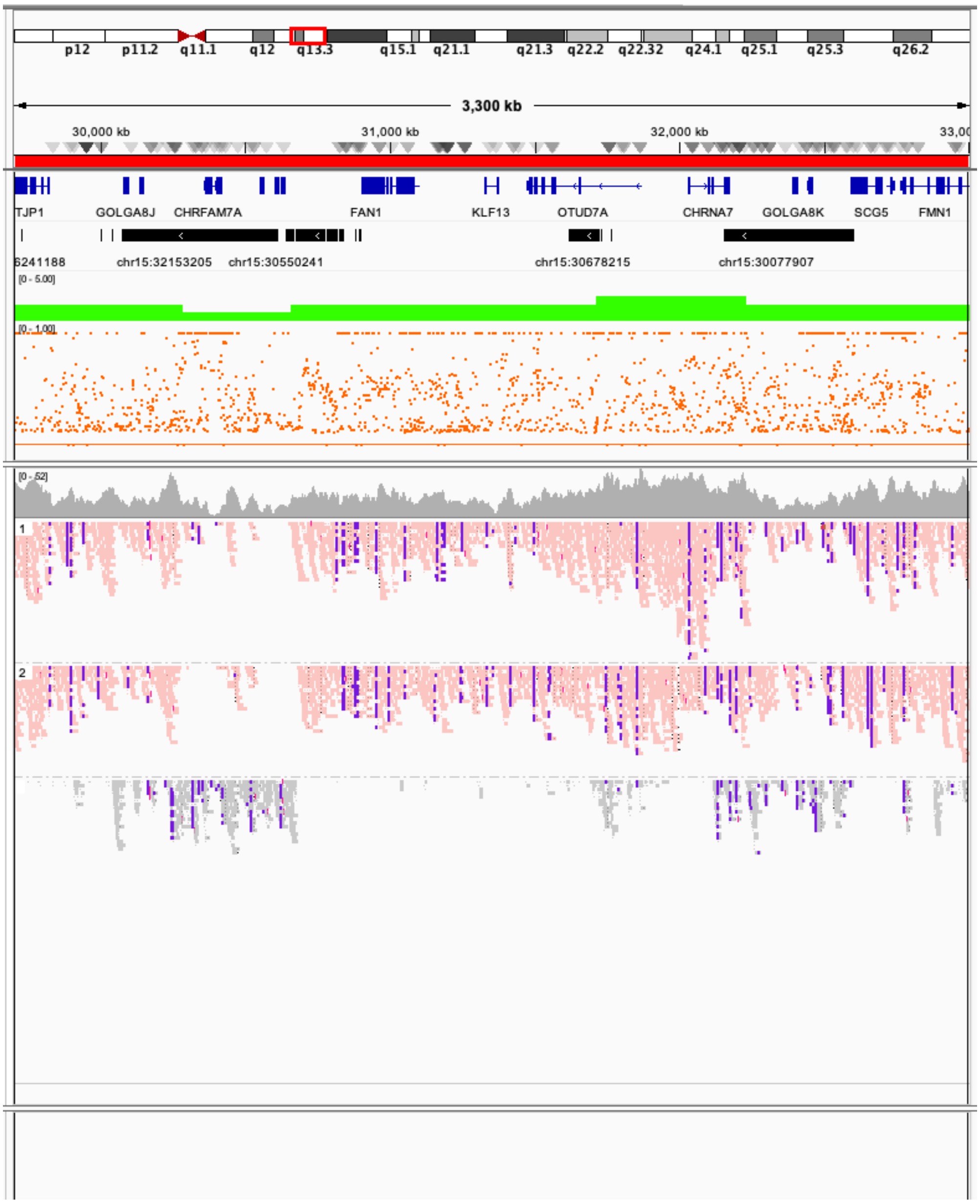


**Supplementary Figure 52. Sequencing reads from the index patient of T10 (inherited *CHRNA7* Duplication).** Track description: see Supplementary Figure 43.

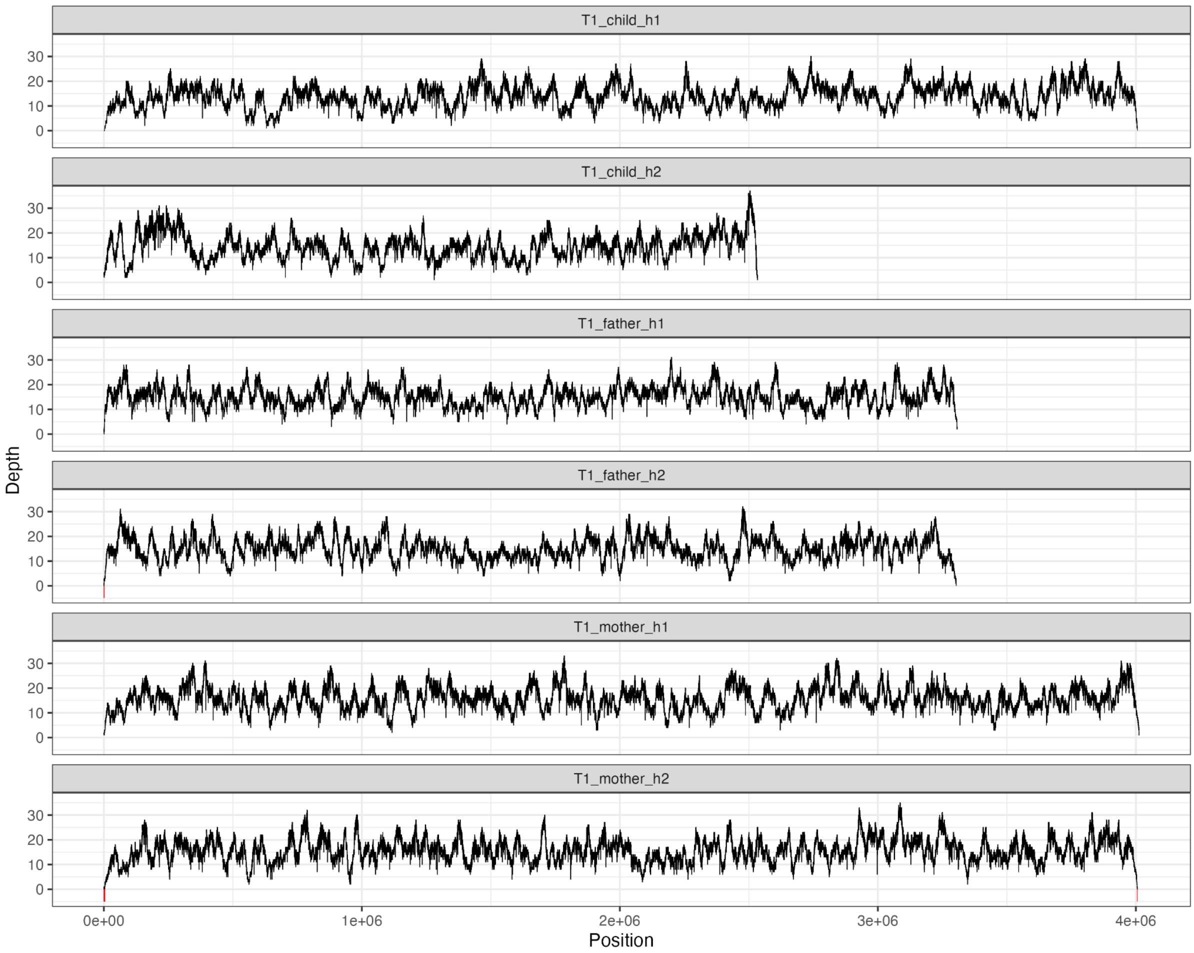


**Supplementary Figure 53. Depth profiles of reads mapping to assembled haplotypes for trio T1.** Reads of each sample were simultaneously mapped against both assembled haplotypes of that sample. Positions not covered by any read (i.e. depth of coverage 0) would indicate potential misassemblies. Such positions are indicated with a red marker under the depth track (see e.g. T1_mother_h2). “depth=0” positions at the beginning and end of assemblies are expected and not indicative of misassemblies.

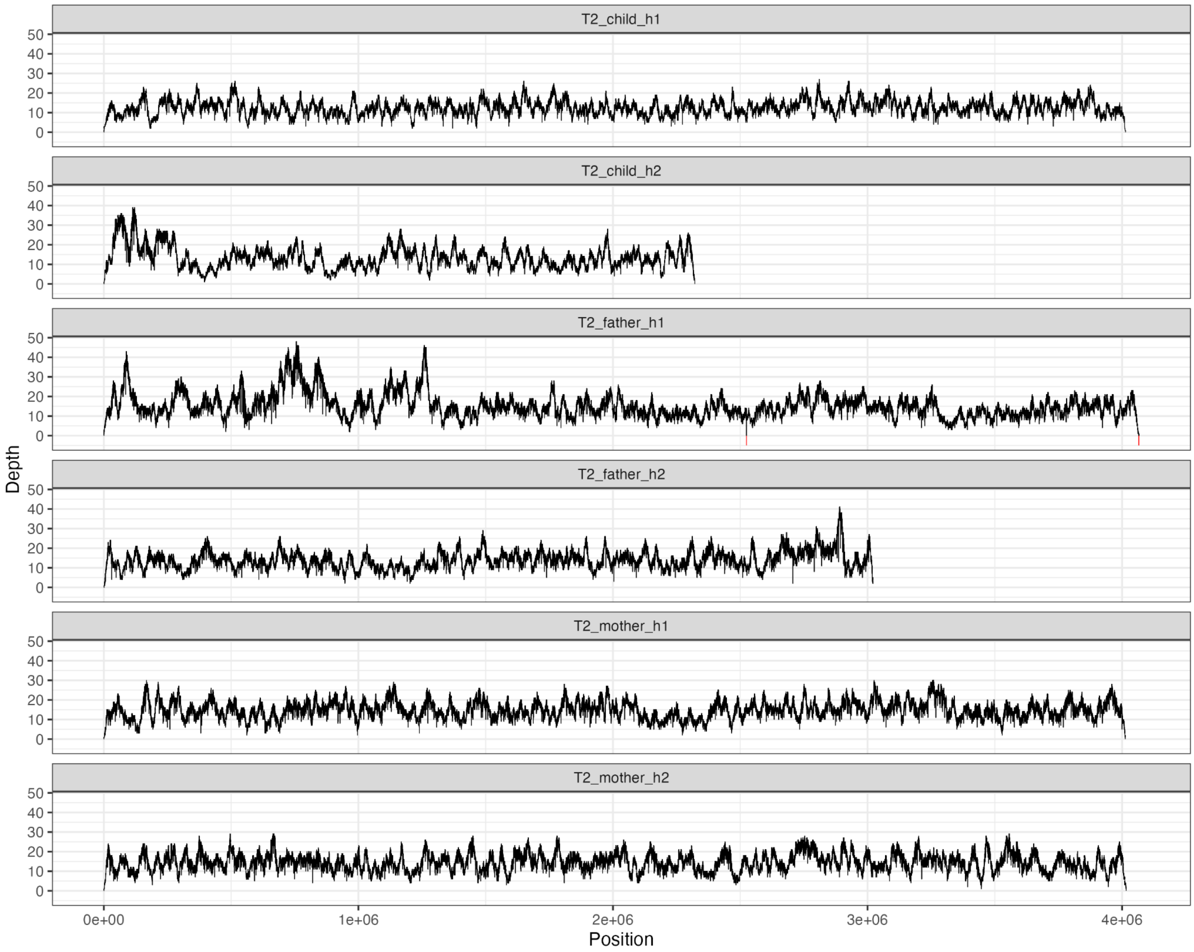


**Supplementary Figure 54. Depth profiles of reads mapping to assembled haplotypes for trio T2.**
Description: See Supplementary Figure 53.

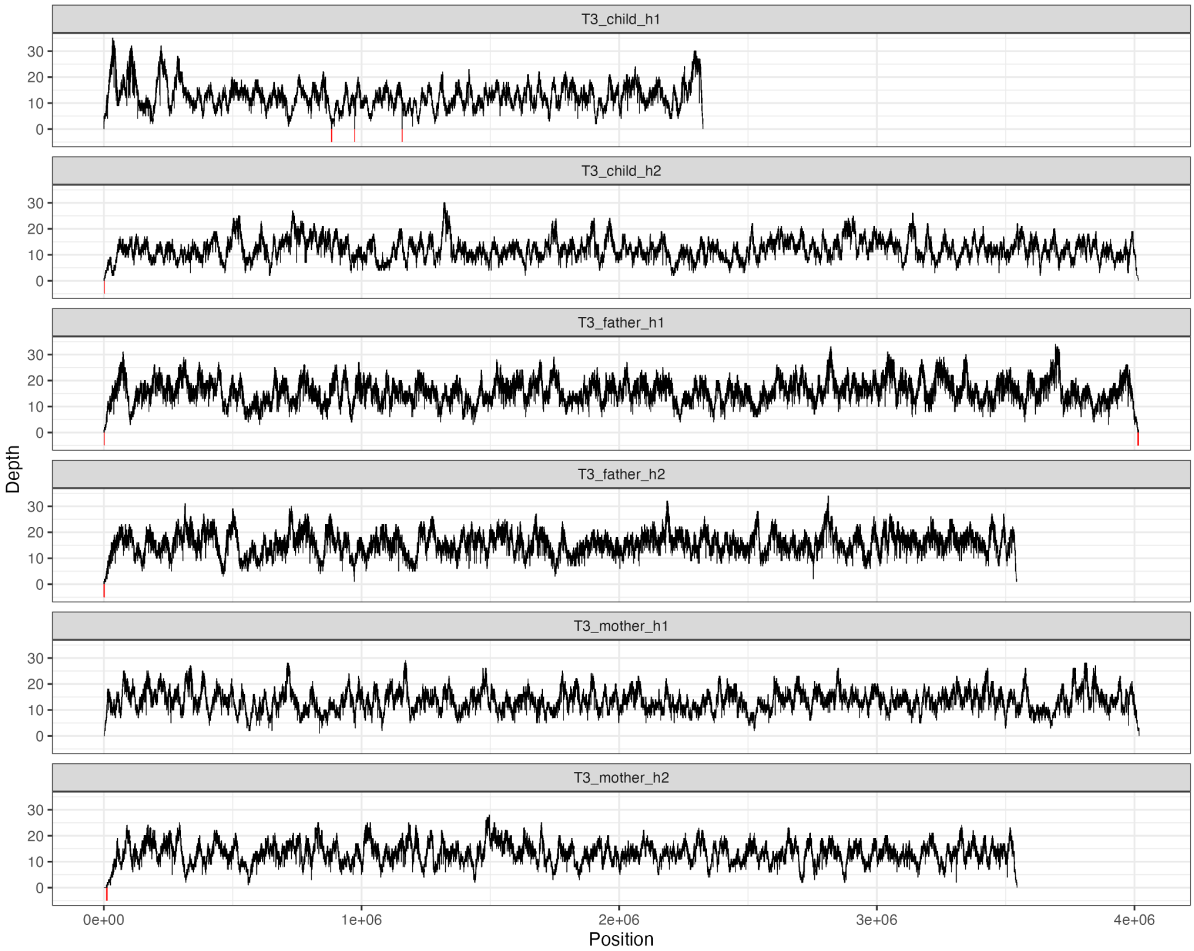


**Supplementary Figure 55**. **Depth profiles of reads mapping to assembled haplotypes for trio T3.**
Description: See Supplementary Figure 53.

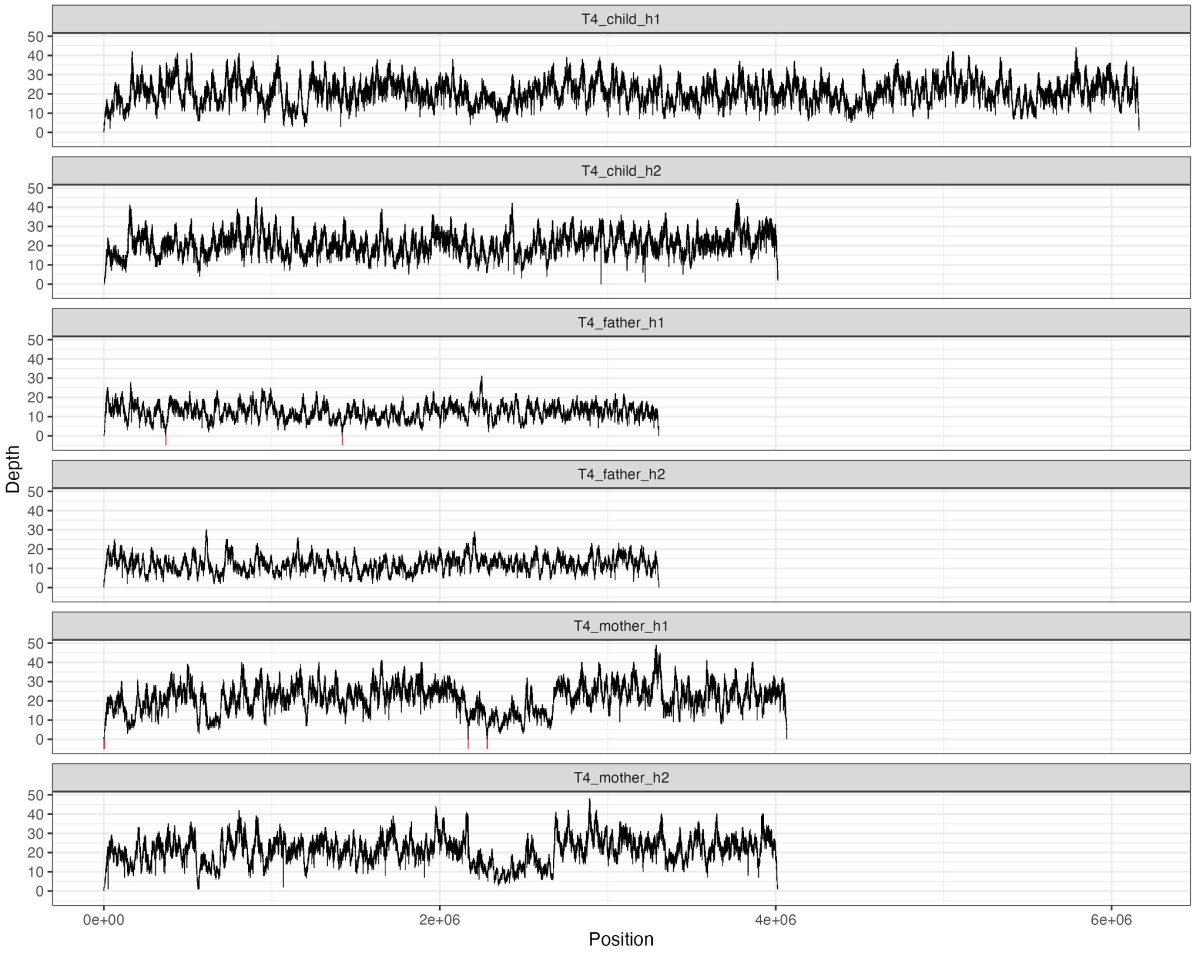


**Supplementary Figure 56. Depth profiles of reads mapping to assembled haplotypes for trio T4.**
Description: See Supplementary Figure 53.

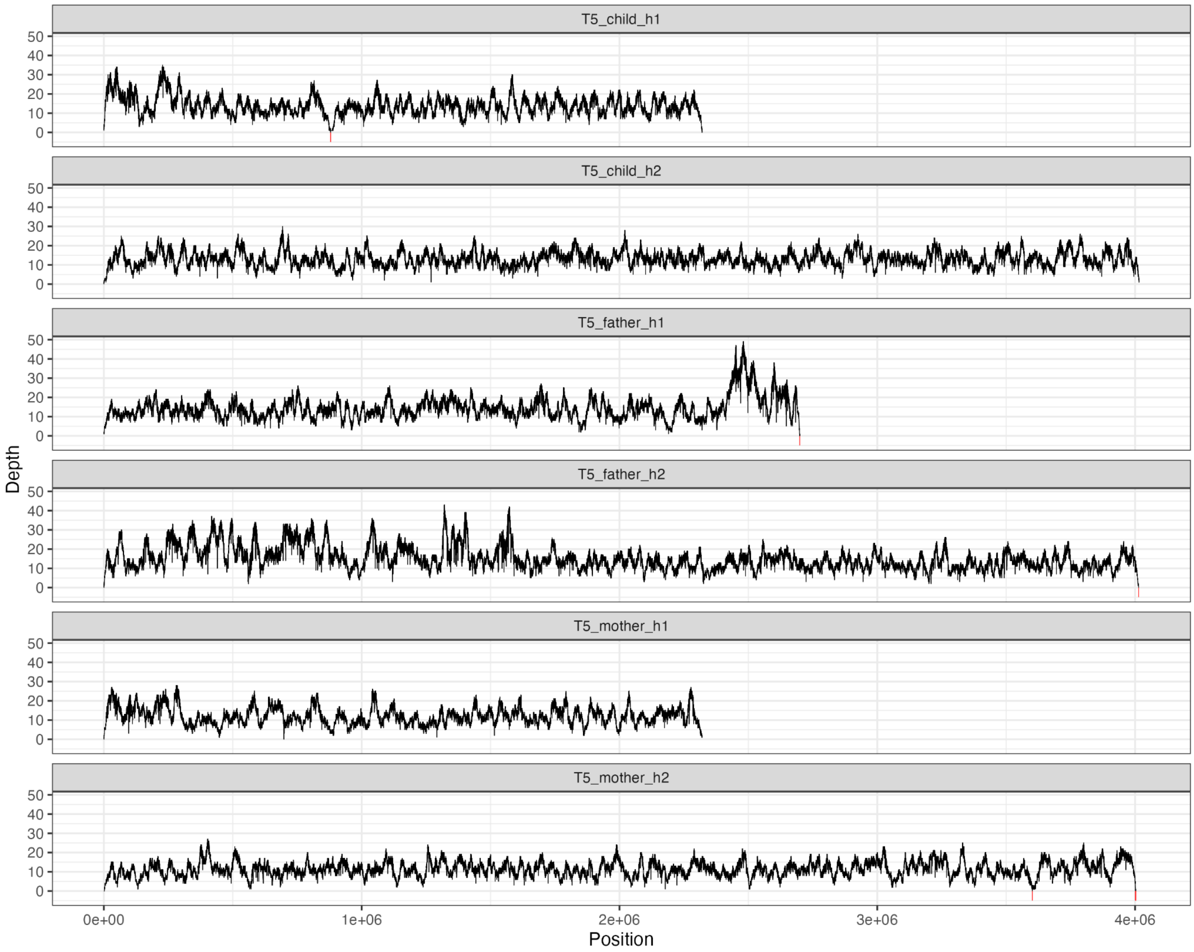


**Supplementary Figure 57. Depth profiles of reads mapping to assembled haplotypes for trio T5.**
Description: See Supplementary Figure 53. T5_father_h1 did not assemble into a contig, producing wavy coverage patterns for these assemblies.

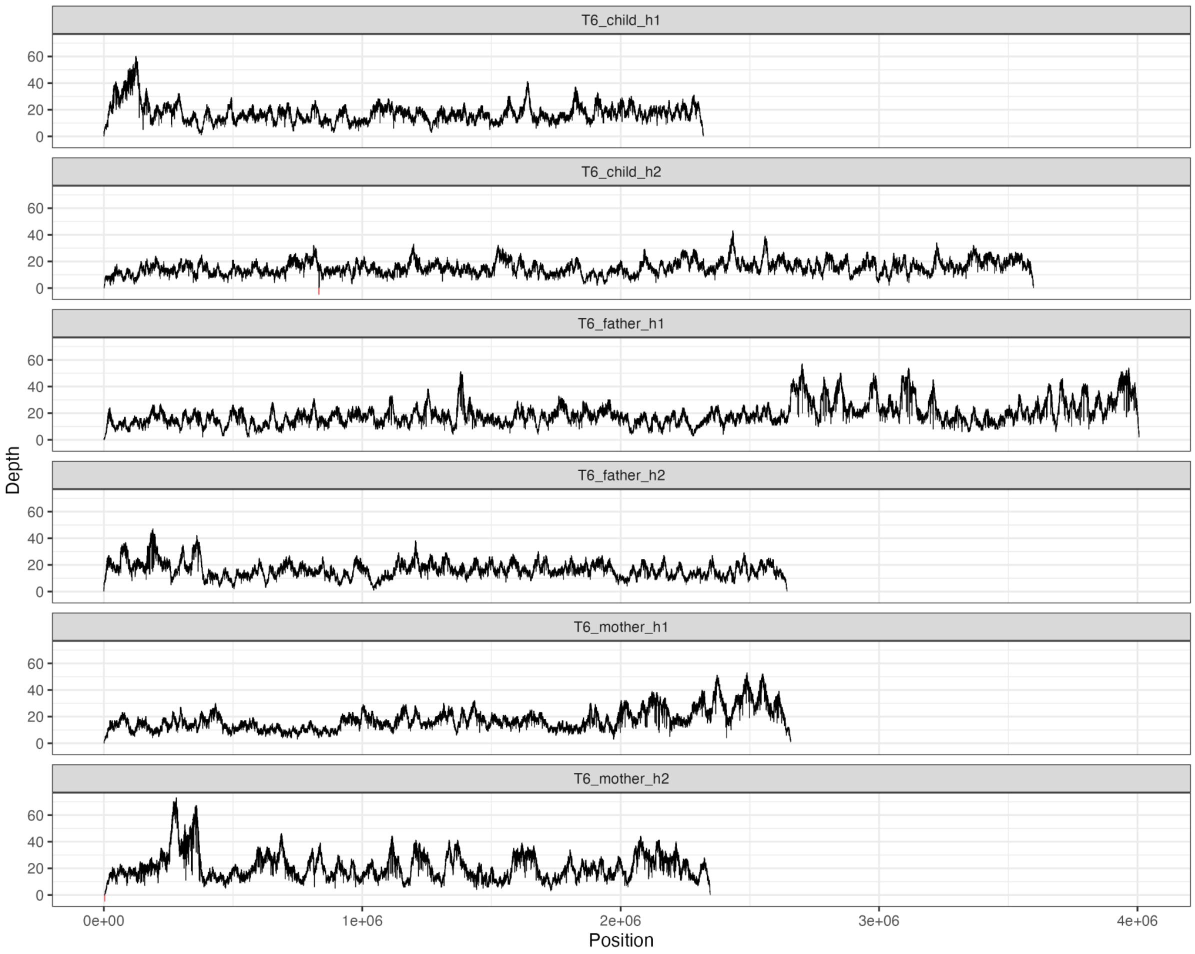


**Supplementary Figure 58**. **Depth profiles of reads mapping to assembled haplotypes for trio T6.** Description: See Supplementary Figure 53. Both parents were incompletely assembled, producing wavy coverage patterns for these samples.

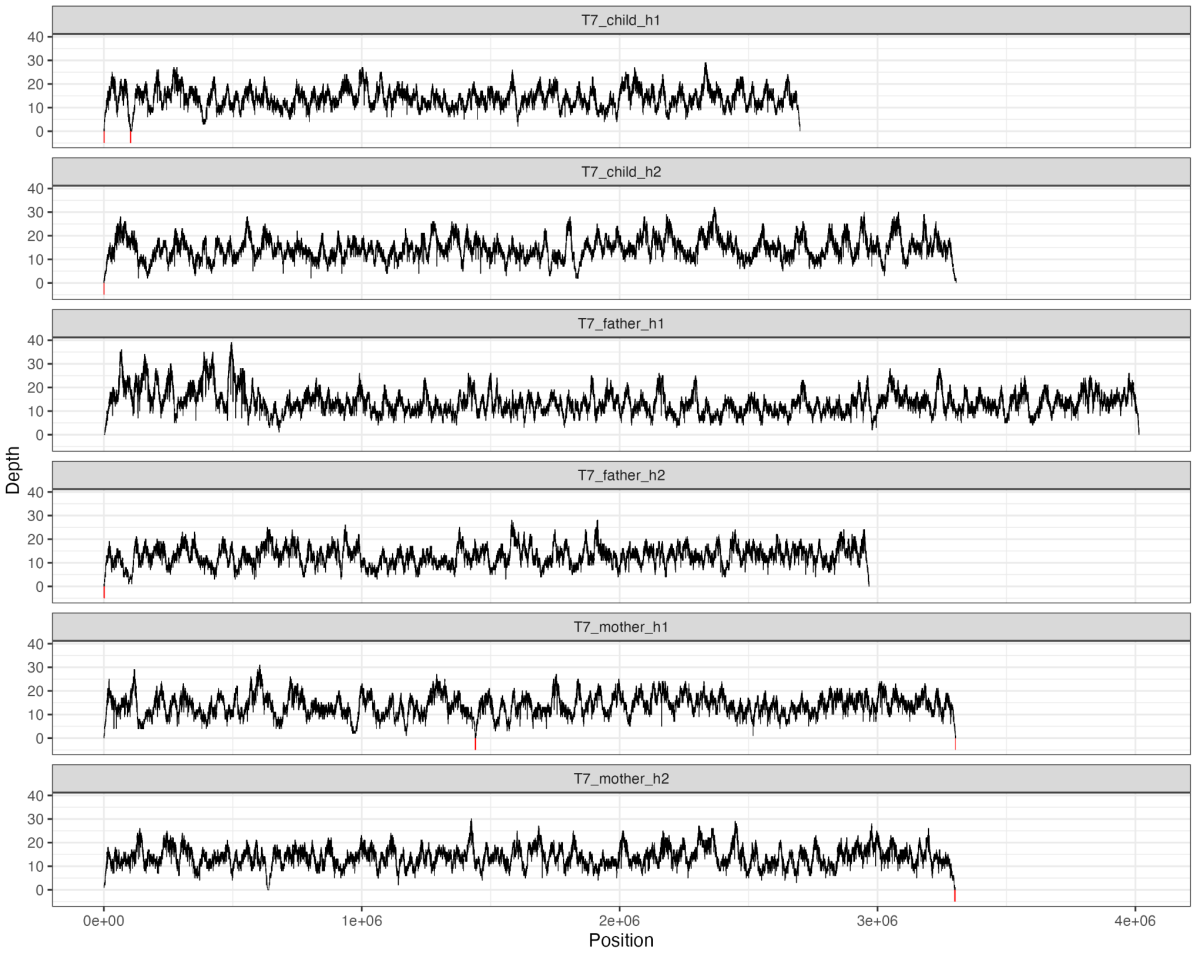


**Supplementary Figure 59**. **Depth profiles of reads mapping to assembled haplotypes for trio T7.**
Description: See Supplementary Figure 53.

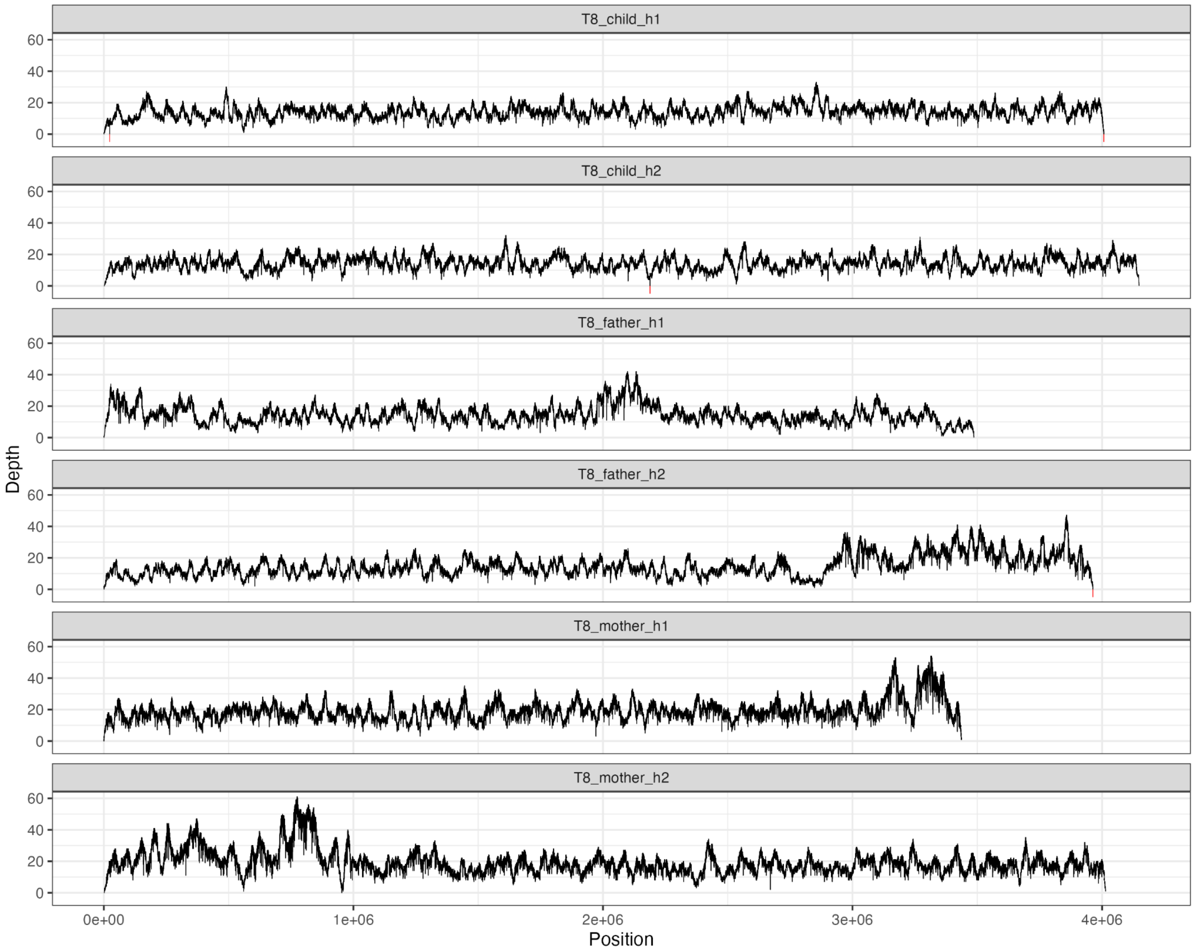


**Supplementary Figure 60**. **Depth profiles of reads mapping to assembled haplotypes for trio T8.**
Description: See Supplementary Figure 53.

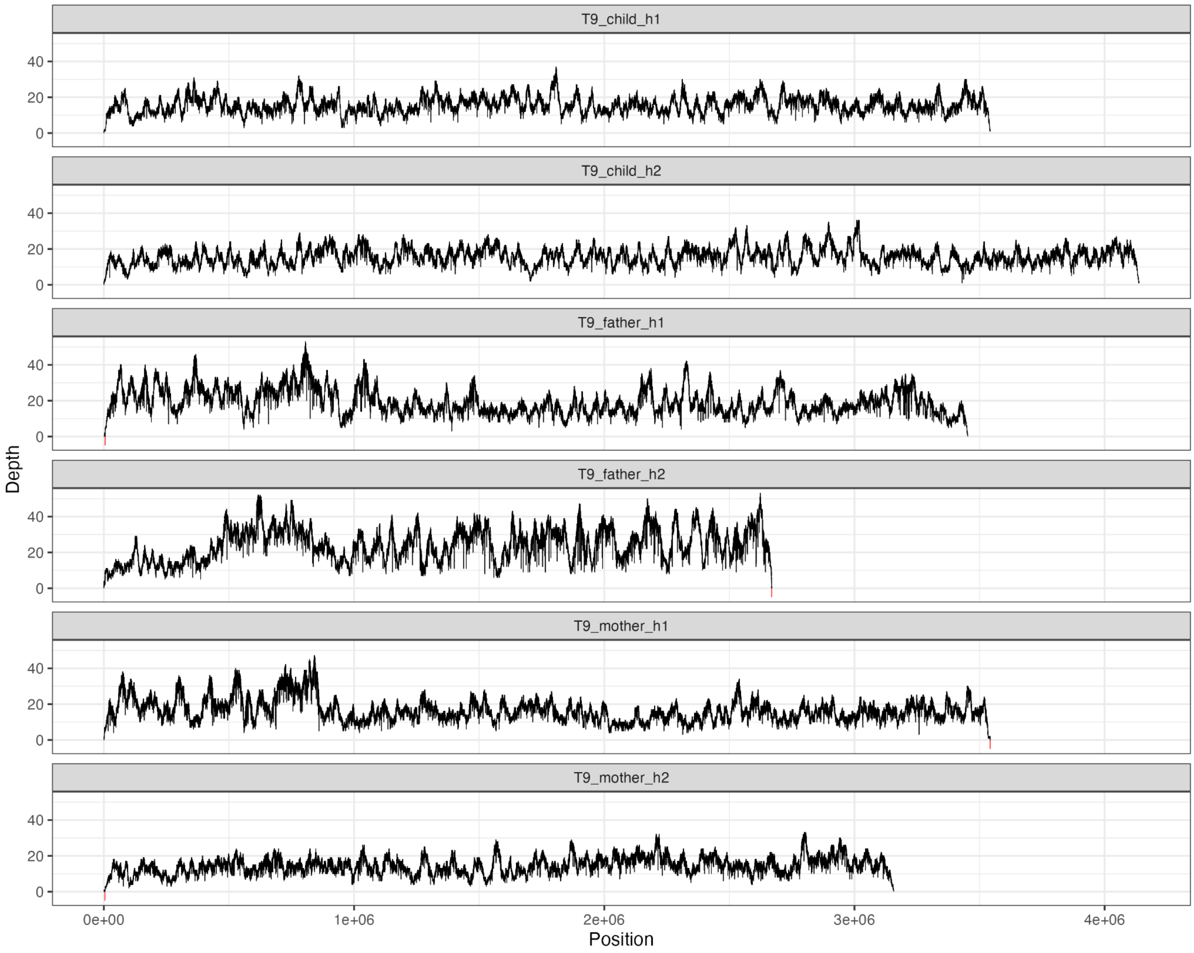


**Supplementary Figure 61. Depth profiles of reads mapping to assembled haplotypes for trio T9.** Description: See Supplementary Figure 53. T9_father_h2 and T9_mother_h1 did not assemble into single contigs, producing wavy coverage patterns for these samples.

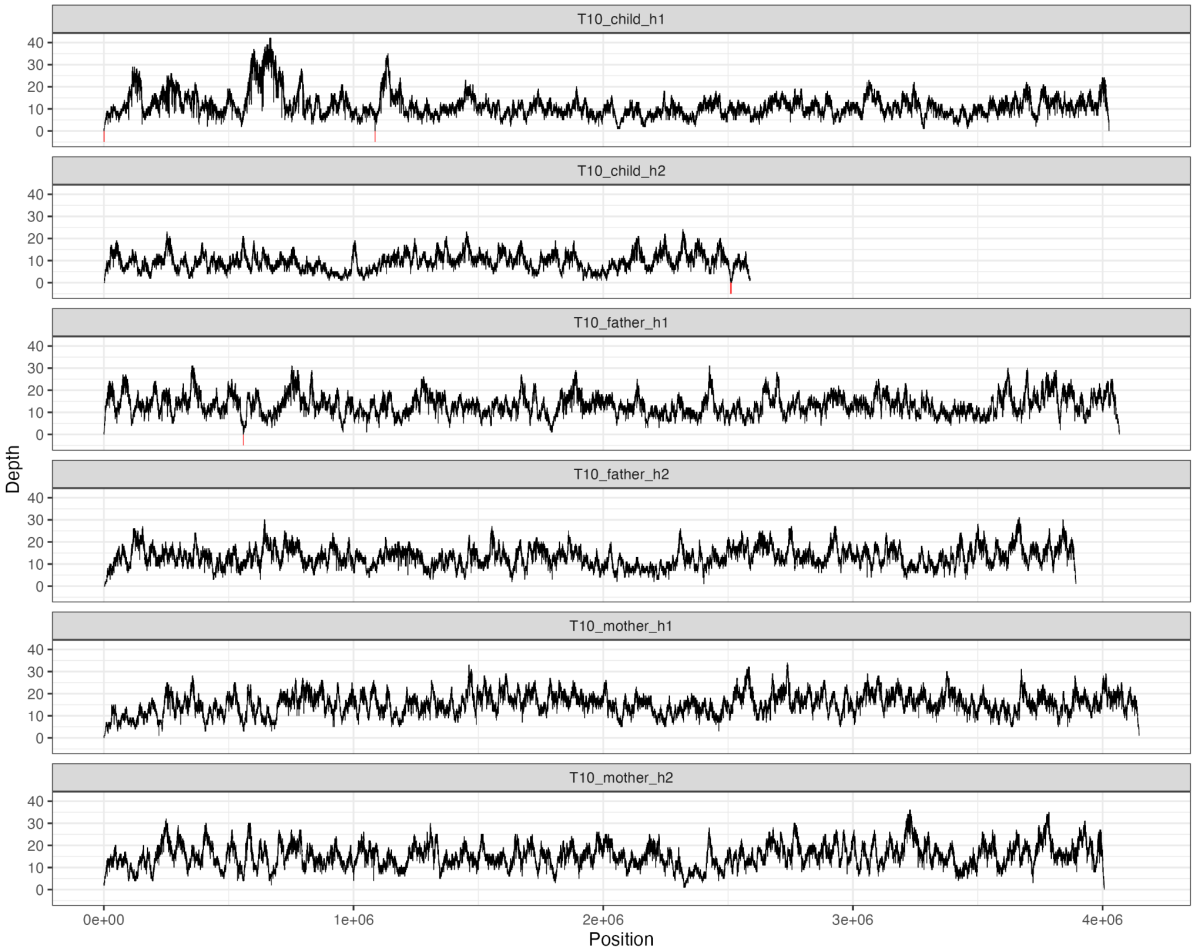


**Supplementary Figure 62**. **Depth profiles of reads mapping to assembled haplotypes for trio T10**.
Description: See Supplementary Figure 53. T10_child_h2 did not assemble into a single contig.

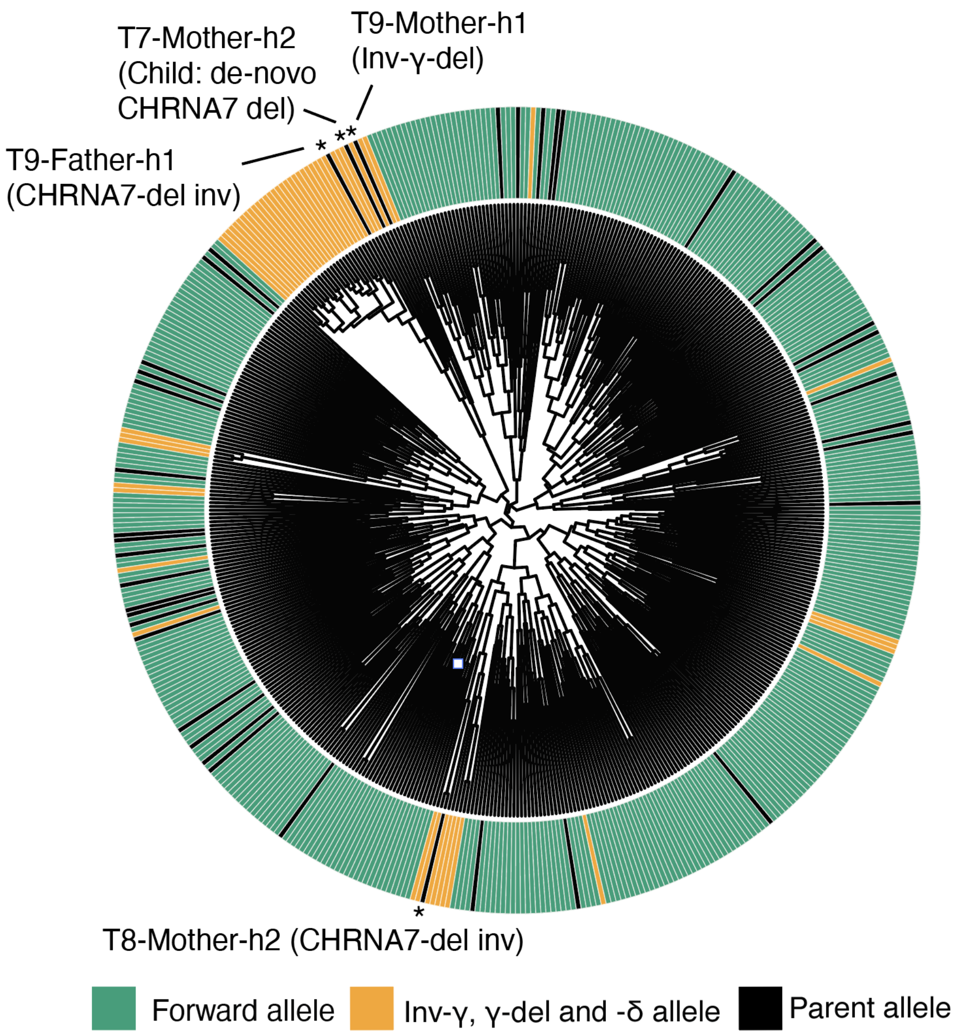


**Supplementary Figure 63. Population structure of BP4-CHRNA7-LCR′ haplotypes.** Phylogenetic tree constructed from common single-nucleotide variants (MAF ≥5%) within the unique sequence between BP4 and CHRNA7-LCR′ across 551 HPRC/HGSVC haplotypes and n=40 local parental haplotype assemblies (black). Previously characterized haplotypes predominantly with Inv-γ, γ-del or δ tend to co-cluster (orange clusters). Patient haplotypes inferred from de novo assemblies to carry Inv-γ or Inv-δ co-cluster within these inversion-associated groups, whereas non-carrier haplotypes do not, supporting the assembly-based inversion calls.

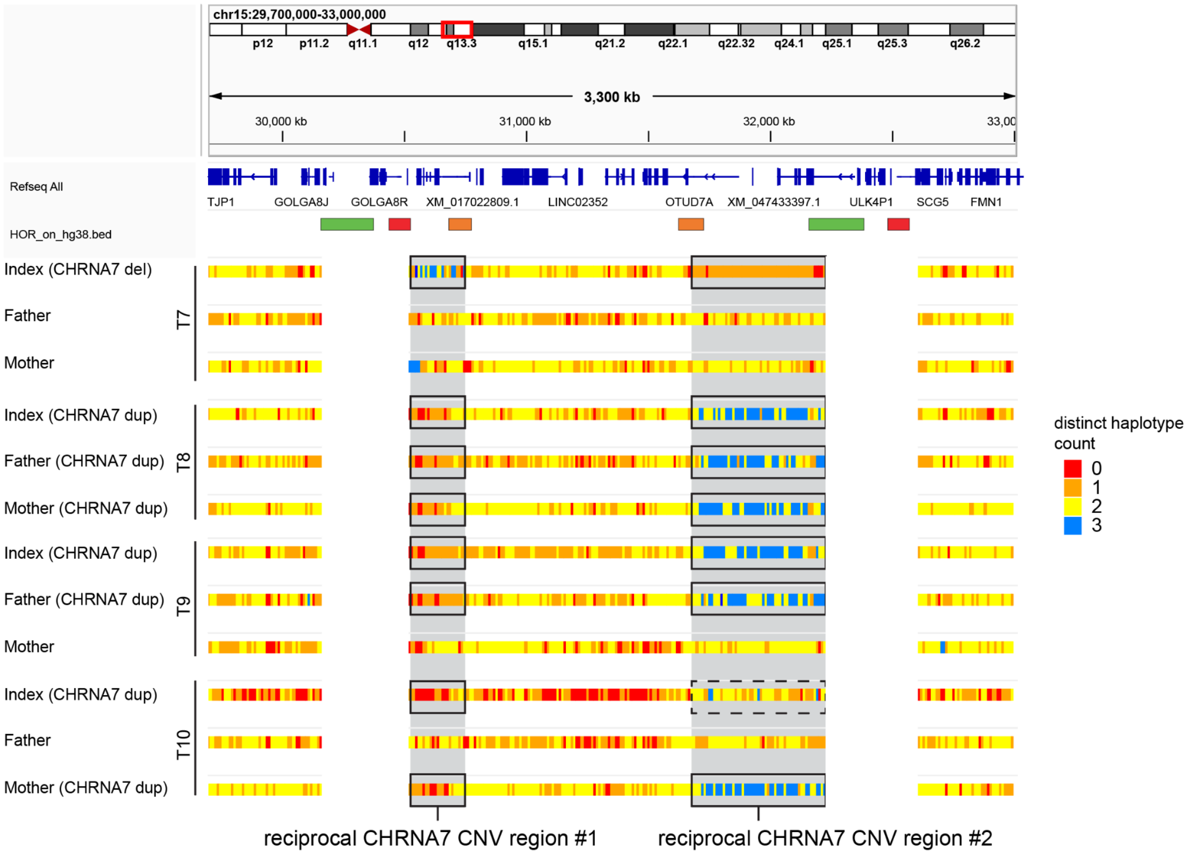


**Supplementary Figure 64. Estimation of the number of distinct alleles in *CHRNA7* CNV Trio samples.** As predicted by our *CHRNA7* CNV formation model, *CHRNA7* deletion (top row) includes acquisition of a third allele before CHRNA7-LCR. Duplications indicate the opposite signal; with three separate copies of the *OTUD7A/CHRNA7* region, and a loss of *ARHGAP11B.*

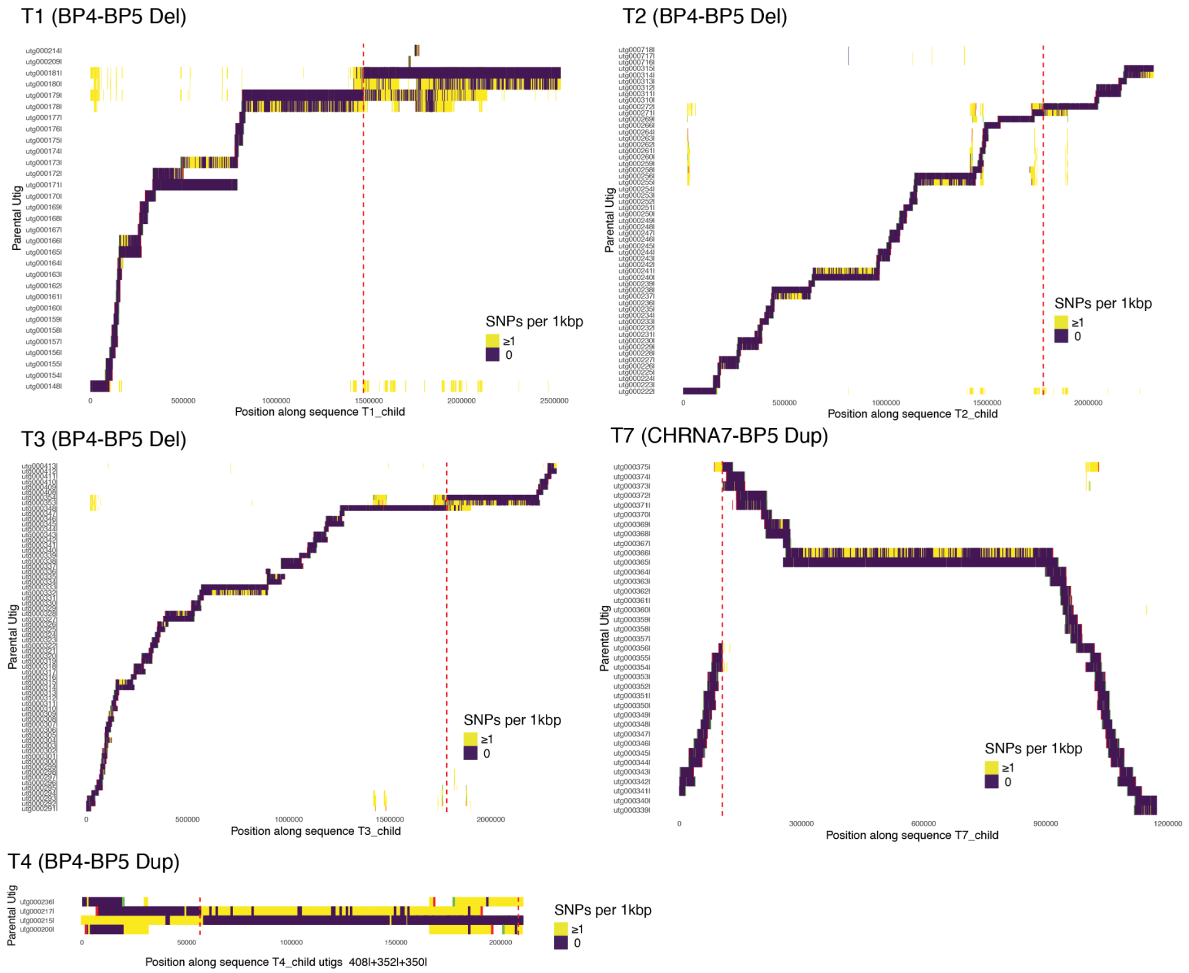


**Supplementary Figure 65. Detection of likely recombination breakpoints based on child assembly and parent unitigs using the crosshair tool.** Parental unitigs are aligned against the haplotype of the index. Aligned unitigs are colored by the number of single nucleotide differences per kb. Breakpoints are indicated by the vertical red dotted line, and visible by observing transitions of the alignments of the unitigs. i.e. two unitigs overlap and each unitig has opposite parts with good alignment (SNPs per kbp = 0) and a part with poor alignment (SNPs per kbp >=1) to the index haplotype.

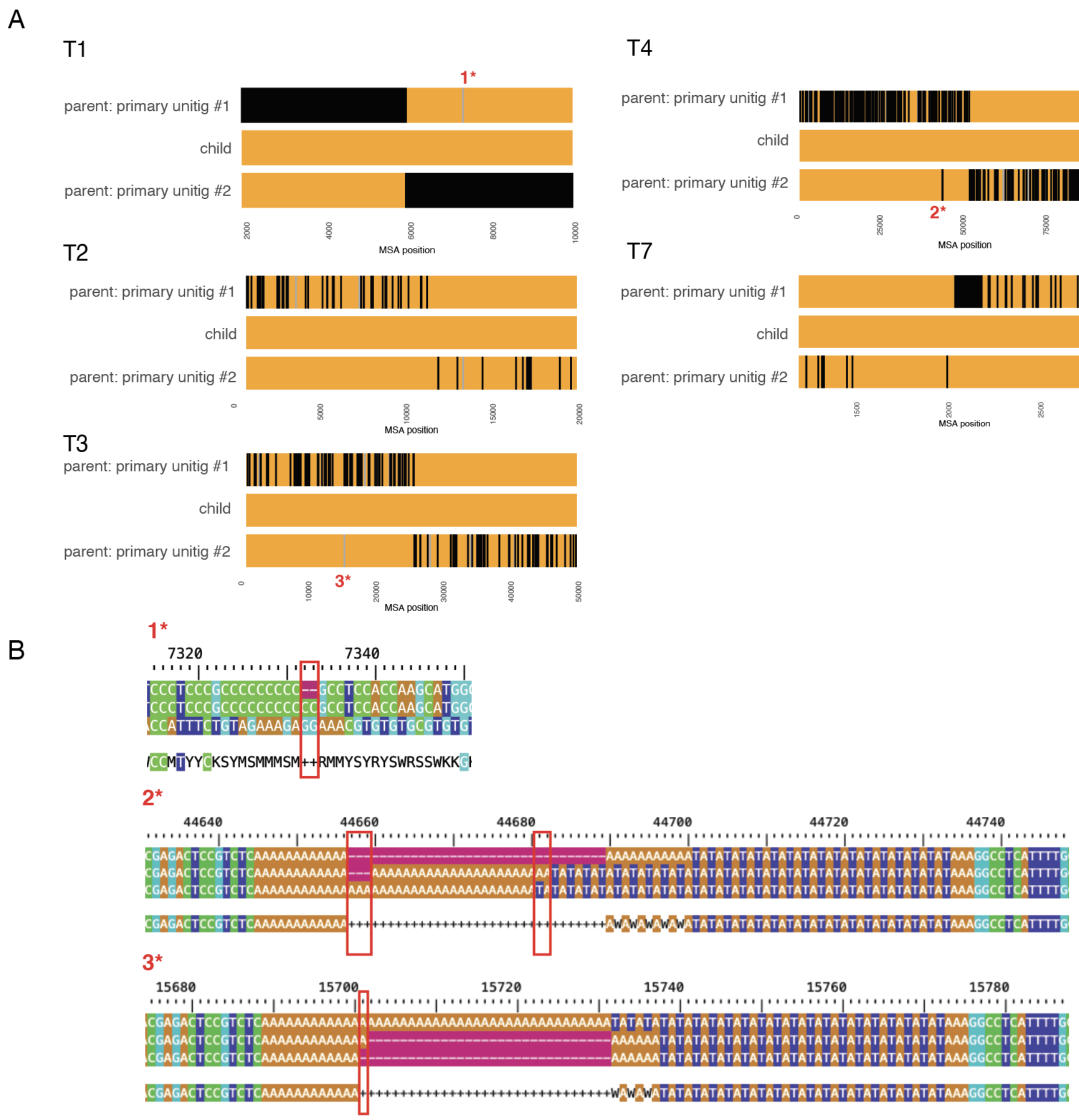


**Supplementary Figure 66. De novo breakpoints detected by multiple sequence alignment.** A) patient SNPs are aligned to the two unitigs of the parent. Orange indicates matching SNPs, black indicates a mismatch. B) Zoom-in into three non-mendelian consistent variants. All three variants are likely sequencing or assembly artifacts caused by monomeric repeats.

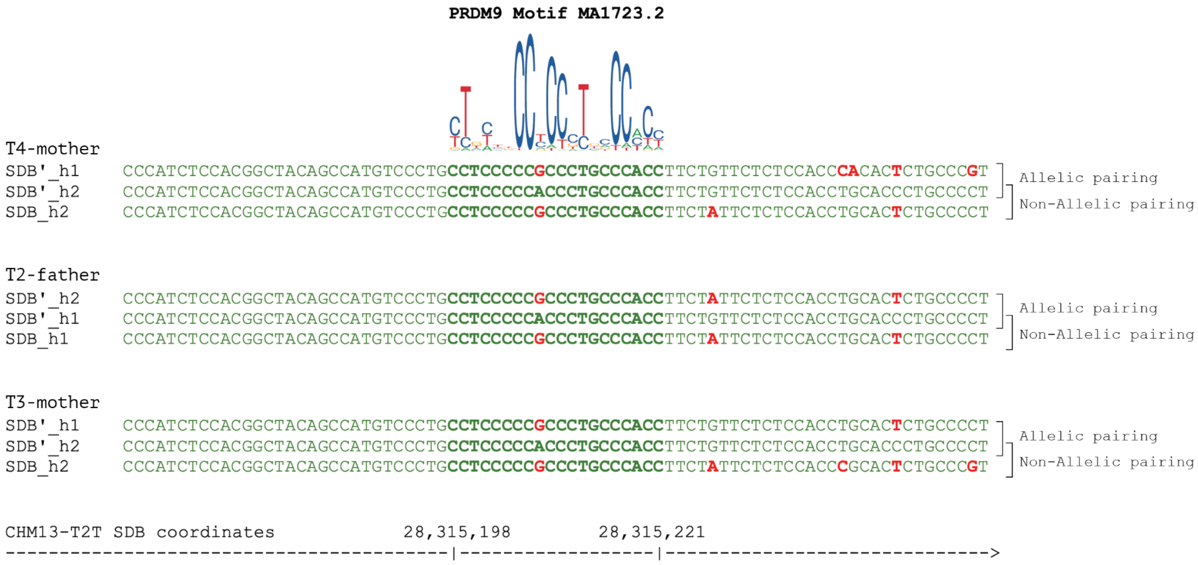


**Supplementary Figure 67. Sequence variation near the 'hot' PRDM9 motif (T2T-CHM13:chr15:28315198-28315221) in the three parental alleles on which de novo breakpoints near the motif occurred.** Only one common A>G variant with low predicted binding importance affects the PRDM9 motif directly. No obvious advantages of the non-allelic over the allelic pairings are visible at this locus.
